# Nature and location of modifier alleles determine the resolution of intralocus sexual conflict

**DOI:** 10.1101/2023.10.25.564090

**Authors:** Akanksha Singh, Koyna Jain, Manas Geeta Arun, N G Prasad

## Abstract

When selection favours different levels of expression for a shared trait in males and females, the population is said to be in Intralocus Sexual Conflict (IaSC). In a landmark study, Connallon and Clark (2010) showed that modifiers (alleles at a different locus that alter the expression of the trait under IaSC in a sex-specific manner), effectively resolve IaSC. Their model assumed that modifier alleles are “perfect”; i.e., they fully rescue the harmful effects of the deleterious allele while leaving the positive effects of the beneficial allele unaffected. In the present study, we relaxed this assumption to include more realistic “imperfect” modifier alleles. Imperfect modifiers are those that reduce the expressions of *both* detrimental allele and beneficial allele to different extents in a given sex. We also ask whether autosomes or X chromosomes are more conducive to resolution of IaSC. Our results suggest that the resolution of IaSC may not always be guaranteed when the modifier allele is “imperfect.” Interestingly, IaSC gets resolved more easily on autosomes only when modifier alleles are “perfect” or near “perfect” in concordance with the assumption made by Connallon and Clark (2010). However, in general, X chromosomes are more conducive to the resolution of IaSC.

## Introduction

Intralocus Sexual Conflict (IaSC) ensues when there are distinct sex-specific fitness optima for traits with a common underlying genetic basis in males and females (Bonduriansky and Chenoweth, 2009). IaSC can cause populations to remain stuck in a maladapted equilibrium, where neither sex is at their respective fitness maximum. This can exert substantial costs on the average fitness of the population as a whole, and can even lead to an increased risk of extinction (Lande, 1980). Numerous empirical studies, particularly over the last two decades, have shown that patterns consistent with IaSC are quite widespread (Barson et al., 2015; Berger et al., 2016; Chippindale et al., 2001; Delph et al., 2011; Eyer et al., 2019).

While empirical research on IaSC, and the term “IaSC” itself, are barely a few decades old, the underlying mathematical logic has been the subject of a large number of theoretical studies over the course of the last 70 years. Owen (1953) was the first to extend the standard population genetics framework to incorporate differential selection in the sexes. Owen (1953) showed that a single locus experiencing sex-specific selection in a diploid population can exhibit as many as three different non-trivial equilibria. A number of studies have built upon Owen’s (1953) framework to identify conditions that facilitate the invasion of alleles with sex-specific fitness effects (Parsons, 1961) and compare sex-specific viability and fertility selection (Bodmer, 1965). Sexually antagonistic (SA) selection is a special case of sex-specific selection, where selection coefficients in males and females have opposite signs. Several studies have subsequently evaluated the conditions under which SA alleles can invade a population (Haldane, 1962) and be maintained in a stable polymorphism (Haldane, 1962; Kidwell et al., 1977). Some studies have compared the efficacy of X chromosomes and autosomes at maintaining SA polymorphisms (Connallon and Clark, 2012; Curtsinger, 1980; Fry, 2010; Pamilo, 1979; Patten and Haig, 2009).

While SA selection coupled with the fact that the sexes largely share the same gene pool can constrain adaptation in males and females, there has been a longstanding consensus that SA or at least sex-specific selection is a necessary precondition for the evolution of sexual dimorphism (Andersson, 1994; Darwin, 1871; Trivers, 1972). Using a quantitative genetic approach, Lande (1980) showed that sex-specific natural and/or sexual selection can lead to the evolution of sexual dimorphism, provided there exists additive genetic variation for sexual dimorphism in the population. However, Lande’s (1980) quantitative genetic approach did not address the biological mechanism underlying the additive genetic variation for sexual dimorphism. A number of population genetic studies investigating the resolution of IaSC have attempted to fill this gap. These studies have invoked several different biological mechanisms including gene duplication (Connallon and Clark, 2011), genomic imprinting (Day and Bonduriansky, 2004), sex-specific dominance (Spencer and Priest, 2016), and modifier alleles bringing about sex-biased gene expression to model the resolution of IaSC (Connallon and Clark, 2010), ultimately leading to sexual dimorphism.

In a landmark study linking gene expression and fitness, Connallon and Clark (2010) adapted a two-locus, diploid population genetic model to a number of different scenarios describing sex-specific and SA selection including antagonistic pleiotropy and SA selection. They were able to show that most values of parameters that allow a SA polymorphism also favour the invasion of a modifier allele bringing about sex-biased gene expression. Furthermore, in a result particularly important in the context of the non-random distribution of SA loci, they also showed that the conditions for expression divergence between the sexes are less stringent on autosomes relative to the X chromosome.

Connallon and Clark’s (2010) model clarified considerable confusion stemming from the variation in the fitness schemes employed by theoretical studies in the past and provided a robust theoretical framework that attempted to explain the idiosyncrasies in sex-specific gene expression data (summarised by Dean and Mank (2014) and Jaquiéry et al. (2013)). However, the model’s prediction regarding the ease with which sex-specific genetic architectures can evolve is at odds with the data at the phenotypic level. In spite of pervasive sex-specific (and even SA) selection (Cox and Calsbeek, 2009; Singh and Punzalan, 2018), strong intersexual genetic correlations persist for a large number of traits (Poissant et al., 2010). One of the reasons for this dissonance could be a simplifying assumption made by Connallon and Clark (2010). In their model, they assumed when divergence in expression proceeded via exaggeration through males, the modifier allele only affected the expression of the male beneficial allele in females, but not the expression of the female beneficial allele in females. On the other hand, when divergence proceeded via exaggeration in females, the modifier only affected the expression of only the female beneficial allele in males. It remains to be investigated whether this is a reasonable assumption. Sex-specific modifier alleles are expected to be transcription factors that are linked to the sex-determination pathway. While they limit the expression of an allele at a certain locus in one of the sexes, it is unlikely that they have absolutely no effect on another allelic variant at the same locus. There is considerable evidence for allele-specific gene regulation. However, the assumption made by Connallon and Clark (2010) describes a highly idealized scenario where the modifier entirely shuts down the expression of only one of the alleles in one of the sexes, while leaving the other allele completely unaffected. More realistically modifiers would alter the expression of both alleles, but to different degrees (see Discussion for a plausible model for the underlying molecular biology). It is, therefore, important to investigate whether the resolution of IaSC is indeed as easily attained under these slightly more realistic conditions as it is under the conditions assumed by Connallon and Clark (2010).

In this study, we developed a variant of Connallon and Clark’s (2010) model for the evolution of sex-biased gene expression at an SA locus. We specifically allowed the modifier allele to modulate the expression of *both* alleles at the SA locus in one of the sexes, and explored the relationship between the effect of the modifier allele on the expression of the two alleles and the likelihood of the resolution of IaSC. We addressed the following questions:

1. Does the exact nature of the effect the modifier has on the expression of the SA alleles affect the resolution of IaSC? If yes, how do different types of modifiers (different values of k1 and k2, see Model below) affect conflict resolution?
2. Would the conflict be resolved more readily when the SA locus and the modifier locus are present on the autosomes or on the X chromosome?
3. What is the effect of selection coefficients and the dominance coefficient at the SA locus, as well as the recombination rates and the initial linkage disequilibrium between the SA locus and the modifier locus on the resolution of IaSC?

### Model

Our analysis builds on the two-locus population genetic model by Connallon and Clark (2010). In their model, locus A is the SA locus, with A1 and A2 being the female and male beneficial alleles respectively. Locus B is the modifier locus. When sexual dimorphism evolves via divergence through males, B1 is a neutral allele, while B2 is modelled such that it is a cis-acting modifier of the expression patterns at locus A in females alone. It is important to note that in the framework employed by Connallon and Clark (2010), B2 completely “rescues” the deleterious effects of A2 on females, but leaves the expression of A1 (i.e., the female beneficial allele) unaffected in females. We relax this assumption in our model. We allow B2 to modulate the expression levels of *both* A1 and A2 in females. We introduce two new parameters, k1 and k2, to model the degree to which the modifier allele reduces the expression levels of A1 and A2 in females, respectively. We allow k1 and k2 to vary between 0 and 1. In this sense, it is an “imperfect allele”, in contrast to the perfect modifier allele described in the Connallon and Clark (2010) model. The fitness schemes employed by us for the autosomal case and the X chromosome case are outlined in Table 1 and Table 2, respectively. Notice that larger values of k1 and k2 correspond to a greater reduction in the levels of expression of A1 and A2 in females, respectively. Therefore, intuitively, modifier alleles with large values of k2 and small values of k1, i.e., modifiers that protect females from the deleterious effects of the male beneficial allele A2, but largely leave the beneficial effects of the female beneficial allele A1 unaffected, should be better at resolving IaSC. Note that both in our model, as well as the one examined by Connallon and Clark (2010), sexual dimorphism evolves and IaSC is resolved when the haplotype A2B2 goes to fixation. This translates to a situation where the male beneficial allele (A2) is fixed, but the females are at least partially protected from the deleterious effects of A2 by the modifier allele B2.

We used the general iterative equations developed by Connallon and Clark (2010) for the case when locus A and locus B are autosomal, as well as the case when they are X-linked. These equations are summarized in the supplementary information. We substituted our fitness schemes in these equations as described in Table 1A-B for loci on autosomes and in Table 2A-B for X-linked loci. It is straightforward to show that when the modifier allele B2 is absent, the system exhibits a polymorphic equilibrium where the frequency of A2 is given by the following expressions derived by Connallon and Clark (2010):

**Table 1A.** Fitness scheme in females when the two loci are present on autosomes.

| Parental Haplotype | A1B1 | A1B2 | A2B1 | A2B2 |
| --- | --- | --- | --- | --- |
| A1B1 | $F_{11} = 1$ | $F_{12} = 1 - k_1 h f s f$ | $F_{21} = 1 - h f s f$ | $F_{22c} = 1 - (1 - k_2) h f s f$ |
| A1B2 | $F_{12}$ | $F_{13} = 1 - k_1 s f$ | $F_{22r} = 1 - [(1 - k_1) h f s f]$ | $F_{23} = 1 - [(1 - k_1 - k_2) h f s f]$ |
| A2B1 | $F_{21}$ | $F_{22r}$ | $F_{31} = 1 - s f$ | $F_{32} = 1 - [1 + h f - h f (1 - k_2) - k_2] s f$ |
| A2B2 | $F_{22c}$ | $F_{23}$ | $F_{32}$ | $F_{33} = 1 - (1 - k_2) s f$ |

**Table 1B.**
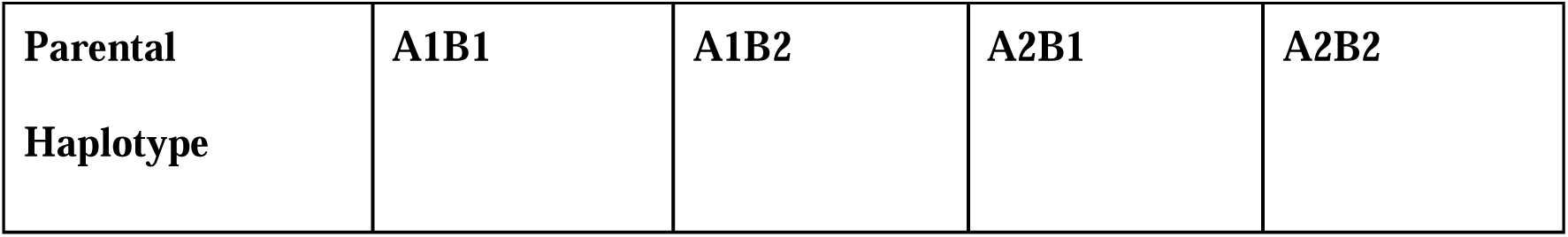
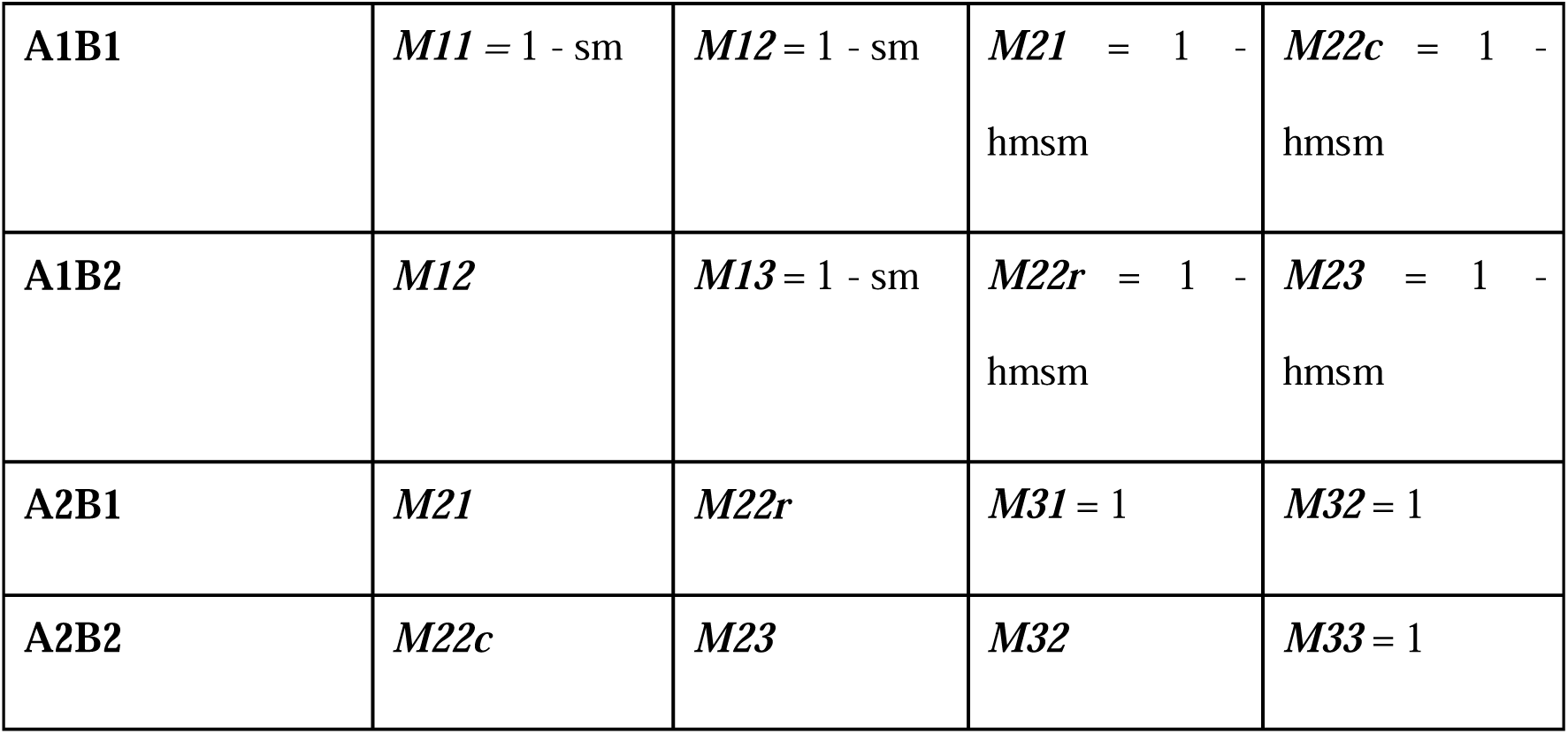
Fitness scheme in males when the two loci are present on autosomes.

**Table 2A.** Fitness scheme in females when the two loci are present on X chromosomes.

| **Parental Haplotype** | **A1B1** | **A1B2** | **A2B1** | **A2B2** |
| --- | --- | --- | --- | --- |
| **A1B1** | $F11 = 1$ | $F12 = 1 - k1hfsf$ | $F21 = 1 - hfsf$ | $F22c = 1 - (1 - k2)hfsf$ |
| **A1B2** | $F12$ | $F13 = 1 - k1sf$ | $F22r = 1 - [(1 - k1)hfsf]$ | $F23 = 1 - [(1 - k1 - k2)hfsf]$ |
| **A2B1** | $F21$ | $F22r$ | $F31 = 1 - sf$ | $F32 = 1 - [1 + hf - hf(1 - k2) - k2]sf$ |
| **A2B2** | $F22c$ | $F23$ | $F32$ | $F33 = 1 - (1 - k2)sf$ |

**Table 2B.** Fitness scheme in males when the the loci are present on X chromosomes.

| **Parental Haplotype** | **A1B1** | **A1B2** | **A2B1** | **A2B2** |
| --- | --- | --- | --- | --- |
| **Y** | **$M1 = 1 - sm$** | **$M2 = 1 - sm$** | **$M3 = 1$** | **$M4 = 1$** |

Autosomal case:

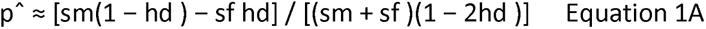

X- linked case:

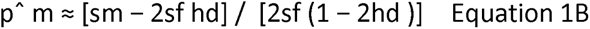

In our analyses, we consider a population polymorphic for the equilibrium given by equations 1A and 1B. We then introduce the modifier allele B2 to the population at low frequencies. The parameter D models the initial linkage disequilibrium between locus A and locus B. We then examine the behaviour of the system for various values of the parameters: the selection coefficients on males and females at locus A (sm and sf), recombination rates (rm and rf), dominance coefficient (h) at locus A. The dominance coefficient can potentially be different in males and females (hm, hf). In this study, however, we have considered the case of sex-specific dominance reversal with the same magnitude of h (i.e. hm=hf). Following Connallon and Clark (2010) we assume that fitness landscapes are concave in the vicinity of fitness optima.

Therefore, we allow h to vary between 0 and 0.5 only. The different parameter ranges in our model have been summarized in Table 3.

**Table 3.**
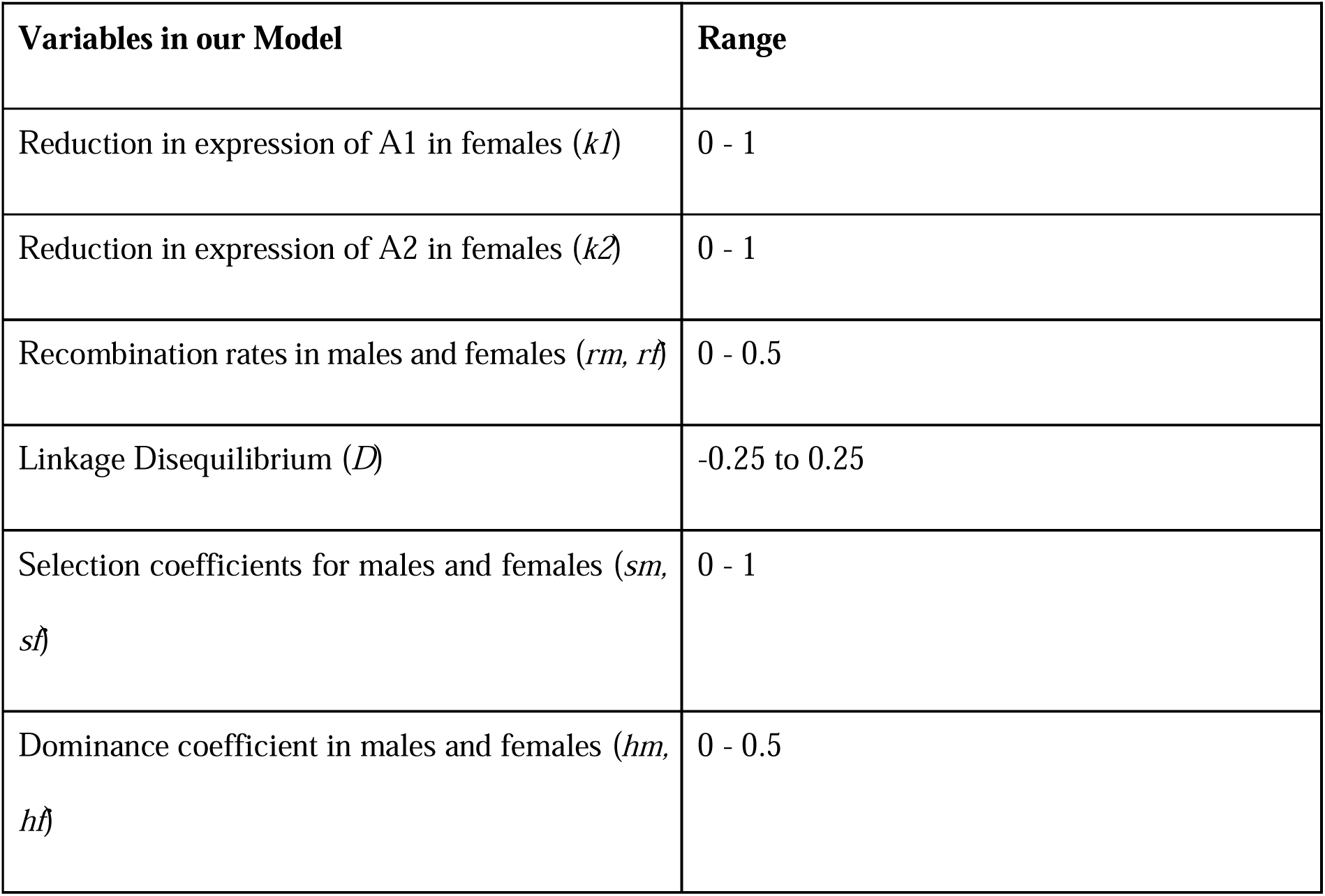
Range of different parameters in the model.

## Results

In all analyses, we investigated the fixation tendencies of the A2B2 haplotype. A2 going to fixation is ideal for males, while fixation of B2 ensures some (depending on the value of k2) rescue from the female detrimental effect of A2. Thus, the fixation of the A2B2 haplotype implies the resolution of IaSC. The swiftness with which A2B2 goes to fixation roughly corresponds to the ease of the resolution of IaSC. Therefore, we tracked the trajectory of the haplotype A2B2 through the course of 3000 generations under varying conditions. Specifically, we measured the following quantities corresponding to the ease of resolution of IaSC:

1. Using our iterative equations, we calculated the frequencies of the haplotype A2B2, allele A2, and allele B2 after 3000 generations for various values of the eight parameters: k1, k2, sm, sf, rm, rf, h, and D. For each pair of values of k1 and k2, we then measured the proportion of parameter space in sm, sf, rm, rf, hm, hf and D for which (a) the A2B2 haplotype went to fixation within 3000 generations, (b) allele A2 went to fixation within 3000 generations, and (c) the modifier allele B2 went to fixation within 3000 generations. We calculated these quantities separately for the case where the two loci are autosomal vs when they are X-linked (Figure 1).
2. For each pair of values of k1 and k2, the average of the number of generations (i.e., averaged over sm, sf, rm, rf, hm, hf, D) it took for the alleles or haplotype to get fixed, in the instances where it did get fixed (Figure 2). If the alleles or haplotype could not get fixed within 3000 generations in any case for a particular k1, k2 combination, they were assigned a time value of 3025. This was done to distinguish cases where there were no fixations from those where fixation was achieved near 3000 generations.

**Figure 1.**
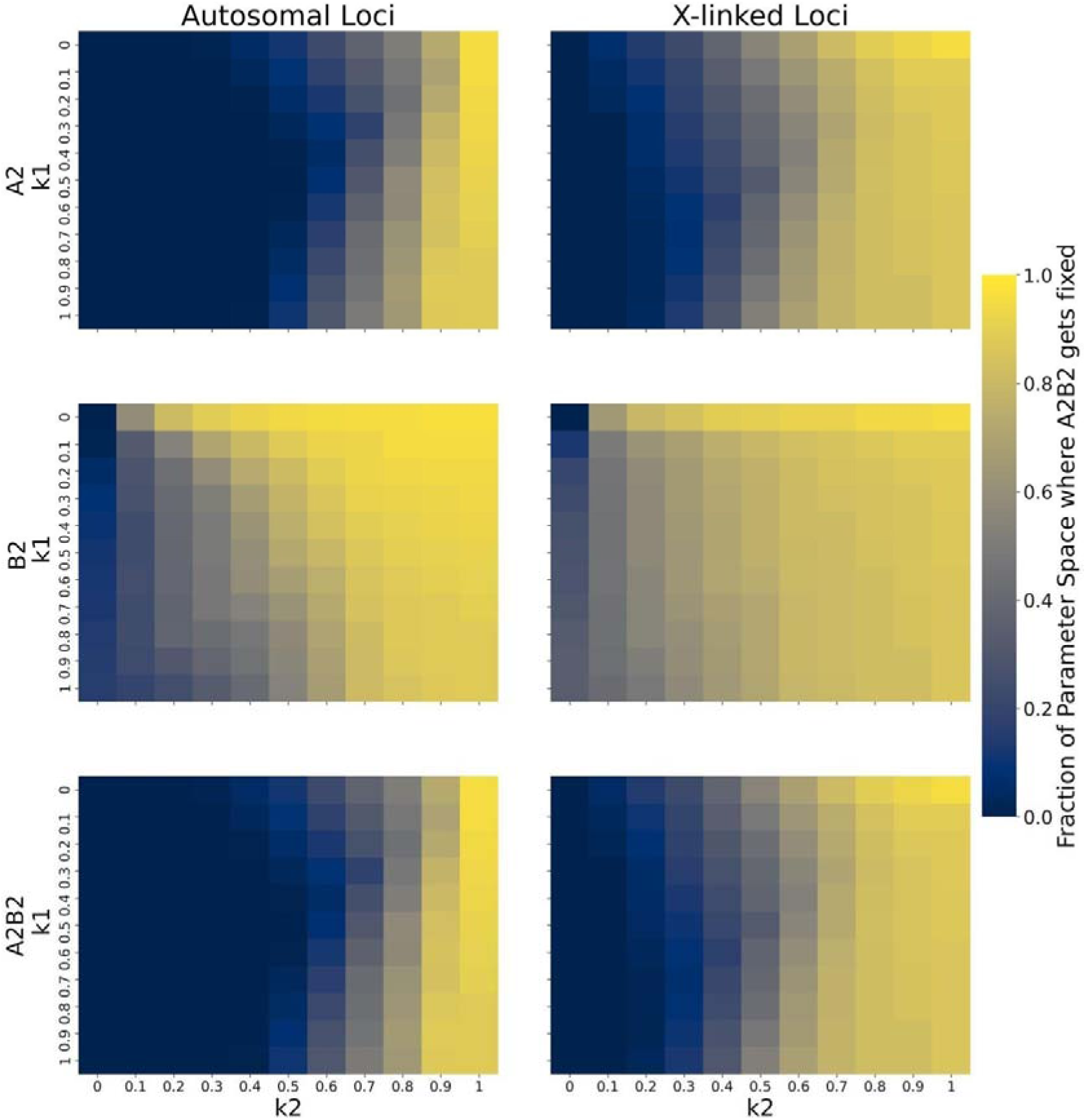
The figure shows the resolution of sexual conflict via different kinds of modifier alleles by looking at the fixation of A2, B2 alleles (first two rows), and A2B2 haplotype (third row). The different modifiers are defined by different values of k1 (suppression of female beneficial allele in females by modifier) and k2 (suppression of female detrimental allele in females by modifier). In the individual plots for the alleles and haplotypes, the x axis shows the values of k2 (step-size = 0.1) and the y axis shows the values of k1 (step-size = 1). The graph depicts the fraction of parameter space where conflict is resolved when both loci are present on the autosome (left column), or on the X-chromosome (right column). For generating the values for every k1,k2 combination, we looked at the entire range of parameters described in Table 3 and calculated the fraction of the 6 dimensional space where the different alleles and haplotype gets fixed within 3000 time steps. The lighter the shade of the block, the higher the fraction is. That is, a lighter colour represents more tendency for conflict resolution.

**Figure 2.**
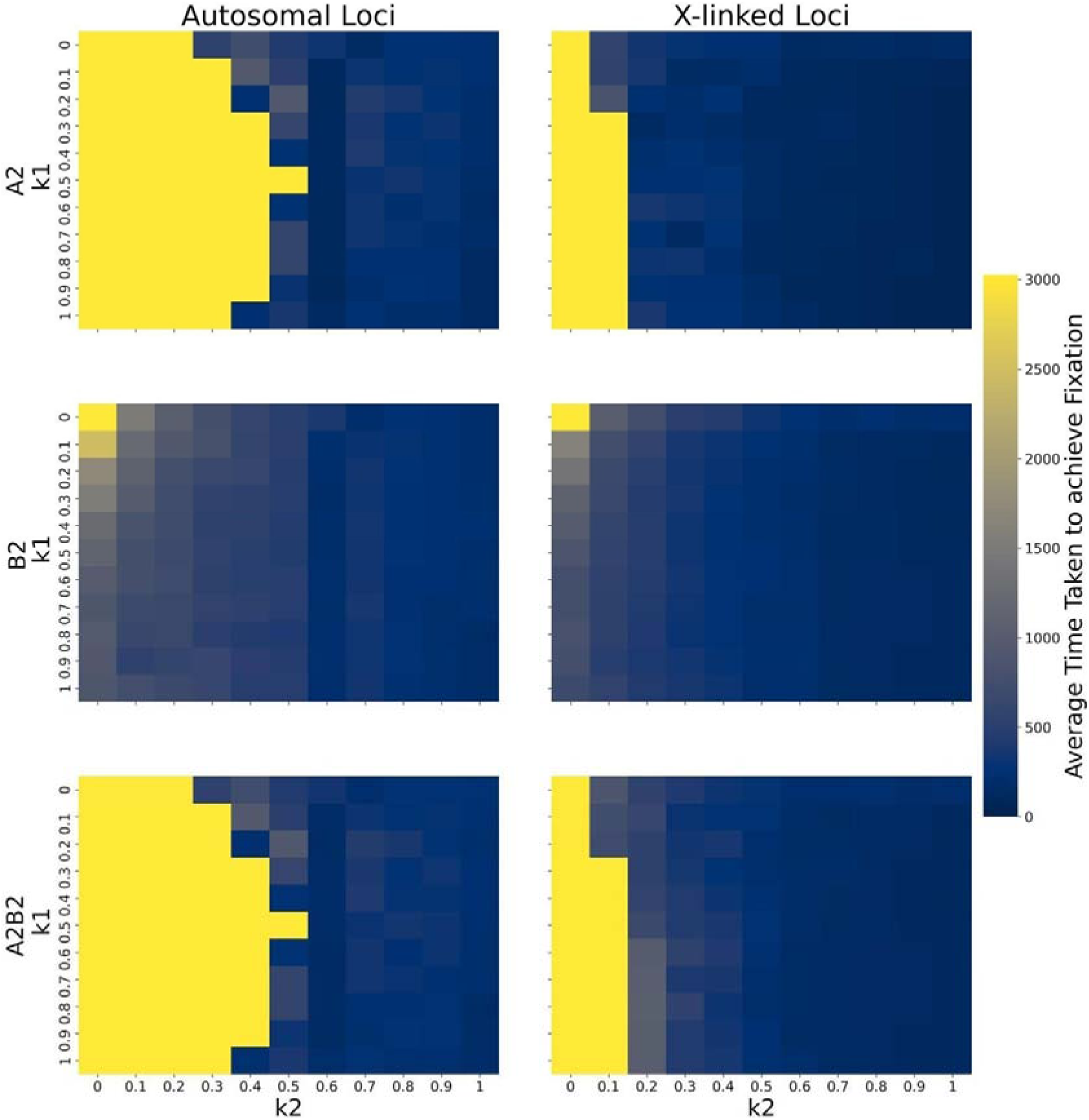
The figure shows the resolution of sexual conflict via different kinds of modifier alleles by looking at the fixation of A2, B2 alleles (first two rows), and A2B2 haplotype (third row). The different modifiers are defined by different values of k1 (suppression of female beneficial allele in females by modifier) and k2 (suppression of female detrimental allele in females by modifier). In the individual plots for the alleles and haplotypes, the x axis shows the values of k2 (step-size = 0.1) and the y axis shows the values of k1 (step-size = 1). The graph depicts the average time taken for the conflict to get resolved when both loci are present on the autosome (left column), or on the X-chromosome (right column). For generating the values for every k1,k2 combination, we looked at the entire range of parameters described in Table 3 and calculated the average time taken to fix the alleles and haplotypes in only the cases where fixation is actually achieved. For the cases where fixation was not achieved in any case, a default value of 3025 (higher than the total number of generations (3000)) was assigned. The lighter the shade of the block, the higher the average time to fixation is. That is, a lighter colour represents a tendency for conflict to not get resolved.

### 1) Effect of the nature of the modifier allele (k1 and k2) on conflict resolution in Autosomal and X-linked Loci

By varying the values of k1 and k2, we looked at how different kinds of modifiers would evolve. First, we discuss the overall effect of the modifier - taken over the entire parameter range of the other 6 parameters. In our model, an increased k2 denotes a greater fitness for A2 allele when present with the B2 allele, while an increase in k1 decreases the fitness of A1 allele in the presence of B2 allele in females. Therefore, the resolution of conflict (by fixation of A2B2) should be easiest for low k1, and high k2 (top right corner areas of Figure 1 and Figure 2).

The proportion of parameter space in sm, sf, rm, rf, h and D for which A2B2 went to fixation (and IaSC was resolved) increased with an increase in the value of k2 (Figure 1). Additionally, as the value of k2 increased, fixation of A2B2 was also attained faster (Figure 2). This suggests that as the degree of protection offered by the modifier allele B2 to females from the deleterious effects of A2 increases, the likelihood of the resolution of IaSC also increases. On the other hand, k1 appeared to have a non-monotonic effect on the ease of resolution of IaSC (Figure 1 and Figure 2). The proportion of parameter space for which A2B2 went to fixation was the lowest for intermediate values of k1 (Figure 1). Correspondingly, the time taken for the fixation of A2B2 was the highest for intermediate values of k1 (Figure 2). These patterns were also reflected in the proportion of parameter space for which the allele A2 went to fixation (Figure 1) and the time taken for the fixation of A2 (Figure 2). However, the effect of k1 and k2 on the fixation tendency of B2 was different. As k1 increased, the fixation tendency of B2 decreased (Figure 1 and Figure 2).

Overall, we find that k2 plays a primary role in determining the ratio of parameter space where the alleles get fixed. These analyses confirm that fixation is achieved in almost the entire parameter space (95.7%) for invasion of male beneficial mutation, under conditions assumed by Connallon and Clark (2010) (equivalent to k2=1, k1=0 in our model; i.e., the top right corner in Figure 1). We also find that for k2=1, i.e., when B2 completely rescues the deleterious effects of A2 in females, the haplotype A2B2 goes to fixation for more than 85% of the parameter space irrespective of the value of k1.

### 2) Comparison of efficiency of IaSC resolution in autosomal vs X-linked Loci

Next, we asked whether the resolution of IaSC via the fixation of the A2B2 haplotype was easier on autosomes or the X chromosome. We compared the fixation tendencies of A2, B2 and A2B2 haplotype averaged across the entire parameter space in sm, sf, rm, rf, h and D for each combination of k1 and k2 when the loci are on autosomes versus when they are on the X chromosome.

When comparing X-linked loci to autosomal loci, we find that a greater proportion of fixation is observed in X-linked loci for both alleles A2 and B2 and therefore, for the haplotype A2B2, particularly for low values of k2 (Figure 1). Correspondingly, we find that the average time taken for fixation of A2B2 haplotype is lower when the loci are on the X chromosome than when on autosomes. This pattern was also observed when we looked at the fixation tendencies of the allele A2 on autosomes vs the X chromosome. Interestingly, the fixation tendencies of B2, both in terms of the proportion of parameter space leading to fixation and the average time to fixation, were fairly similar across autosomes and X chromosomes (Figure 1 and Figure 2).

Overall, it is clear that for low k2 modifier alleles, the conflict is more likely to be resolved when the loci are located on the X-chromosome.

An interesting difference in the autosomal and X-linked case is the sudden low time of fixation for k2=0.6 when loci are present on autosomes. When the loci are present on autosomes, A2B2 appears to go to fixation *earlier* than B2 at k2=0.6 - which looks counterintuitive at first sight. However, keeping in mind that B2 goes to fixation in more cases than A2 (Figure 1), it is possible that the time taken by B2 to go to fixation in the cases where A2 also goes to fixation is lower than when B2 alone goes to fixation, bringing down the average time taken for A2B2 to get fixed. When the loci are X-linked, A2B2 seems to go to fixation earlier.

### 3) Effect of initial linkage disequilibrium, recombination rates, selection coefficients, and dominance coefficients on resolution of conflict

We explored an 8-dimensional parameter space, with the prime parameters being k1 and k2. We divided the six other variables into two groups - D, rm, rf and h, sm,sf for the ease of analysis. When analysing the effect of variables of one group, the variables of the other group were fixed at 0.2 (except for D, which was fixed to 0 when not being analysed). Analysis of each case comprised two parts: 1) the frequency of A2B2 at generation 3000 and 2) the time taken for A2B2 to reach fixation. All of these analyses were carried out for both when the loci were on autosome and when they were on X chromosome.

#### (A) Selection Coefficients and Dominance Coefficient

In this case, the initial linkage disequilibrium coefficient (D) is assumed to be 0 and recombination rates for males and females (rm, rf) are kept constant to 0.2. These results are represented in the supplementary information Figures S1.1-S1.5, S2.1-S2.5 (for loci on autosomes), S3.1-S3.5, S4.1-S4.5 (for X-linked loci), and in Supplementary Videos 1, 2, 3, 4.

Within each subplot, we observe a decline in the frequencies of A2B2 at generation 3000 with increase in sf and an increase in fixation with increasing sm. Increase in the value of sm and sf leads to reduced time to fixation. We also observe a somewhat diagonal relationship in the subplots. Larger values of sm relative to sf corresponded to the equilibrium frequencies of A2B2 being close to 1. On the other hand, decreasing sm relative to sf corresponded to very low equilibrium frequencies of A2B2.

Increasing the value of dominance coefficient, we find that the diagonal relationship intensifies, that is, there is a trend for A2B2 to go to complete fixation above the diagonal, while it gets completely eradicated below the diagonal. On the diagonal, the frequency of A2B2 is found to increase with h and as the value of h increases, the time taken for A2B2 to get fixed decreases. When k2 is sufficiently high, at high values of h, the subplots start to resemble each other. Thus, k1 and k2 do not make significant contributions to final frequencies in this region of the parameter space.

A2B2 does not go to fixation if there is no selection on the A locus for either of the sexes. It also does not go to fixation if k2 (the rescue effect that the modifier allele has on the presence of A2 in females) equals 0. For some values of sm and sf, A2B2 goes to fixation below the diagonal at low h, but does not go to fixation at high h. As the value of h increases, the time taken for A2B2 to get fixed decreases. Increase in h also brings about fixation of A2B2 at lower values of k2 (Figure S1, S2, S3, S4 and videos 1,2,3,4).

#### (B) Effect of Recombinant Rate and Initial Linkage Disequilibrium

Here, we look into the effect of recombination rates in both sexes (rm, rf) and the initial linkage disequilibrium (D) on resolution of conflict. In this case selection coefficients and dominance coefficient are kept constant at 0.2. The results are summarized in supplementary Figures S5.1-S5.5, S6.1-S6.5 (for loci on autosomes), S7, and S8 (for X-linked loci) and videos 5 and 6. There is no effect of varying recombination rates in males and females even at different initial linkage disequilibrium values. Only when rm=rf=0, that is, there is total linkage between the two loci, a lesser frequency of A2B2 is observed for lower values of k2 and more time for fixation of haplotype A2B2. These observations lead us to conclude that there is no effect of recombination rates for the resolution of conflict. k1 and k2 are the main determining factors once h, sm and sf are kept constant. In saying so, we must note that the initial linkage disequilibrium could be varied only between −0.02 to 0.02, because the initial frequencies were too high or low to accommodate a larger range of D (Figure S5, S6, S7, S8 and videos 5,6).

## Discussion

IaSC is a consequence of males and females sharing the same gene pool while experiencing markedly different selection pressures (Schenkel et al., 2018). IaSC is usually defined for traits that have a common underlying genetic basis in males and females, but have vastly different sex-specific fitness optima (Bonduriansky and Chenoweth, 2009). IaSC is typically thought to get resolved by the evolution of sex-specific genetic architectures, leading to the evolution of sexual dimorphism. One of the ways in which this can happen is through the invasion of alleles with sex-specific effects on phenotypes (Rhen, 2000). While variation for alleles with entirely sex-limited effects are rare, they have been detected in a few organisms. For example, in domestic sheep, three alleles have been shown to control horn development. Two of the alleles (one causing a lack of horns in both sexes and another promoting horn development in both sexes) are expressed in both sexes. However, the third allele, being androgen dependent, causes horn development only in males (Montgomery et al., 1996). Thus, horns may become entirely male-limited if this third allele invades the population and goes to fixation. However, given that strong intersexual genetic correlations for traits abound in nature (Poissant 2010), a majority of the mutations are likely to be expressed in both sexes, albeit to differing degrees. Therefore, it is unlikely that most sexual dimorphism observed in the nature is a consequence of the invasion of completely sex-limited alleles. Alternatively, it is possible that the allele coding for trait exaggeration (say, larger horns) in both sexes increases in frequency in conjunction with a “modifier” allele at an entirely different locus, that reduces or entirely inhibits the expression of the exaggerated trait (e.g., horn) in females. Such “modifier mechanism” is one of the most commonly invoked mechanisms for the resolution of IaSC leading to sexual dimorphism. In a landmark study, Connallon and Clark (2010) showed, among other important results, that a modifier reducing the expression of the deleterious allele in one of the sexes could invade the population. In this study, we modified a version of Connallon and Clark’s (2010) model by specifically allowing the modifier to modulate the expression of both male beneficial as well as female beneficial alleles in females at the SA locus and examined the efficacy of various kinds of modifier alleles at resolving IaSC. Our main findings are as follows:

(1) When the modifier allele is an “ideal” allele (i.e., k1 = 0, k2 = 1) resolution of IaSC is largely guaranteed. However, as the protection offered by B2 to females against the deleterious effects of A2 diminishes (i.e., as k2 decreases), the resolution of IaSC becomes harder. The effect of k1 on the resolution of IaSC was non-monotonic, particularly in autosomes. Lower and higher values of k1 corresponded to faster resolution of IaSC than intermediate values of k1.
(2) Overall, X chromosomes were found to be more conducive than autosomes to the resolution of IaSC leading to the evolution of sexual dimorphism.
(3) Assuming conditions of sex-specific dominance reversals, increasing the dominance coefficient (h) led to a reduced probability of IaSC being resolved.
(4) When males were under selection for increased expression, stronger selection in males relative to females favoured resolution of IaSC.
(5) Recombination rate did not have a significant effect on the efficiency with which IaSC was resolved.

Below, we discuss the implications of these findings.

### A. Resolution of IaSC via modifiers bringing about sex-specific gene expression may not be guaranteed

Sex-specific, and even SA, selection is incredibly common in nature (Cox and Calsbeek, 2009; Singh and Punzalan, 2018). A large number of theoretical studies predict that such selection can favour the evolution of sex-specific genetic architecture resolving IaSC (Day and Bonduriansky, 2004; Connallon and Clark, 2011, 2010; Spencer and Priest, 2016). Consistent with theoretical predictions, patterns of sex biased gene expression have been reported in many organisms (Ellegren and Parsch, 2007; Grath and Parsch, 2016). However, most studies that have investigated sex-biased gene expression have assumed that extant sex bias in gene expression is a relic of past SA selection, without providing any independent evidence (but see Wright et al. (2018)). Some studies have argued that sex-biased gene expression can arise via evolutionary mechanism other than IaSC, and it may not be a reliable indicator of SA selection (Kasimatis et al., 2017; Rowe et al., 2018; Ruzicka et al., 2020). Often, sex bias in gene expression is limited to the gonads, and genes expressed in somatic tissues are rarely expressed in a sex-biased manner (Stewart et al., 2010). It is, therefore, not very surprising that these patterns at the molecular level seldom translate to patterns at the phenotypic level (Dean and Mank, 2014). In fact, in spite of strong sex-specific selection (Singh and Punzalan, 2018), strong intersexual genetic correlations have persisted (Poissant et al., 2010), preventing sex-specific adaptation. Stewart and Rice (2018) performed artificial sexually antagonistic selection on body size in *Drosophila melanogaster*, which has a strong intersexual genetic correlation for body size. They showed that in spite of strong disruptive selection between the sexes, body size did not evolve in a sex-specific manner for a substantial period of time. Taken together, these findings seem to suggest that intersexual genetic correlations may not be as easily resolved via the evolution of sex-specific genetic architecture, as previously thought.

It is in this context that our finding, suggesting that the resolution of IaSC may not be always guaranteed via modifiers brining about sex-biased gene expression, is relevant. Crucially, we showed that upon relaxing Connallon and Clark’s (2010) assumption that the modifiers completely inhibit the expression (in one of the sexes) of the deleterious allele while leaving the beneficial allele unaffected, invasion of modifiers becomes more constrained. This corresponds to the condition k2=1, k1=0 as per our model. Upon looking at an expanded set of modifier alleles, we found that the k2=1, k1=0 is a special case of the system, where conflict almost always gets resolved. This result is not the representative behavior of the entire system. That is, if the modifier allele invades a population, there is a good chance that it may not go to fixation and therefore, resolve sexual conflict. In fact, even when the negative impact of the modifier allele (k1) equals 0, conflict resolution is not guaranteed. Hence, the value of k2 must be above a certain threshold (that varies non-monotonically with k1) in order to ensure fixation of A2B2 and thereby, resolve sexual conflict.

To understand why Connallon and Clark’s (2010) assumption may reflect a highly idealised scenario, it might be instructive to think of what it translates to in terms of the molecular biology of the system. While there is ample evidence of allele-specific regulation of gene expression (Buckland, 2004; Knight, 2004; Pastinen, 2010), it is not clear whether this is a product of the complete silencing of one allele accompanied by no change to the expression of the other allele. Additionally, in Connallon and Clark’s (2010) case the regulator (i.e., the modifier allele) needs to perform such highly extreme sex-specific regulation only in one of the sexes. Imagine that the modifier allele B2 is a transcription factor that controls the expression at locus A only in females by binding to the promoter and preventing transcription. In theory, it is possible that B2 binds to the promoter of A2 with a very good efficiency, but does not bind to the promoter of A1 *at all.* However, a more realistic scenario is one where B2 binds to the promoters of both A1 and A2, but with different binding efficiencies, such that the expression levels of both A1 and A2 are both reduced, but to different degrees. In our model, in some sense, k1 and k2 model the binding efficiencies of B2 to the promoters of A1 and A2, respectively. Our results imply that as long as B2 binds imperfectly to the promoter of A2 (along with also binding to the promoter of A1), the fixation of the A2B2 haplotype, and therefore the resolution of IaSC, becomes much more difficult. This argument can also be extended to scenarios when expression regulation happens by chromatin remodelling or by micro RNAs.

While conceptualizing our variant of the Connallon and Clark (2010) model, we predicted that ease of conflict resolution would vary monotonically with both k1 and k2. As k1 is the reduction of the expression of the beneficial allele in females, we expected resolution of conflict to become more difficult with increase in k1, because a higher k1 implied a ‘worse’ modifier. Since k2 stands for the reduction in expression of the detrimental allele in females (and therefore increase in female fitness), we expected resolution to increase with increase in k2, since this was a ‘better’ modifier. Our results do seem to comply with our predictions for monotonic increase in resolution with k2. However, it is not so simple with k1. We find that resolution is more difficult for moderate values of k1, than higher values of k1. Thus, there seems to be a non-monotonic effect of k1 on ease of resolution. One possible explanation for this observation could be that at higher values of k1, in the presence of modifier allele, the female beneficial allele (A1) renders less fitness to females than the female detrimental allele (A2) (Figure 1 and 2). This happens because of the higher values of the detrimental effect of the modifier on the female beneficial allele, as characteristic of high k1. Thus, in the presence of the modifier, A2 becomes the favored allele even in females, subsequently leading to resolution of sexual conflict (as A2 is already favoured in males).

As we increase the selection coefficient in males (sm), selection pressure on male increases. Since A2 is the male beneficial allele, its frequency in the population increases as a result of strong male selection. However, A2 is detrimental to females. In such conditions, the presence of B2 has the potential to rescue the phenotype in females. Therefore, the tendency of A2B2 to go to fixation increases. When there is an increased selection on females (i.e., an increased sf), A2 will be disfavored in the population, and therefore we see lower frequencies of A2B2 in the population. This brings about a diagonal relationship in the subplots of Figures S1.1-S1.5, S2.1-S2.5, S3.1-S3.5, and S4.1-S4.5. When we looked at the effect of dominance coefficient, we found that it plays a significant role in the evolutionary trajectory. One explanation for the trends observed could be that - when h is low, it implies that the phenotype of the heterozygotes have fitness similar to that of the beneficial homozygote. Thus, there is a broad range of parameter values where the heterozygote can survive even without the rescue effect of the modifier. However, when h is large, there is a huge difference between the homozygous beneficial phenotype and the heterozygous phenotype, increasing the need for resolution of conflict through modifiers.

We also found that when the two loci are perfectly linked, the tendency for conflict resolution is even lower as depicted in Figures S5.1-S5.5, S6.1-S6.5, S7, and S8. When the loci are not completely linked, then it does not seem to matter what the recombination rates or initial linkage disequilibrium are (at least in the analyses we performed). This could potentially be an artifact of the values of selection and dominance coefficients used for these analyses.

### B. X chromosomes are more conducive to the resolution of IaSC relative to autosomes

A number of theoretical studies have investigated the fate of SA polymorphisms on X chromosomes compared to autosomes. Rice (1984) showed that X chromosomes can maintain SA polymorphisms for a much broader set of parameter values than autosomes, provided that female-beneficial alleles are at least partially dominant, and the dominance coefficients are equal between males and females. However, other studies that explored other dominance conditions did not find support for the idea that X chromosomes should be “hotspots” for SA fitness variation (Connallon and Clark, 2012; Curtsinger, 1980; Fry, 2010; Pamilo, 1979; Patten and Haig, 2009). It is important to note that our model, like Fry (2010) and Connallon and Clark (2012), assumes conditions that correspond to sex-specific dominance reversal, i.e. the female-beneficial allele is dominant in females and the male-beneficial allele is dominant in males. In their two-locus model, Connallon and Clark (2010) assumed similar dominance conditions and showed that the resolution of IaSC is much easier on autosomes relative to X chromosomes. However, Connallon and Clark (2010) investigated only “ideal” modifier alleles, which correspond to the k2=1, k1=0 condition as per our model. Our model relaxes this condition to allow imperfect modifier alleles that do not entirely rescue the deleterious effects of A2 in females, and also affect the expression of A1 in females. Like Connallon and Clark (2010), at extremely high values of k2, for every k1, our model predicts that autosomes resolve sexual conflict in a larger fraction of the parameter space. However, our results are in sharp contrast to the findings of Connallon and Clark (2010) for the rest of the k1-k2 plane. In general, we find that X chromosomes lead to conflict resolution more effectively in terms of both - fraction of parameter space, as well as, average time taken (Figure 1 and 2). They take less time to resolve conflict than autosomes, even in the cases mentioned above, where k2 is high.

A number of empirical studies have investigated the relative abundance of genes that are expressed in a sex biased manner on autosomes vs X chromosomes. Interestingly, there is no consensus among gene expression studies with respect to whether there should be an over-representation of sex biased genes on X chromosomes. Some studies have indeed detected a disproportionately high abundance of sex biased genes on X chromosomes, while other failed to detect such a trend (see Dean and Mank (2014) and Jaquiéry et al. (2013) for a summary). Nevertheless, there is phenotypic evidence for sexual dimorphism or sex-dependence associated with the X chromosome for many traits including lifespan (Griffin et al., 2016), body size (Matthews et al., 2019) and locomotory activity (Long and Rice, 2007).

## Conclusion

In this study, we investigated the resolution of Intralocus sexual conflict and its dependence on the nature of modifier allele. Our model allows for “imperfect” modifier alleles i.e., the modifiers do not rescue the effects of the deleterious allele fully and/or also reduce the expression of the beneficial allele. It is not unreasonable to expect that a large proportion of naturally-occurring modifier alleles are imperfect. We find that the resolution of IaSC via modifiers that bring about sex-biased gene expression may not be guaranteed, particularly if the modifiers are “imperfect”. X chromosomes are more conducive to the resolution of IaSC and the evolution of sexual dimorphism than autosomes especially when the modifier allele is “imperfect”. Therefore, the nature of the modifier allele is crucial in determining the evolutionary trajectory traversed by the population when faced with Intralocus Sexual Conflict.

## Supporting information

X-h-Time

X-h-Frequency

A-h-Time

A-h-Frequency

A-D-Time

A-D-Frequency

## Notes

### Competing Interest Statement

The authors have declared no competing interest.

