## Supplementary Material for "Nature and location of modifier alleles determine the resolution of intralocus sexual conflict"

Supplementary Information

### Recursion equations

We used the same recursion equations as Connallon and Clark (2010) to simulate the system.

Haplotypes and sex-specific frequencies

| Haplotype | Frequency in Sperm | Frequency in Eggs |
| --- | --- | --- |
| A1B1 | y1 | x1 |
| A1B2 | y2 | x2 |
| A2B1 | y3 | x3 |
| A2B2 | y4 | x4 |

##### The recursion equations for loci on autosomes are -

y1’ = [2x1 y1m11 + (x1 y2 + x2 y1)m12 + (x1 y3 + x3 y1)m21 + (1 − rm)m22C(x1 y4 + x4 y1) + rmm22R(x2 y3 + x3 y2)]/ 2wm

y2’ = [2x2 y2m13 + (x1 y2 + x2 y1)m12 + (x2 y4 + x4 y2)m23 + (1 − rm)m22R(x3 y2 + x2 y3) + rmm22C(x4 y1 + x1 y4) ]/2wm

y3’ = [2x3 y3m31 + (x1 y3 + x3 y1)m21 + (x3 y4 + x4 y3)m32 + (1 − rm)m22R(x2 y3 + x3 y2) + rmm22C(x4 y1 + x1 y4)]/ 2wm

y4’ =[ 2x4 y4m33 + (x2 y4 + x4 y2)m23 + (x4 y3 + x3 y4)m32 + (1 − rm)m22C(x4 y1 + x1 y4) + rmm22R(x2 y3 + x3 y2)]/2 wm

x1’ = [2x1 y1 f11 + (x1 y2 + x2 y1) f12 + (x1 y3 + x3 y1) f21 + (1 − rf ) f22C(x1 y4 + x4 y1) + rf f22R(x2 y3 + x3 y2)]/2wf

x2’ = [2x2 y2 f13 + (x1 y2 + x2 y1) f12 + (x2 y4 + x4 y2) f23 + (1 − rf ) f22R(x2 y3 + x3 y2) + rf f22C(x4 y1 + x1 y4)]/2w f

x3’ = [2x3 y3 f31 + (x1 y3 + x3 y1) f21 + (x3 y4 + x4 y3) f32 + (1 − rf ) f22R(x2 y3 + x3 y2) + rf f22C(x4 y1 + x1 y4)]/2w f

x4’ = [2x4 y4 f33 + (x2 y4 + x4 y2) f23 + (x4 y3 + x3 y4) f32 + (1 − rf ) f22C(x4 y1 + x1 y4) + rf f22R(x2 y3 + x3 y2)]/2w f

Where mean fitnesses (wm and wf) are calculated as -

wm = x1 y1m11 + (x1 y2 + x2 y1)m12 + x2 y2m13 + (x1 y3 + x3 y1)m21 + (x2 y3 + x3 y2)m22R + (x1 y4 + x4 y1)m22C + (x2 y4 + x4 y2)m23 + x3 y3m31 + (x3 y4 + x4 y3)m32 + x4 y4m33

wf = x1 y1 f11 + (x1 y2 + x2 y1) f12 + x2 y2 f13 + (x1 y3 + x3 y1) f21 + (x2 y3 + x3 y2) f22R + (x1 y4 + x4 y1) f22C + (x2 y4 + x4 y2) f23 + x3 y3 f31 + (x3 y4 + x4 y3) f32 + x4 y4 f33.

The equations for the haplotype frequencies in females remain the same as before when the loci are X-linked. Following Connallon and Clark (2010), the recursion equations for when both loci are X-linked are as follows -

y1’ = x1m1 wm

y2’ = x2m2 wm

y3’ = x3m3 wm

y4’ = x4m4 wm

Where mean fitness in males is defined as -

wm = x1m1 + x2m2 + x3m3 + x4m4

### Selection Coefficients and Dominance Coefficient

##### **The effect of varying dominance coefficient and selection coefficient on the frequency of A2B2 haplotype when the loci that are present on autosomes**

**Figure - S1.1**

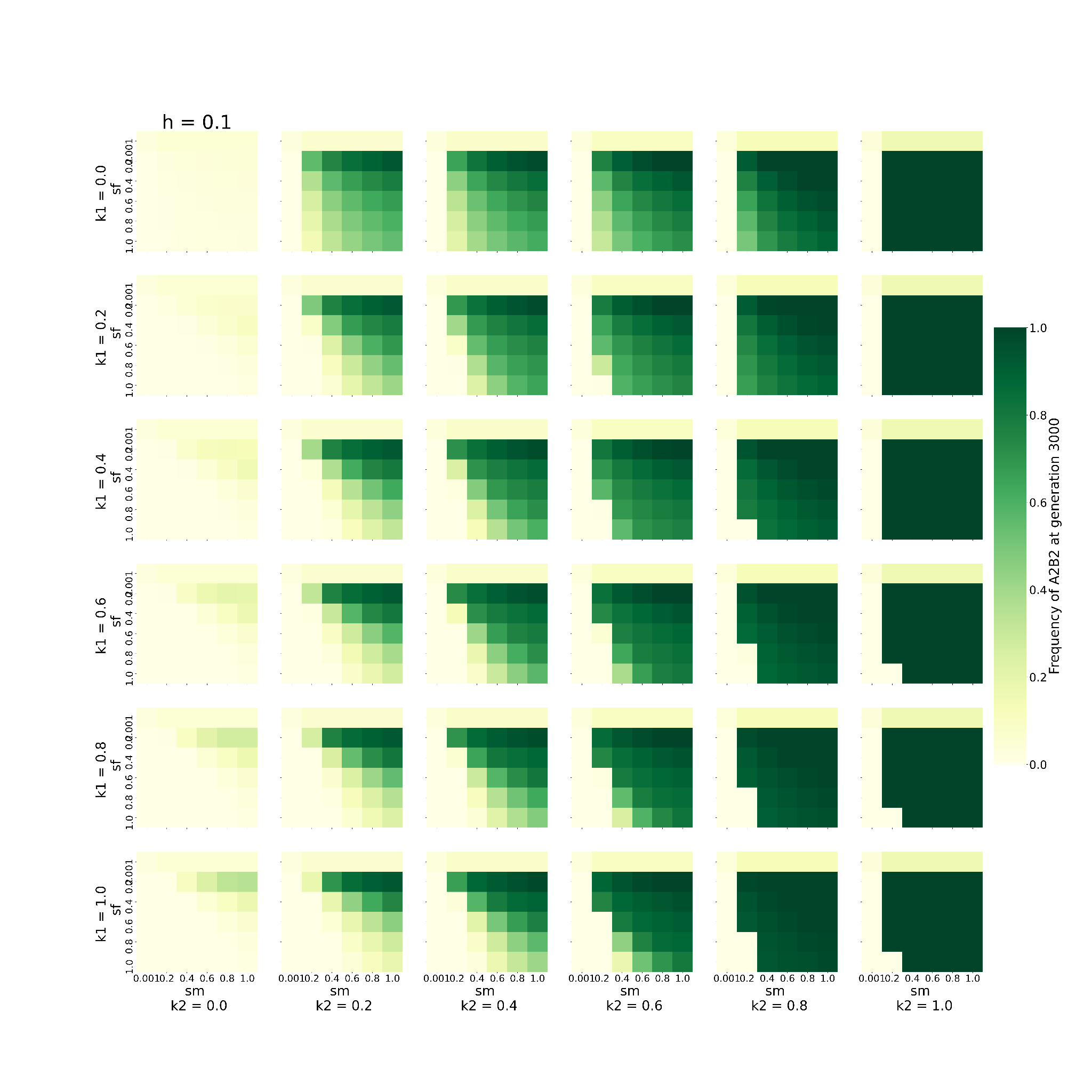

This graph shows the behaviour of the system when varying 2 of the 6 parameters, namely, sm, and sf, keeping h fixed to 0.1 (h= 0.1), given that the loci are present on the autosome. The remaining 3 parameters are kept constant at 0.2 except for D which is fixed to 0.0. Every block shows one combination of k1 and k2. k1 increases downwards, while k2 increases rightwards. The individual X axis for every k1-k2 denotes values of sm increasing *towards right* and the Y axis represents values of sf *increasing downwards*. Every box (sm-sf combination) is coloured to denote the frequency of A2B2 haplotype at 3000th generation. The greener the colour, the higher the frequency.

**Figure - S1.2**

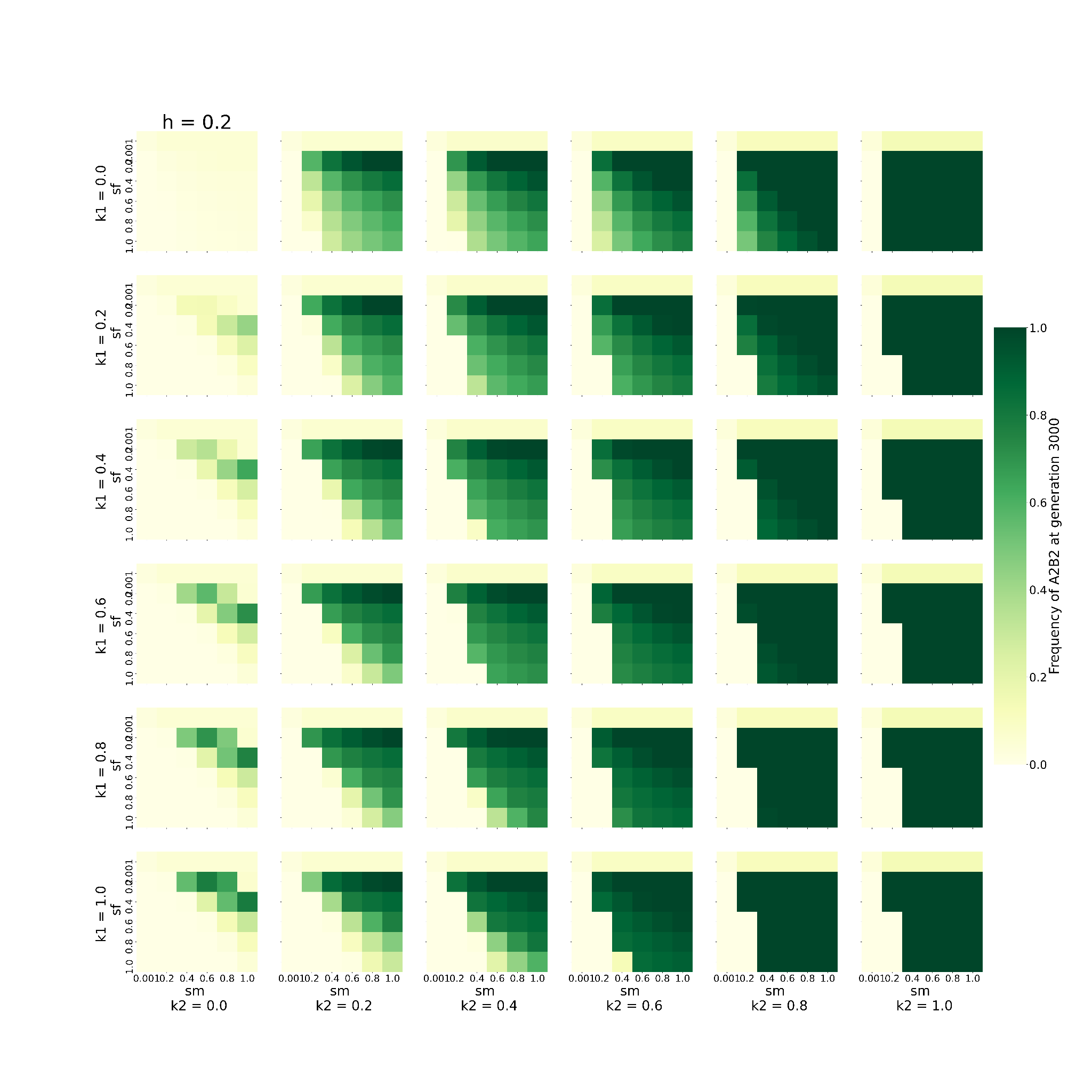

This graph shows the behaviour of the system when varying 2 of the 6 parameters, namely, sm, and sf, keeping h fixed to 0.2 (h= 0.2), given that the loci are present on the autosome. The remaining 3 parameters are kept constant at 0.2 except for D which is fixed to 0.0.. Every block shows one combination of k1 and k2. k1 increases downwards, while k2 increases rightwards. The individual X axis for every k1-k2 denotes values of sm increasing *towards right* and the Y axis represents values of sf *increasing downwards*. Every box (sm-sf combination) is coloured to denote the frequency of A2B2 haplotype at 3000th generation. The greener the colour, the higher the frequency.

**Figure - S1.3**

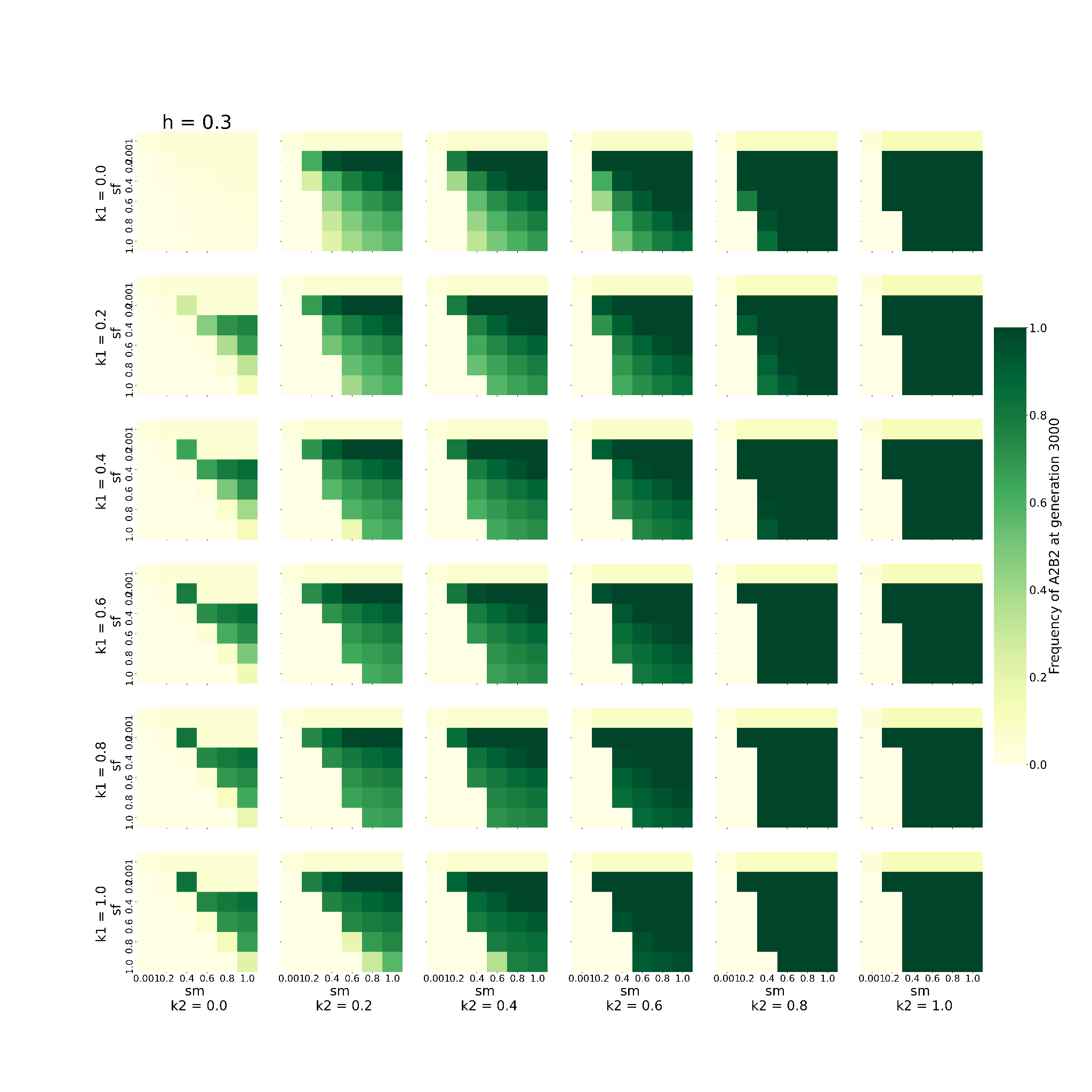

This graph shows the behaviour of the system when varying 2 of the 6 parameters, namely, sm, and sf, keeping h fixed to 0.3 (h= 0.3), given that the loci are present on the autosome. The remaining 3 parameters are kept constant at 0.2 except for D which is fixed to 0.0. Every block shows one combination of k1 and k2. k1 increases downwards, while k2 increases rightwards. The individual X axis for every k1-k2 denotes values of values of sm increasing *towards right* and the Y axis represents values of sf *increasing downwards*. Every box (sm-sf combination) is coloured to denote the frequency of A2B2 haplotype at 3000th generation. The greener the colour, the higher the frequency.

**Figure - S1.4**

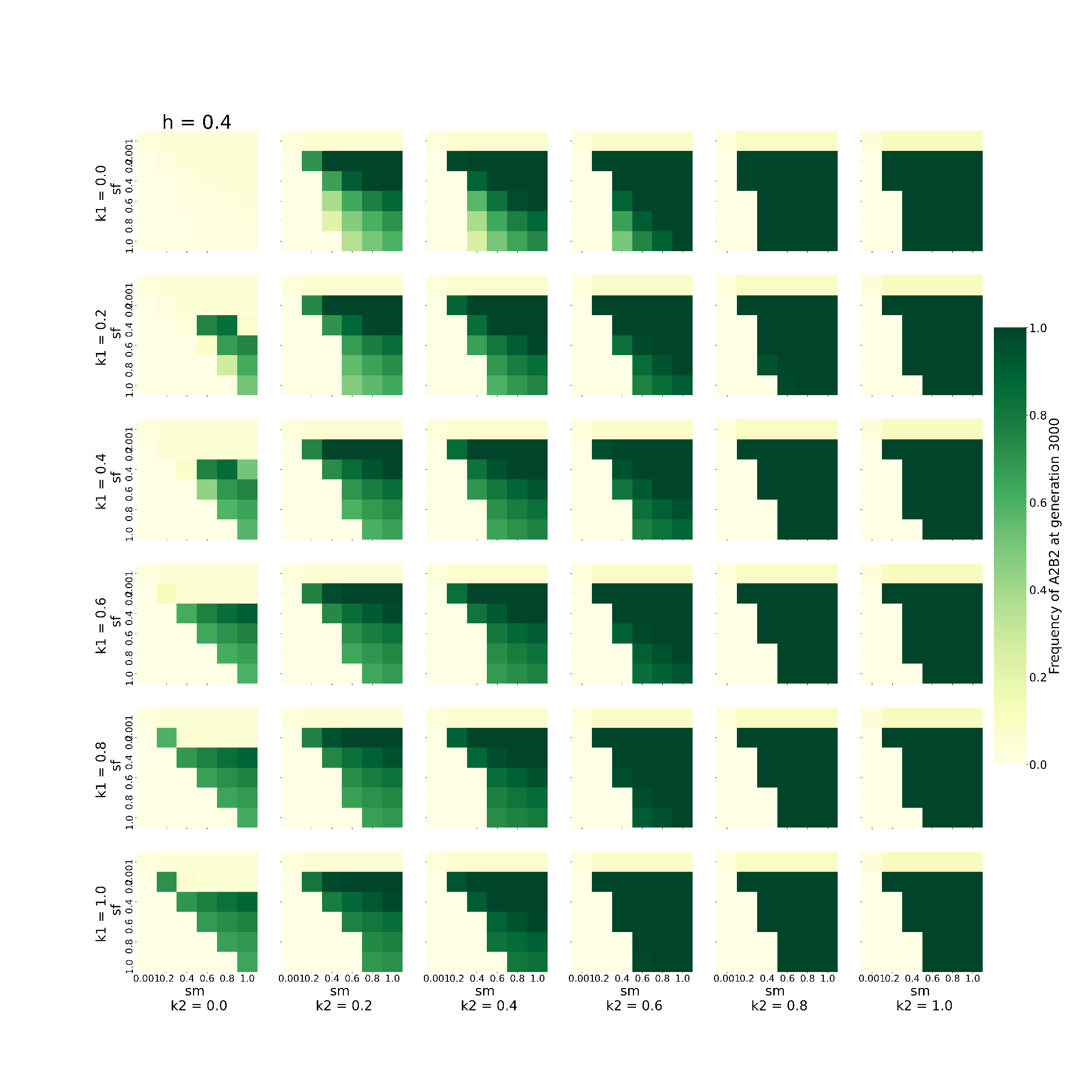

This graph shows the behaviour of the system when varying 2 of the 6 parameters, namely, sm, and sf, keeping h fixed to 0.4 (h= 0.4), given that the loci are present on the autosome. The remaining 3 parameters are kept constant at 0.2 except for D which is fixed to 0.0.. Every block shows one combination of k1 and k2. k1 increases downwards, while k2 increases rightwards. The individual X axis for every k1-k2 denotes values of sm increasing *towards right* and the Y axis represents values of sf *increasing downwards*. Every box (sm-sf combination) is coloured to denote the frequency of A2B2 haplotype at 3000th generation. The greener the colour, the higher the frequency.

**Figure - S1.5**

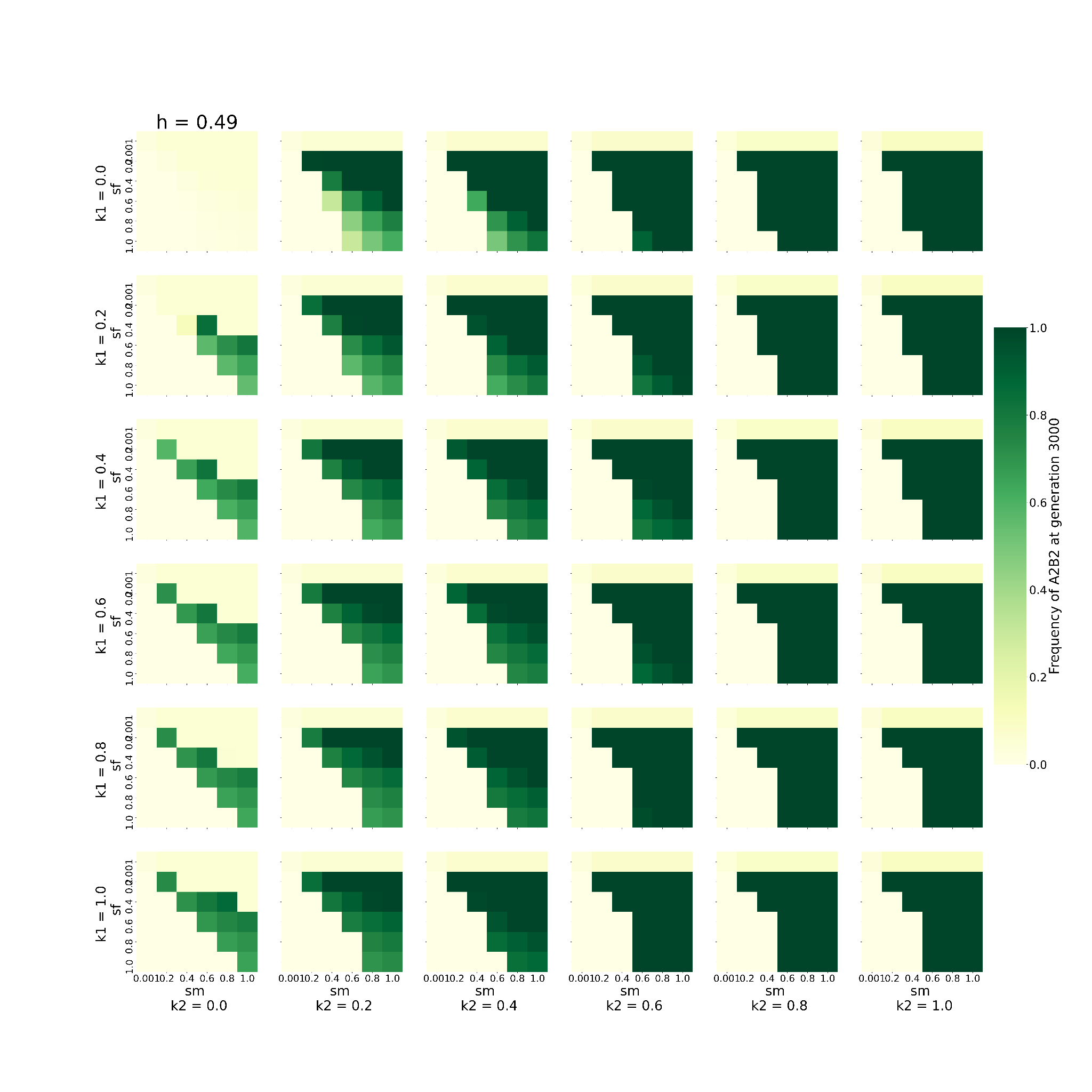

This graph shows the behaviour of the system when varying 2 of the 6 parameters, namely, sm, and sf, keeping h fixed to 0.49 (h= 0.49), given that the loci are present on the autosome. The remaining 3 parameters are kept constant at 0.2 except for D which is fixed to 0.0. Every block shows one combination of k1 and k2. k1 increases downwards, while k2 increases rightwards. The individual X axis for every k1-k2 denotes values of sm increasing *towards right* and the Y axis represents values of sf *increasing downwards*. Every box (sm-sf combination) is coloured to denote the frequency of A2B2 haplotype at 3000th generation. The greener the colour, the higher the frequency.

##### **The effect of varying dominance coefficient and selection coefficient on the time taken by A2B2 haplotype to get fixed when the loci are present on autosomes**

**Figure - S2.1**

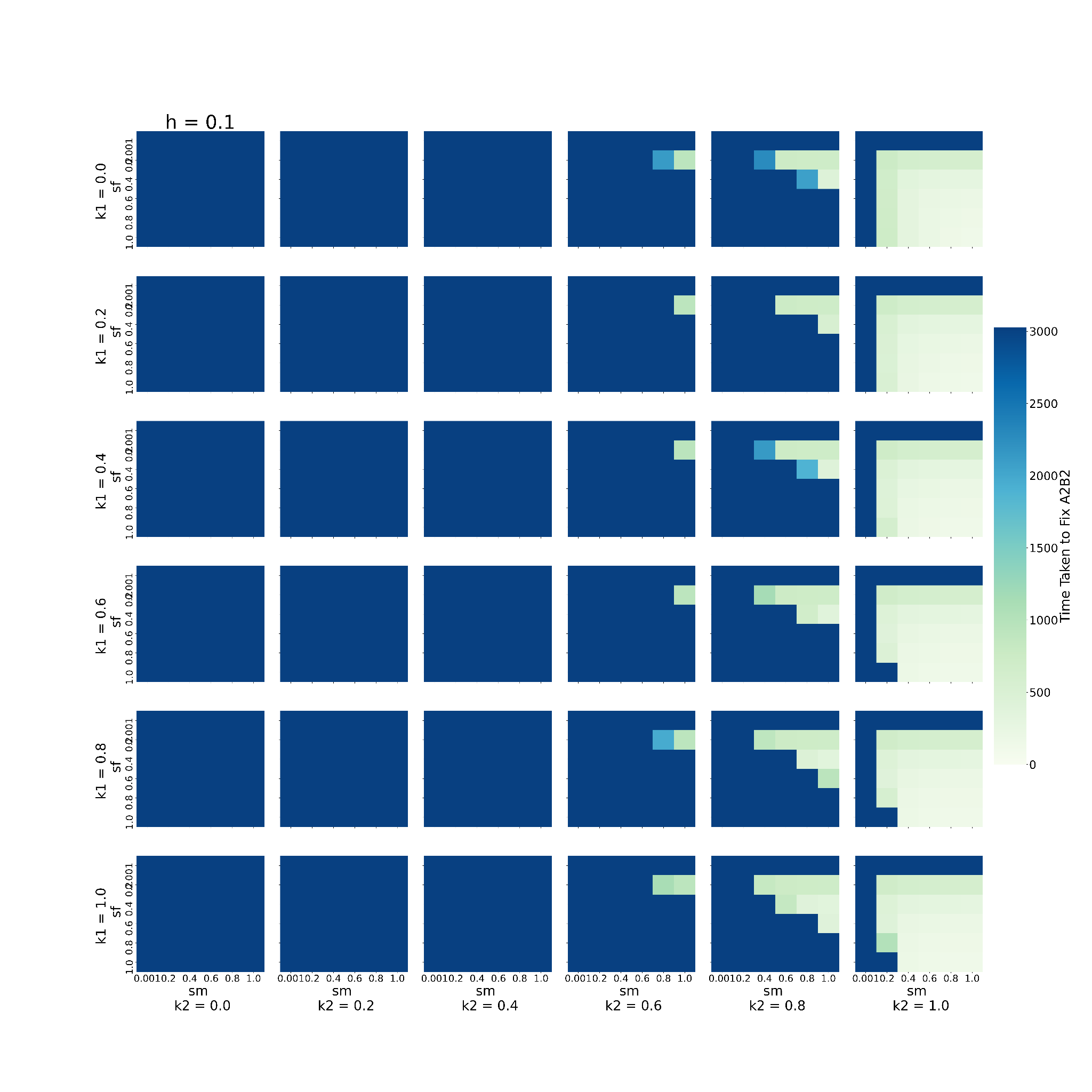

This graph shows the behaviour of the system when varying 2 of the 6 parameters, namely, sm, and sf, keeping h fixed to 0.1 (h= 0.1), given that the loci are present on the autosome. The remaining 3 parameters are kept constant at 0.2 except for D which is fixed to 0.0. Every block shows one combination of k1 and k2. k1 increases downwards, while k2 increases rightwards. The individual X axis for every k1-k2 denotes values of sm increasing *towards right* and the Y axis represents values of sf *increasing downwards*. Every box (sm-sf combination) is coloured to denote the time taken to fix A2B2 in the population. Bluer colours indicate the higher time taken to fix A2B2.

**Figure - S2.2**

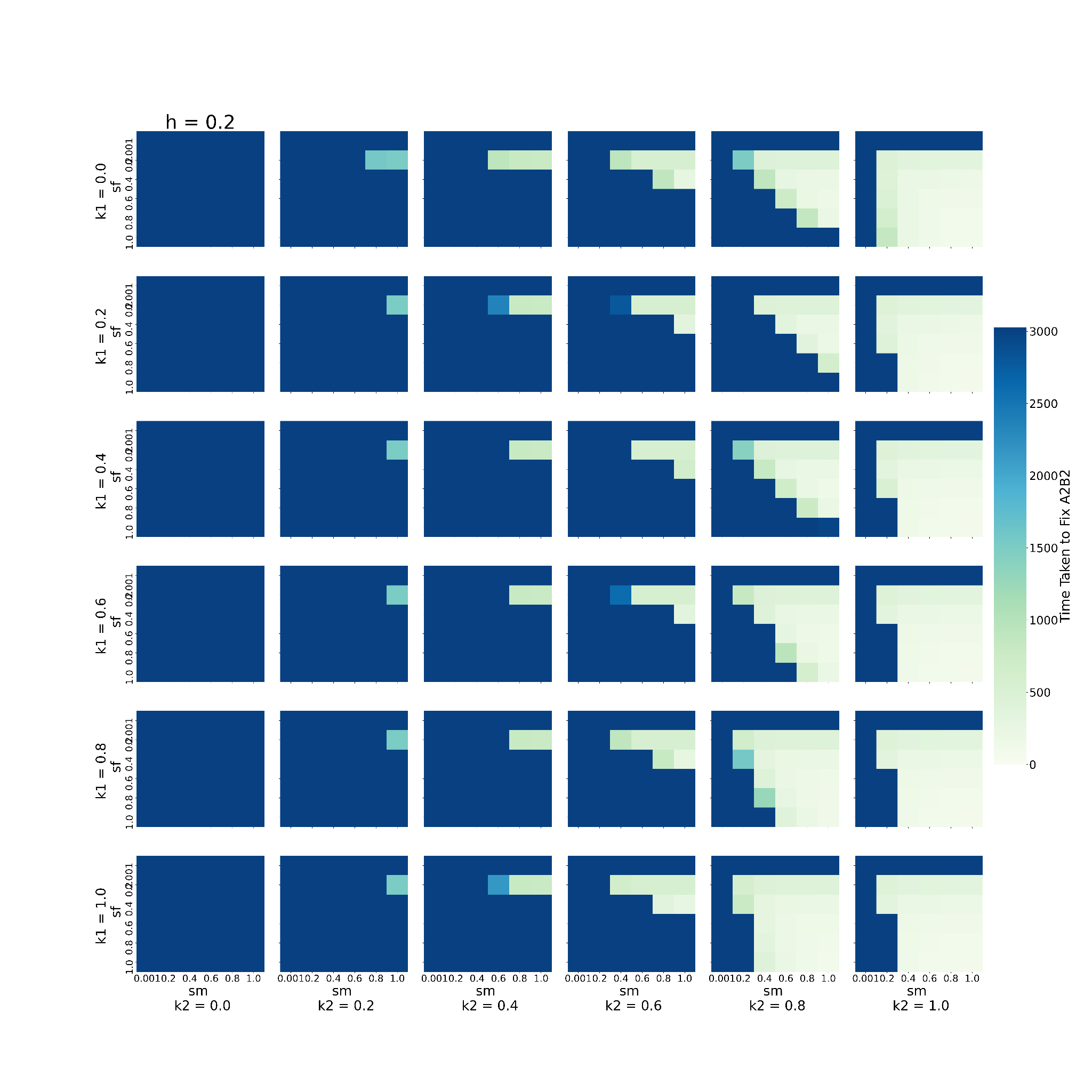

This graph shows the behaviour of the system when varying 2 of the 6 parameters, namely, sm, and sf, keeping h fixed to 0.2 (h= 0.2), given that the loci are present on the autosome. The remaining 3 parameters are kept constant at 0.2 except for D which is fixed to 0.0.. Every block shows one combination of k1 and k2. k1 increases downwards, while k2 increases rightwards. The individual X axis for every k1-k2 denotes values of sm increasing *towards right* and the Y axis represents values of sf *increasing downwards*. Every box (sm-sf combination) is coloured to denote the time taken to fix A2B2 in the population. Bluer colours indicate the higher time taken to fix A2B2.

**Figure - S2.3**

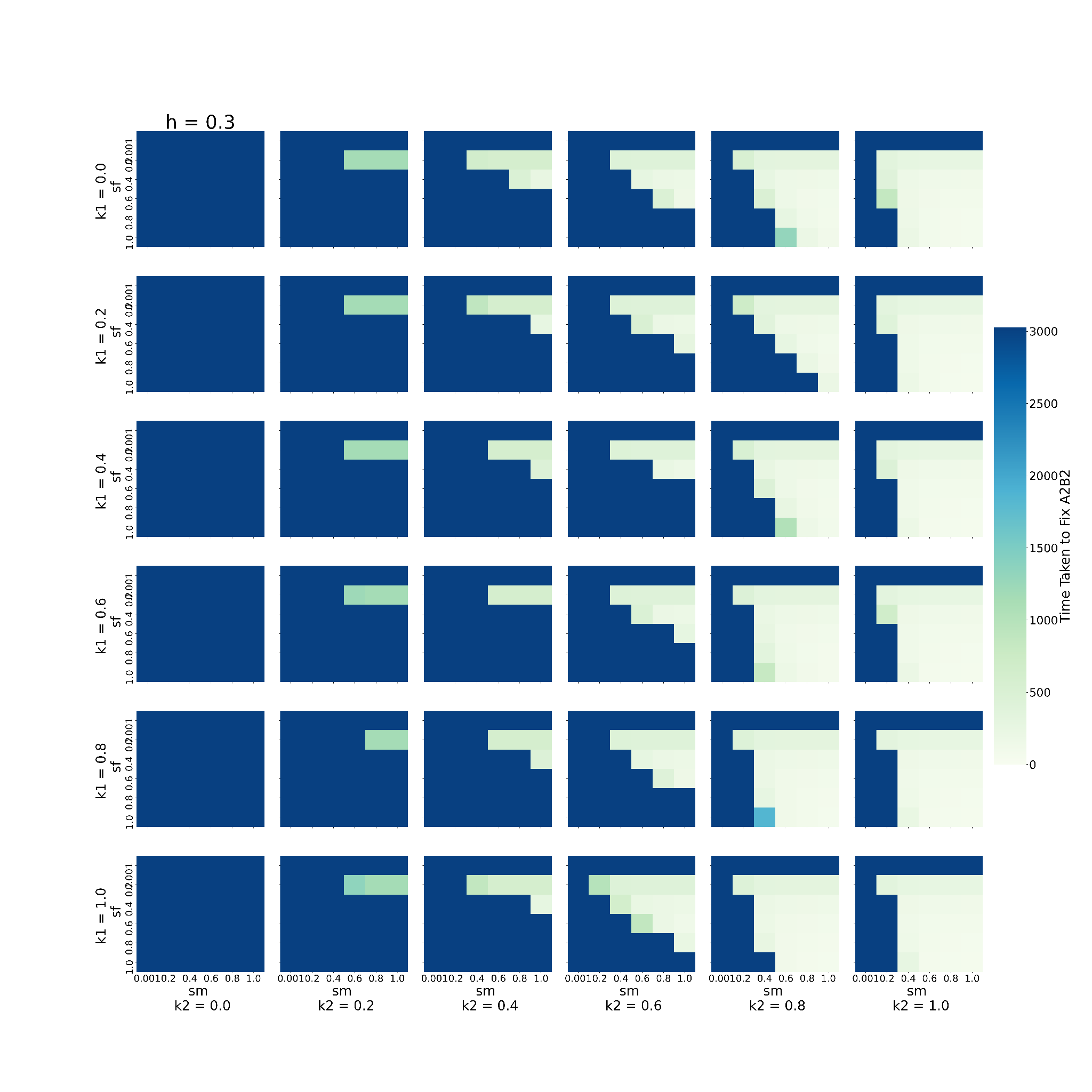

This graph shows the behaviour of the system when varying 2 of the 6 parameters, namely, sm, and sf, keeping h fixed to 0.3 (h= 0.3), given that the loci are present on the autosome. The remaining 3 parameters are kept constant at 0.2 except for D which is fixed to 0.0. Every block shows one combination of k1 and k2. k1 increases downwards, while k2 increases rightwards. The individual X axis for every k1-k2 denotes values of sm increasing *towards right* and the Y axis represents values of sf *increasing downwards*. Every box (sm-sf combination) is coloured to denote the time taken to fix A2B2 in the population. Bluer colours indicate the higher time taken to fix A2B2.

**Figure - S2.4**

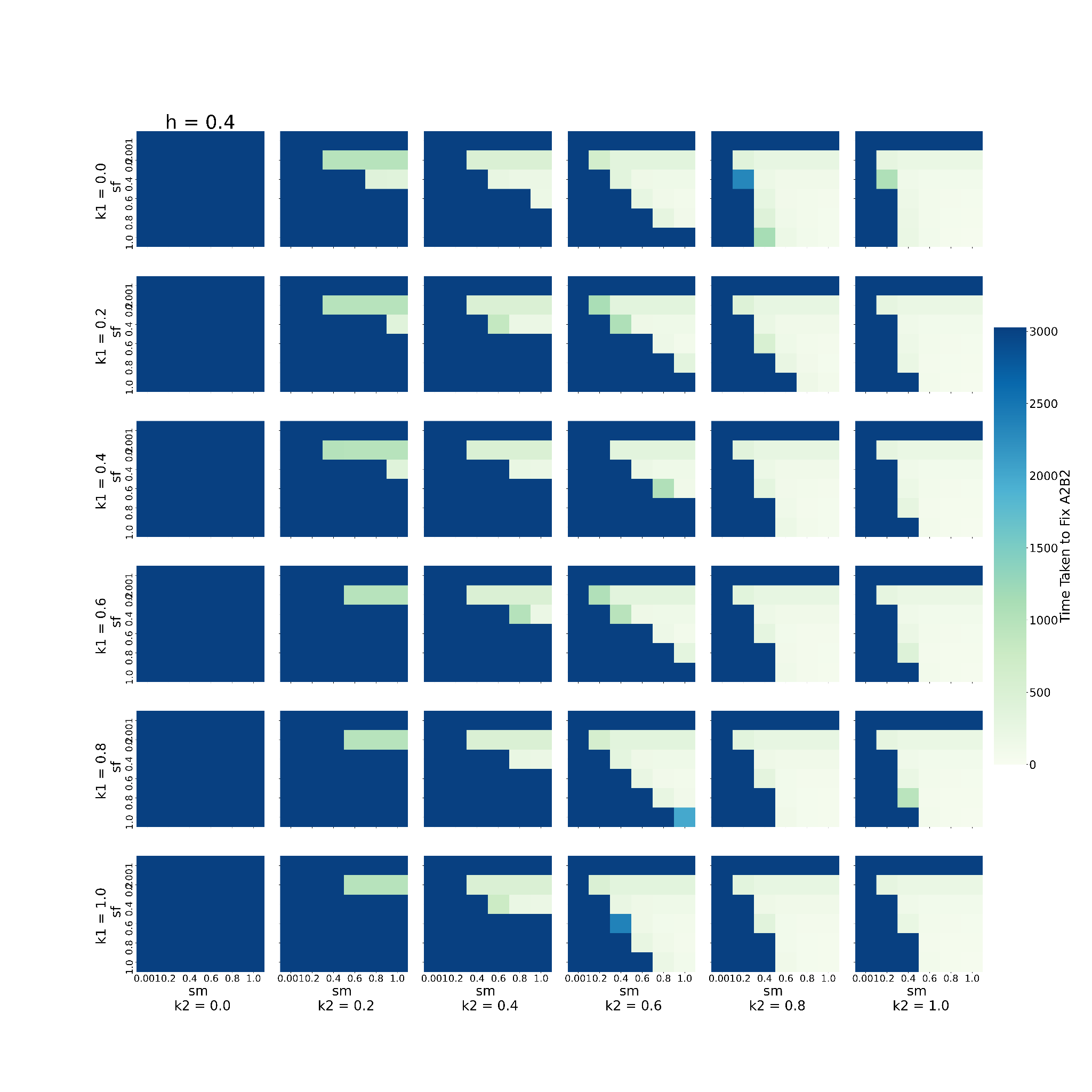

This graph shows the behaviour of the system when varying 2 of the 6 parameters, namely, sm, and sf, keeping h fixed to 0.4 (h= 0.4), given that the loci are present on the autosome. The remaining 3 parameters are kept constant at 0.2 except for D which is fixed to 0.0. Every block shows one combination of k1 and k2. k1 increases downwards, while k2 increases rightwards. The individual X axis for every k1-k2 denotes values of sm increasing *towards right* and the Y axis represents values of sf *increasing downwards*. Every box (sm-sf combination) is coloured to denote the time taken to fix A2B2 in the population. Bluer colours indicate the higher time taken to fix A2B2.

**Figure - S2.5**

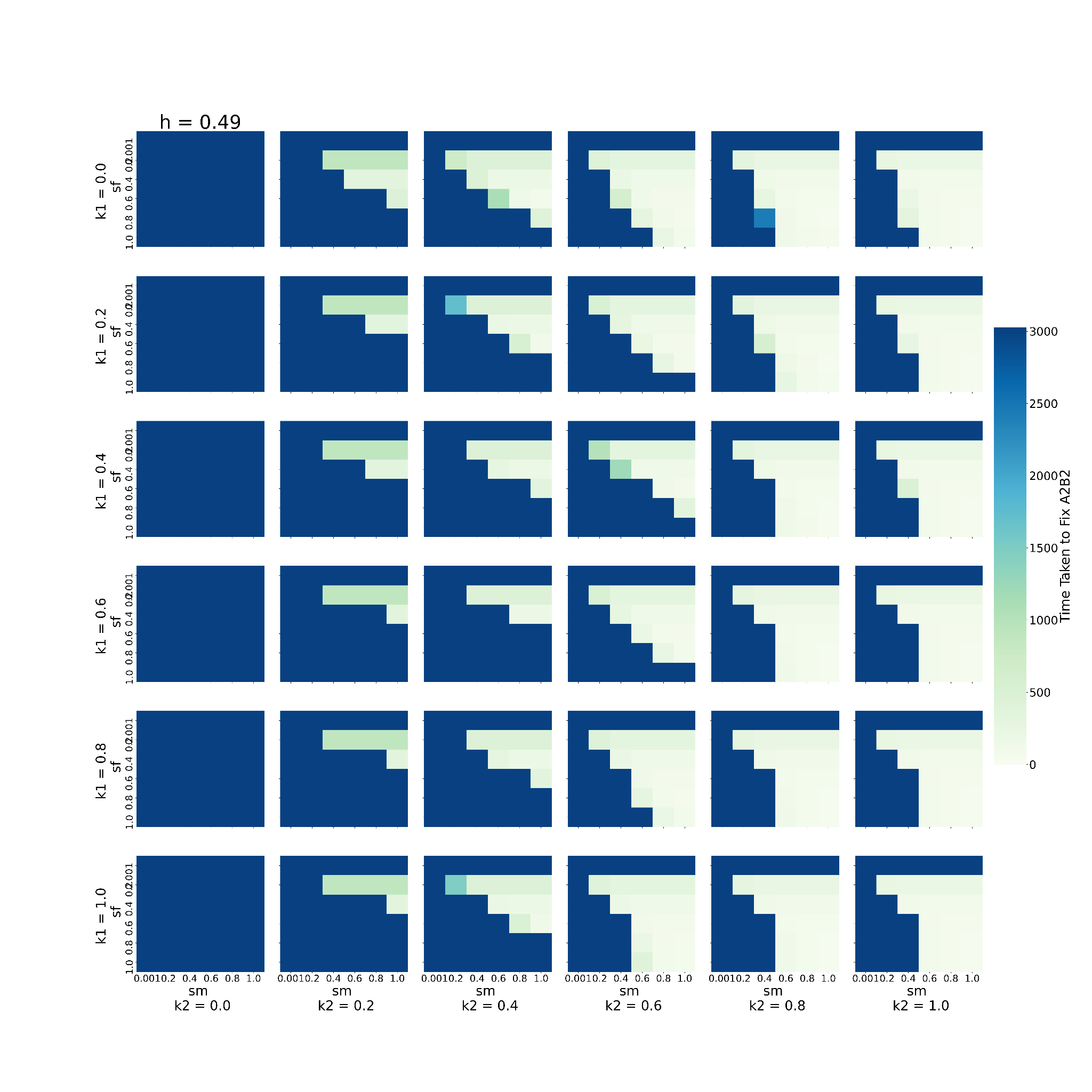

This graph shows the behaviour of the system when varying 2 of the 6 parameters, namely, sm, and sf, keeping h fixed to 0.49 (h= 0.49), given that the loci are present on the autosome. The remaining 3 parameters are kept constant at 0.2 except for D which is fixed to 0.0. Every block shows one combination of k1 and k2. k1 increases downwards, while k2 increases rightwards. The individual X axis for every k1-k2 denotes values of sm increasing *towards right* and the Y axis represents values of sf *increasing downwards*. Every box (sm-sf combination) is coloured to denote the time taken to fix A2B2 in the population. Bluer colours indicate the higher time taken to fix A2B2.

##### **The effect of varying dominance coefficient and selection coefficient on the frequency of A2B2 haplotype when the loci are X-linked**

**Figure - S3.1**

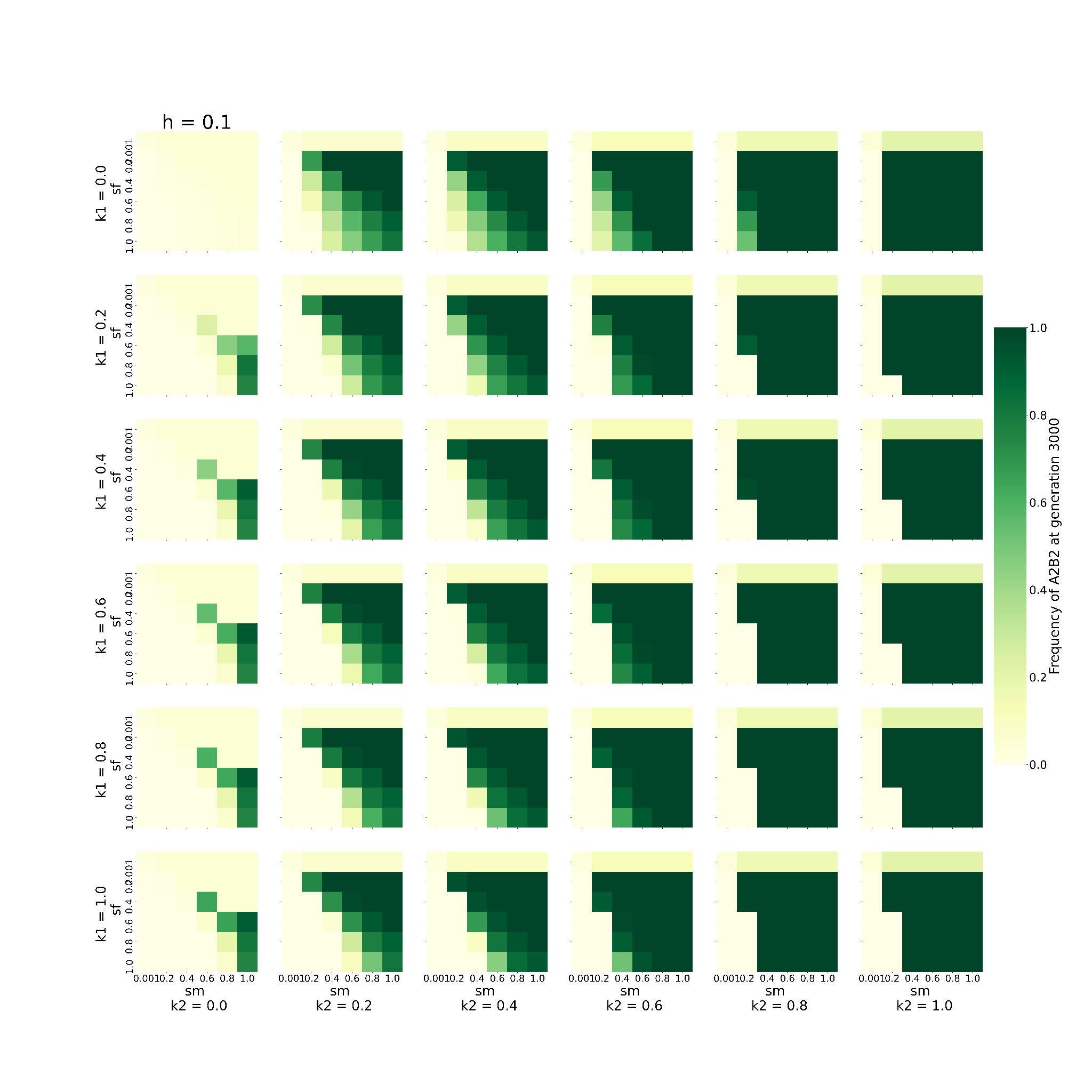

This graph shows the behaviour of the system when varying 2 of the 6 parameters, namely, sm, and sf, keeping h fixed to 0.1 (h= 0.1), given that the loci are X-linked. The remaining 3 parameters are kept constant at 0.2, except for D which is fixed to 0. Every block shows one combination of k1 and k2. k1 increases downwards, while k2 increases rightwards. The individual X axis for every k1-k2 denotes values of sm increasing *towards right* and the Y axis represents values of sf *increasing downwards*. Every box (sm-sf combination) is coloured to denote the frequency of A2B2 haplotype at 3000th generation. The greener the colour, the higher the frequency.

**Figure - S3.2**

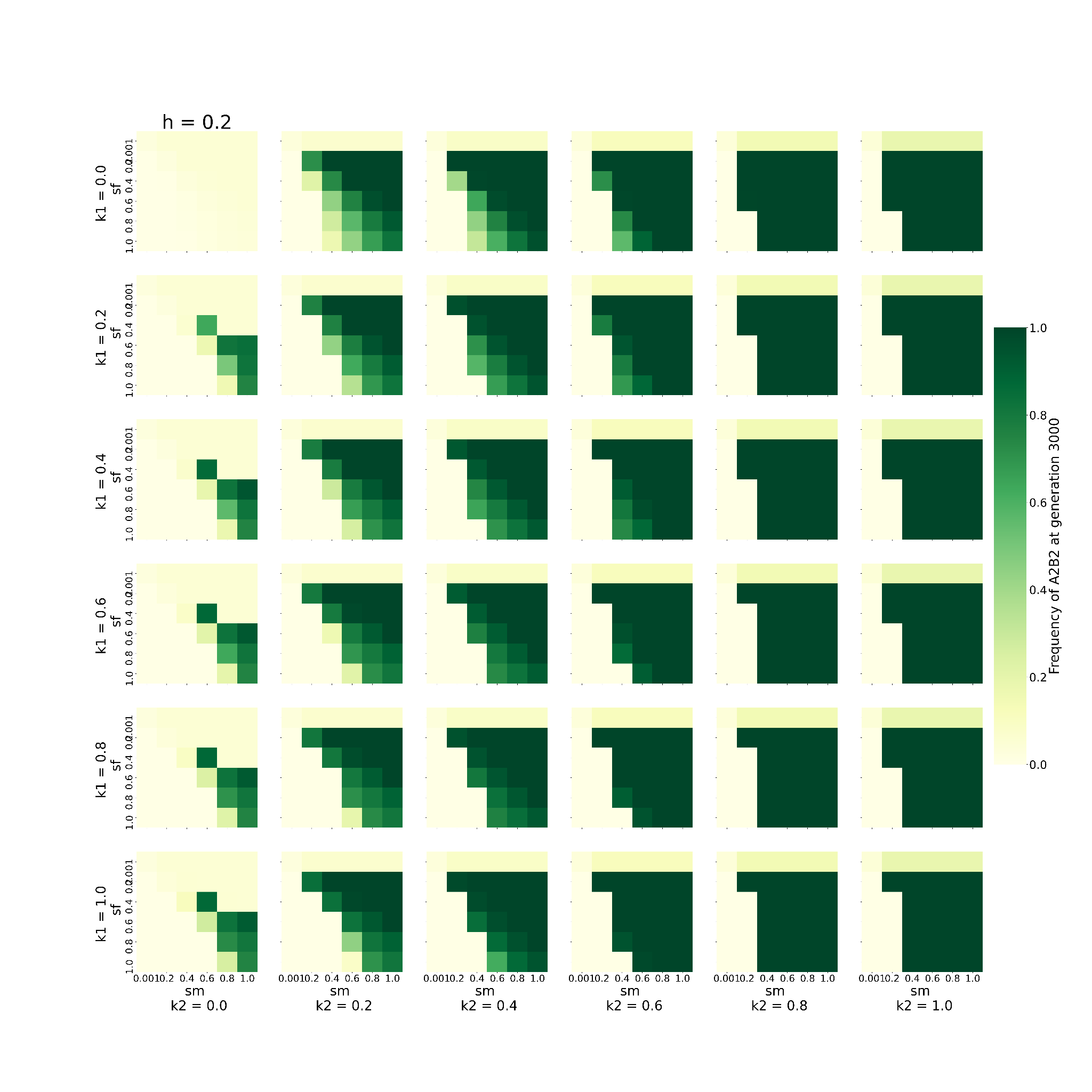

This graph shows the behaviour of the system when varying 2 of the 6 parameters, namely, sm, and sf, keeping h fixed to 0.2 (h= 0.2), given that the loci are X-linked. The remaining 3 parameters are kept constant at 0.2, except for D which is fixed to 0.. Every block shows one combination of k1 and k2. k1 increases downwards, while k2 increases rightwards. The individual X axis for every k1-k2 denotes values of sm increasing *towards right* and the Y axis represents values of sf *increasing downwards*. Every box (sm-sf combination) is coloured to denote the frequency of A2B2 haplotype at 3000th generation. The greener the colour, the higher the frequency.

**Figure - S3.3**

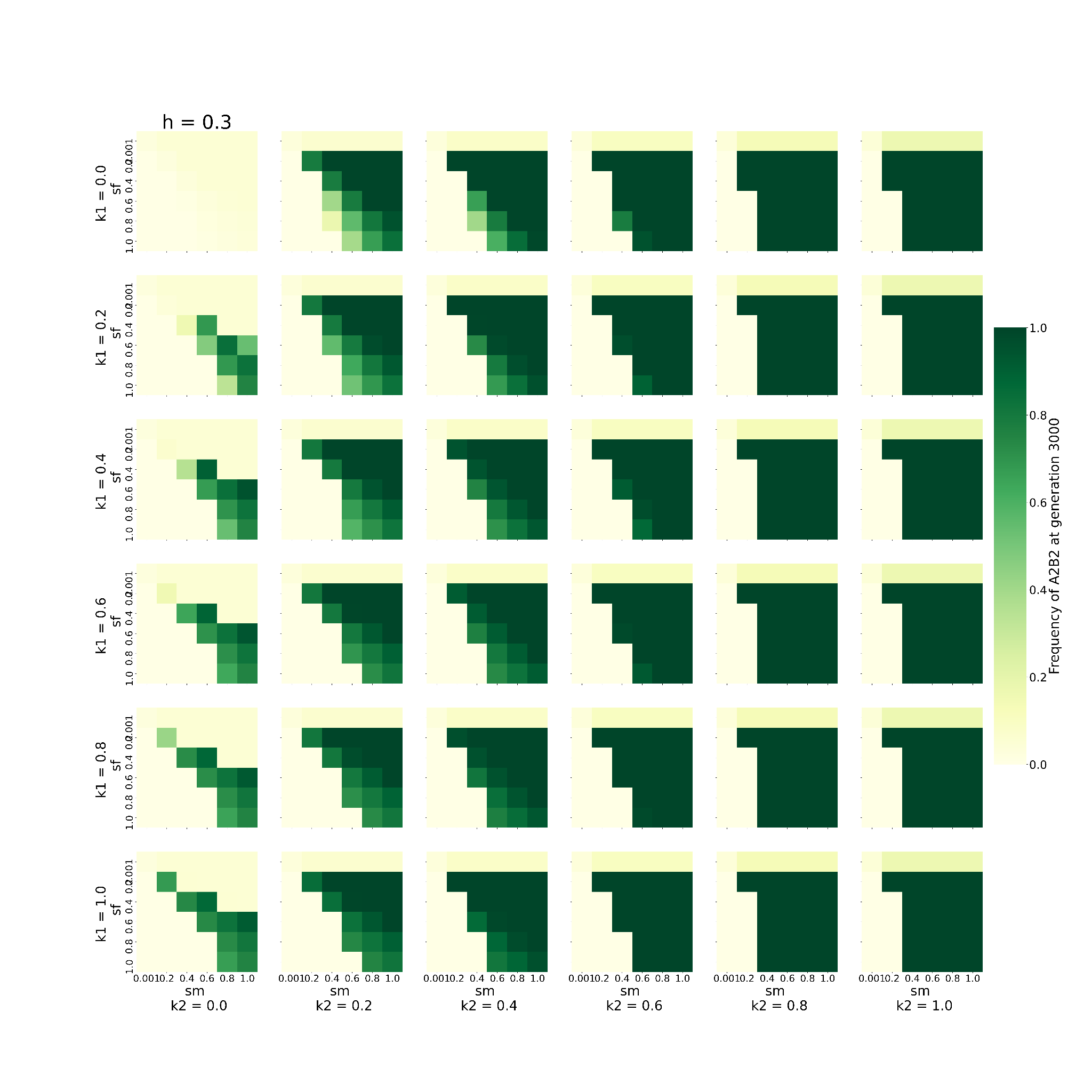

This graph shows the behaviour of the system when varying 2 of the 6 parameters, namely, sm, and sf, keeping h fixed to 0.3 (h= 0.3), given that the loci are X-linked. The remaining 3 parameters are kept constant at 0.2, except for D which is fixed to 0. Every block shows one combination of k1 and k2. k1 increases downwards, while k2 increases rightwards. The individual X axis for every k1-k2 denotes values of sm increasing *towards right* and the Y axis represents values of sf *increasing downwards*. Every box (sm-sf combination) is coloured to denote the frequency of A2B2 haplotype at 3000th generation. The greener the colour, the higher the frequency.

**Figure - S3.4**

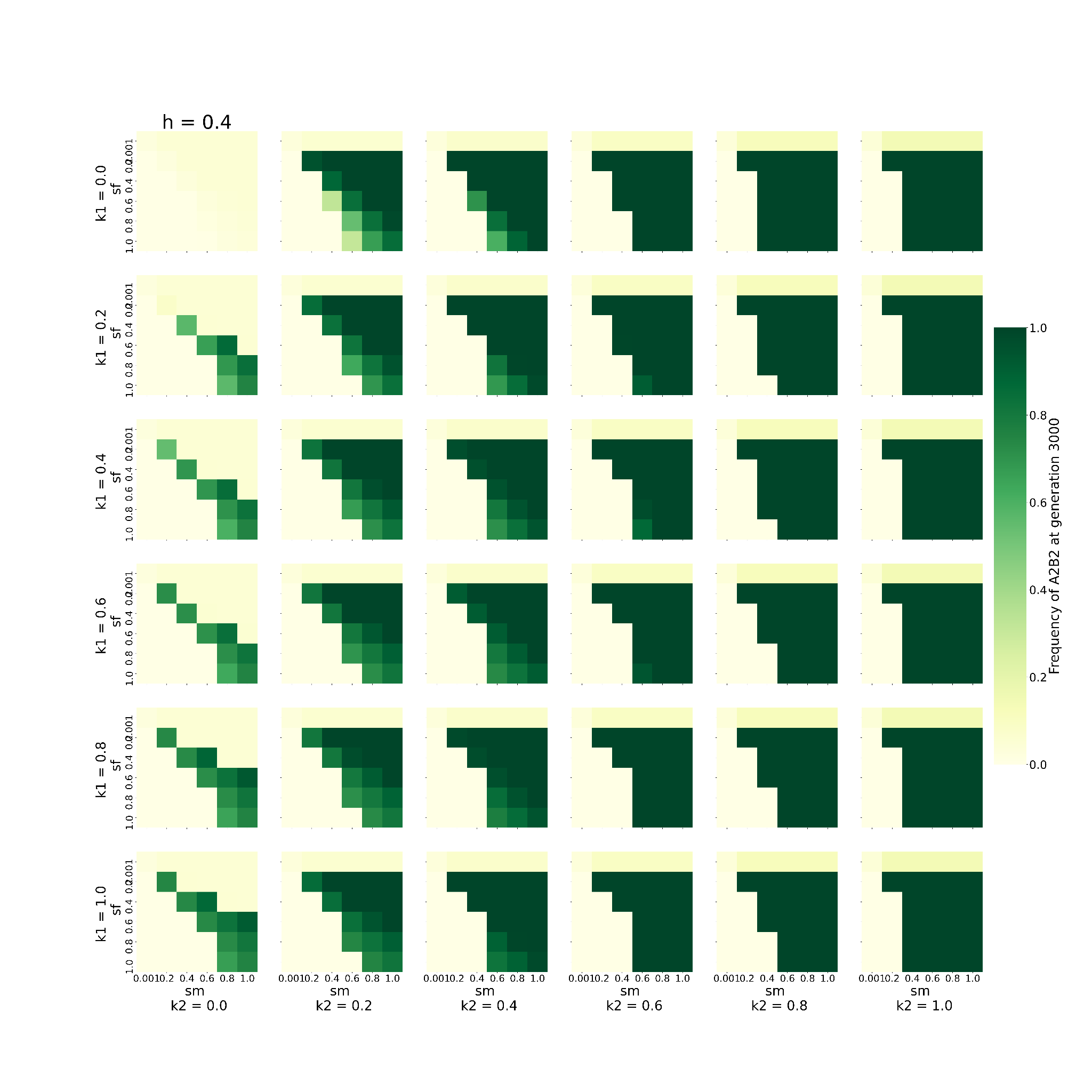

This graph shows the behaviour of the system when varying 2 of the 6 parameters, namely, sm, and sf, keeping h fixed to 0.4 (h= 0.4), given that the loci are X-linked. The remaining 3 parameters are kept constant at 0.2, except for D which is fixed to 0. Every block shows one combination of k1 and k2. k1 increases downwards, while k2 increases rightwards. The individual X axis for every k1-k2 denotes values of sm increasing *towards right* and the Y axis represents values of sf *increasing downwards*. Every box (sm-sf combination) is coloured to denote the frequency of A2B2 haplotype at 3000th generation. The greener the colour, the higher the frequency.

**Figure - S3.5**

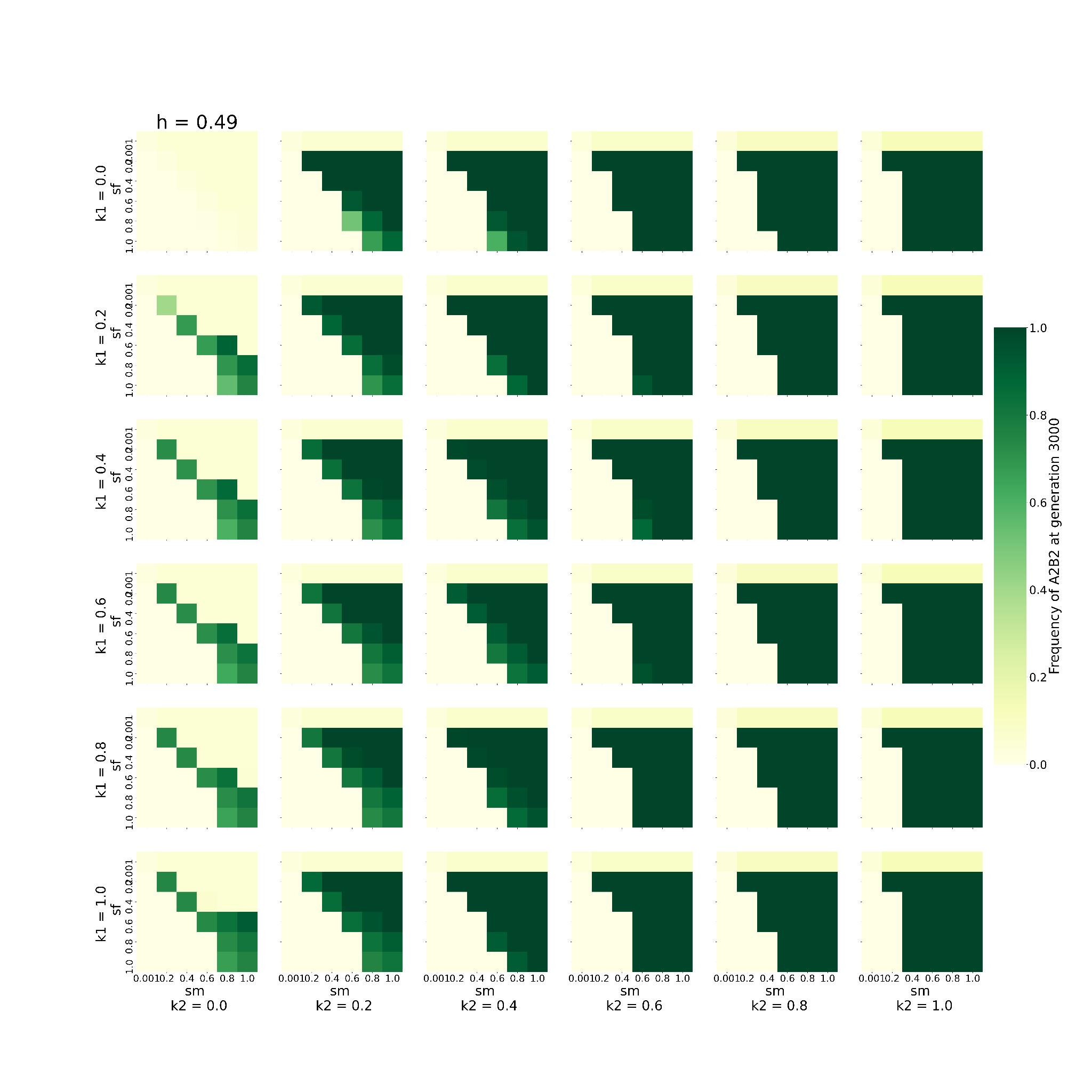

This graph shows the behaviour of the system when varying 2 of the 6 parameters, namely, sm, and sf, keeping h fixed to 0.49 (h= 0.49), given that the loci are X-linked. The remaining 3 parameters are kept constant at 0.2, except for D which is fixed to 0. Every block shows one combination of k1 and k2. k1 increases downwards, while k2 increases rightwards. The individual X axis for every k1-k2 denotes values of sm increasing *towards right* and the Y axis represents values of sf *increasing downwards*. Every box (sm-sf combination) is coloured to denote the frequency of A2B2 haplotype at 3000th generation. The greener the colour, the higher the frequency.

##### **The effect of varying dominance coefficient and selection coefficient on the time taken by A2B2 haplotype to get fixed when the loci are X-linked**

**Figure - S4.1**

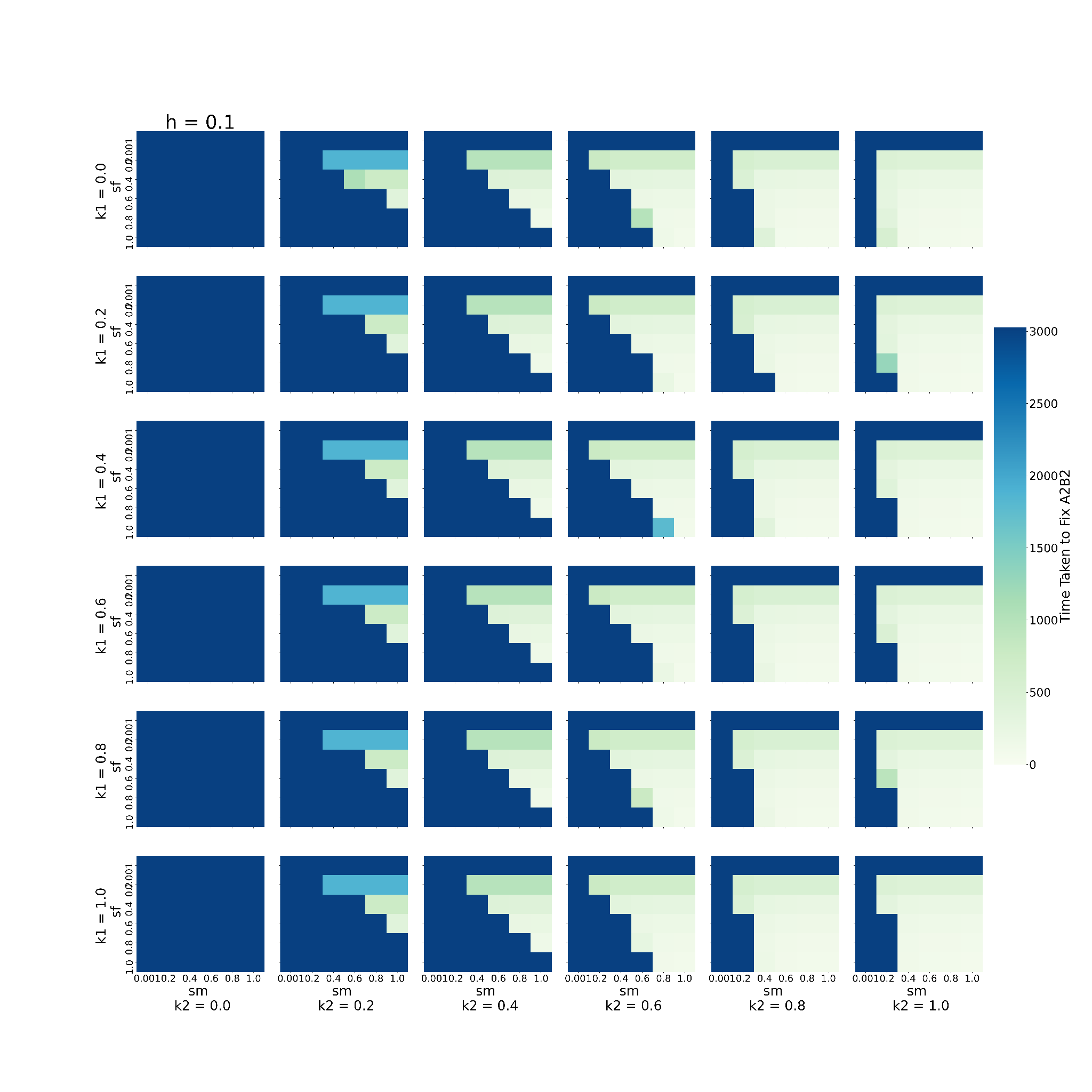

This graph shows the behaviour of the system when varying 2 of the 6 parameters, namely, sm, and sf, keeping h fixed to 0.1 (h= 0.1), given that the loci are X-linked. The remaining 3 parameters are kept constant at 0.2, except for D which is fixed to 0. Every block shows one combination of k1 and k2. k1 increases downwards, while k2 increases rightwards. The individual X axis for every k1-k2 denotes values of sm increasing *towards right* and the Y axis represents values of sf *increasing downwards*. Every box (sm-sf combination) is coloured to denote the time taken to fix A2B2 in the population. Bluer colours indicate the higher time taken to fix A2B2.

**Figure - S4.2**

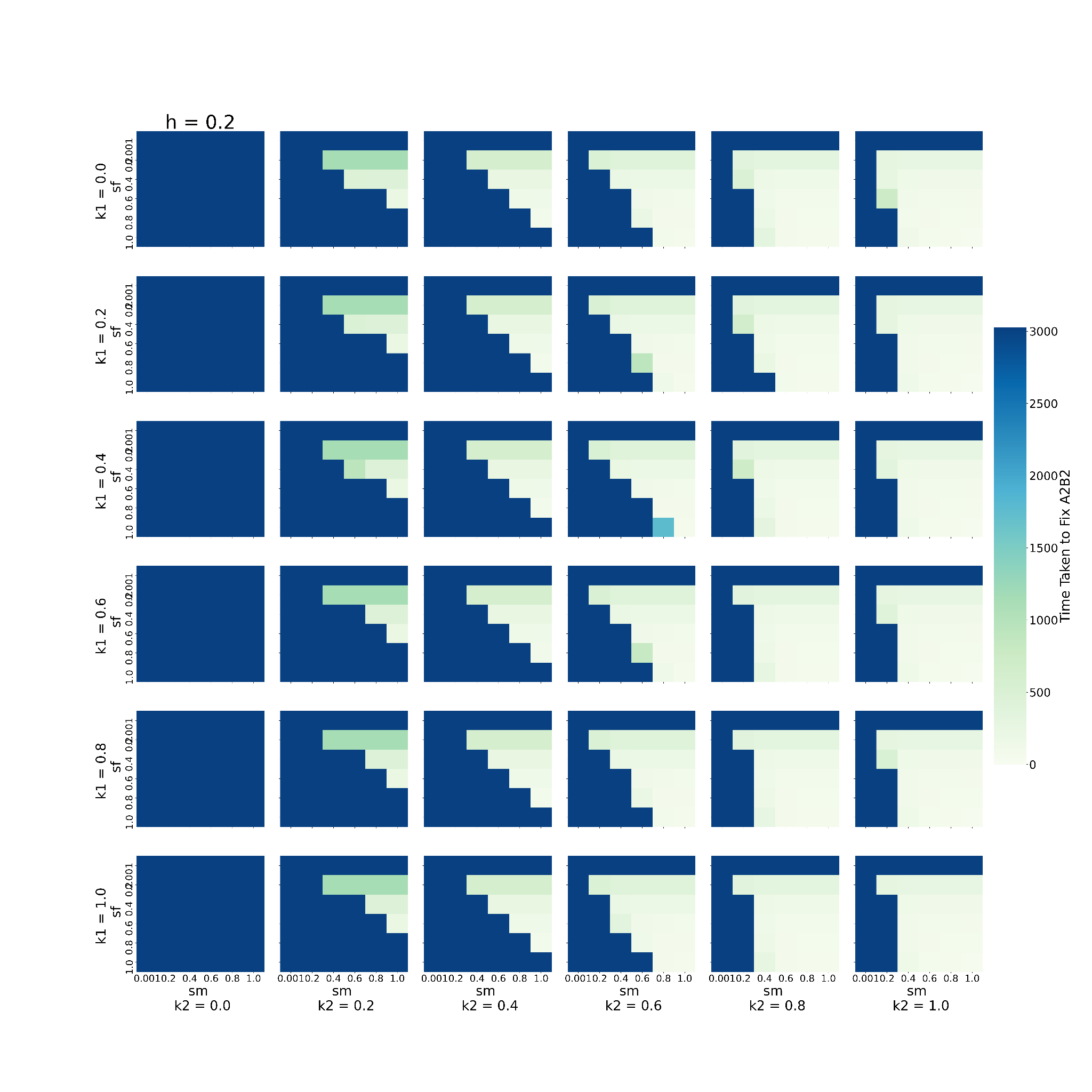

This graph shows the behaviour of the system when varying 2 of the 6 parameters, namely, sm, and sf, keeping h fixed to 0.2 (h= 0.2), given that the loci are X-linked. The remaining 3 parameters are kept constant at 0.2, except for D which is fixed to 0. Every block shows one combination of k1 and k2. k1 increases downwards, while k2 increases rightwards. The individual X axis for every k1-k2 denotes values of sm increasing *towards right* and the Y axis represents values of sf *increasing downwards*. Every box (sm-sf combination) is coloured to denote the time taken to fix A2B2 in the population. Bluer colours indicate the higher time taken to fix A2B2.

**Figure - S4.3**

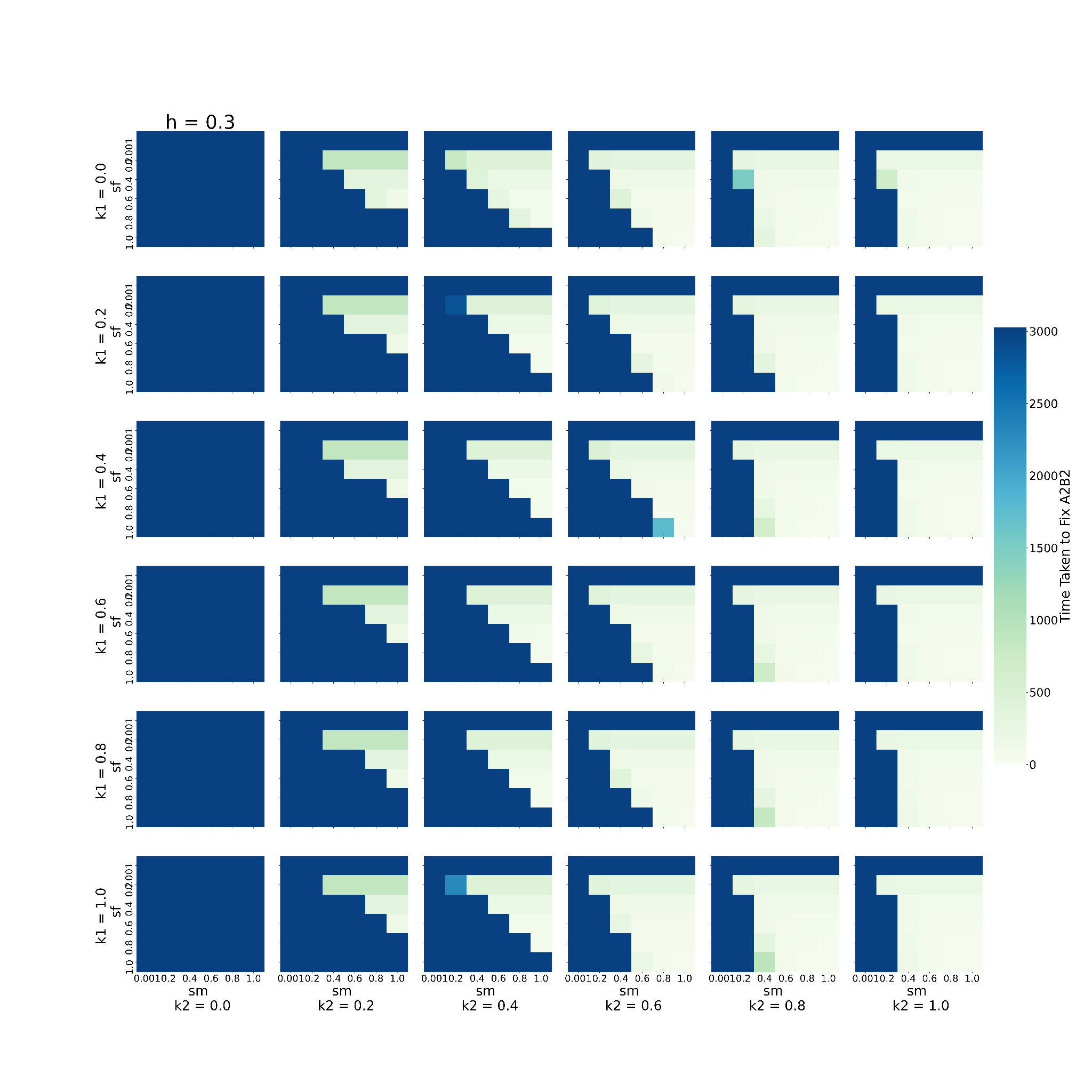

This graph shows the behaviour of the system when varying 2 of the 6 parameters, namely, sm, and sf, keeping h fixed to 0.3 (h= 0.3), given that the loci are X-linked. The remaining 3 parameters are kept constant at 0.2, except for D which is fixed to 0. Every block shows one combination of k1 and k2. k1 increases downwards, while k2 increases rightwards. The individual X axis for every k1-k2 denotes values of sm increasing *towards right* and the Y axis represents values of sf *increasing downwards*. Every box (sm-sf combination) is coloured to denote the time taken to fix A2B2 in the population. Bluer colours indicate the higher time taken to fix A2B2.

**Figure - S4.4**

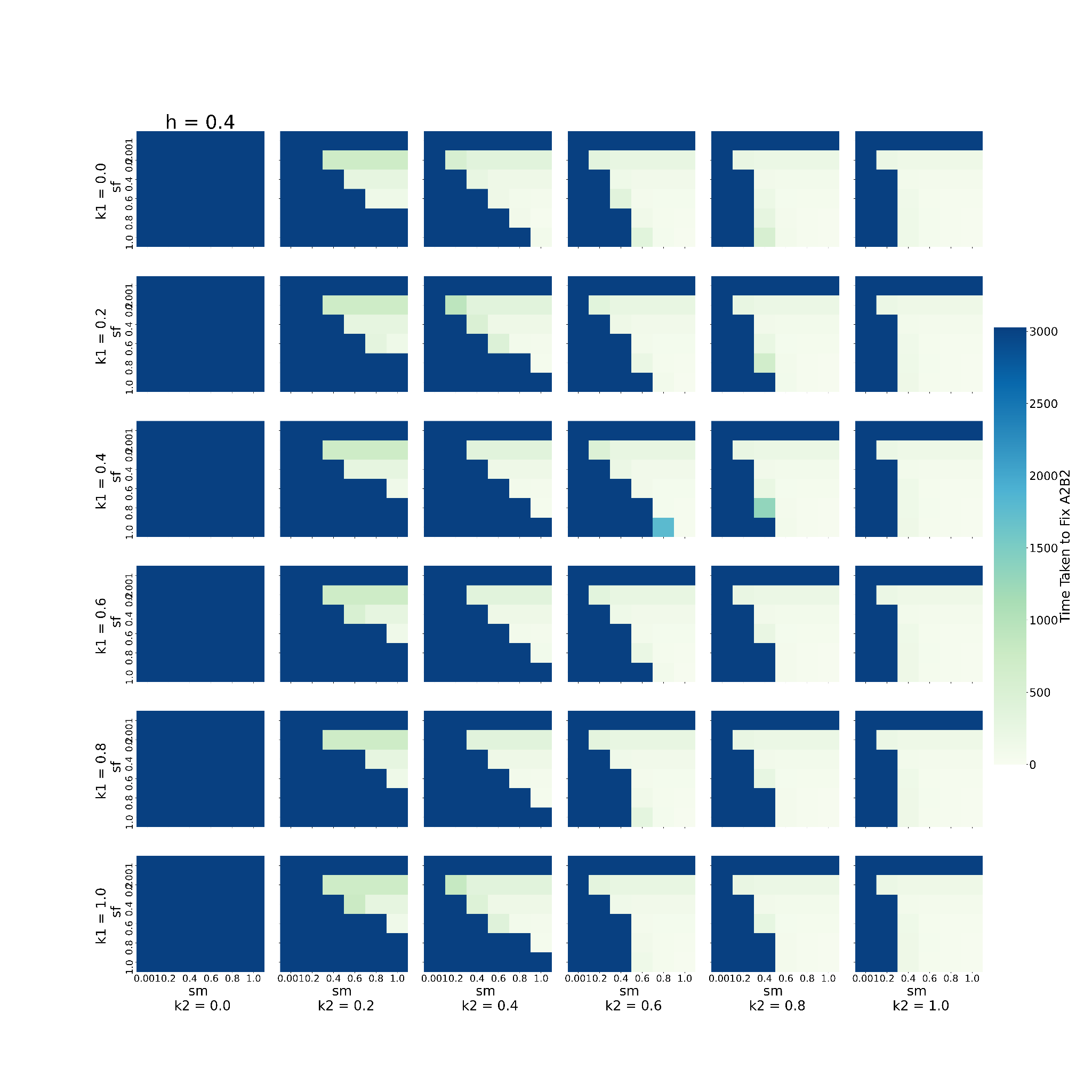

This graph shows the behaviour of the system when varying 2 of the 6 parameters, namely, sm, and sf, keeping h fixed to 0.4 (h= 0.4), given that the loci are X-linked. The remaining 3 parameters are kept constant at 0.2, except for D which is fixed to 0. Every block shows one combination of k1 and k2. k1 increases downwards, while k2 increases rightwards. The individual X axis for every k1-k2 denotes values of sm increasing *towards right* and the Y axis represents values of sf *increasing downwards*. Every box (sm-sf combination) is coloured to denote the time taken to fix A2B2 in the population. Bluer colours indicate the higher time taken to fix A2B2.

**Figure - S4.5**

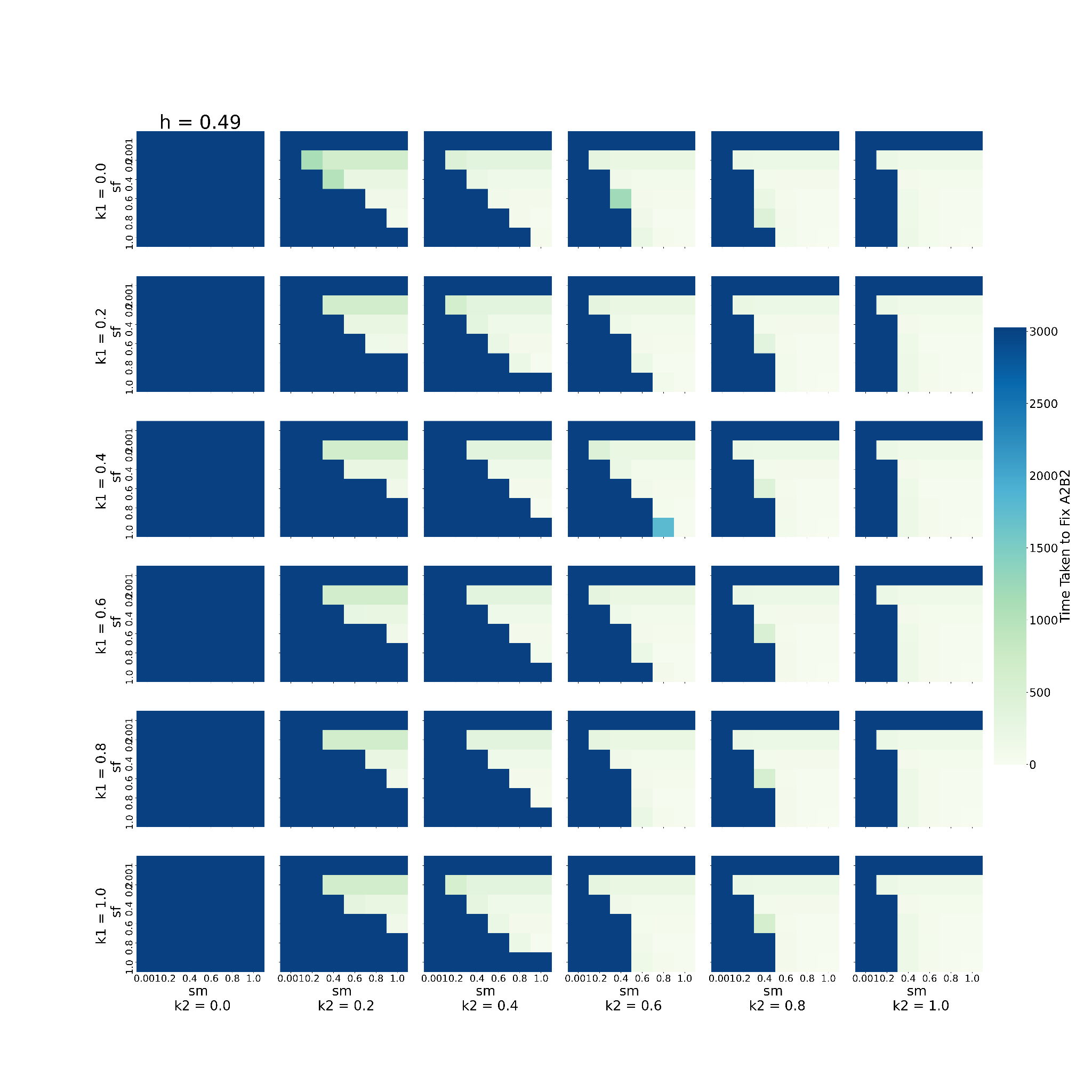

This graph shows the behaviour of the system when varying 2 of the 6 parameters, namely, sm, and sf, keeping h fixed to 0.49 (h= 0.49), given that the loci are X-linked. The remaining 3 parameters are kept constant at 0.2, except for D which is fixed to 0. Every block shows one combination of k1 and k2. k1 increases downwards, while k2 increases rightwards. The individual X axis for every k1-k2 denotes values of sm increasing *towards right* and the Y axis represents values of sf *increasing downwards*. Every box (sm-sf combination) is coloured to denote the time taken to fix A2B2 in the population. Bluer colours indicate the higher time taken to fix A2B2.

### Initial Linkage Disequilibrium and Recombination Rates

##### **The effect of varying Initial Linkage Disequilibrium and recombination rates on the frequency of A2B2 haplotype when the loci that are present on autosomes**

**Figure - S5.1**

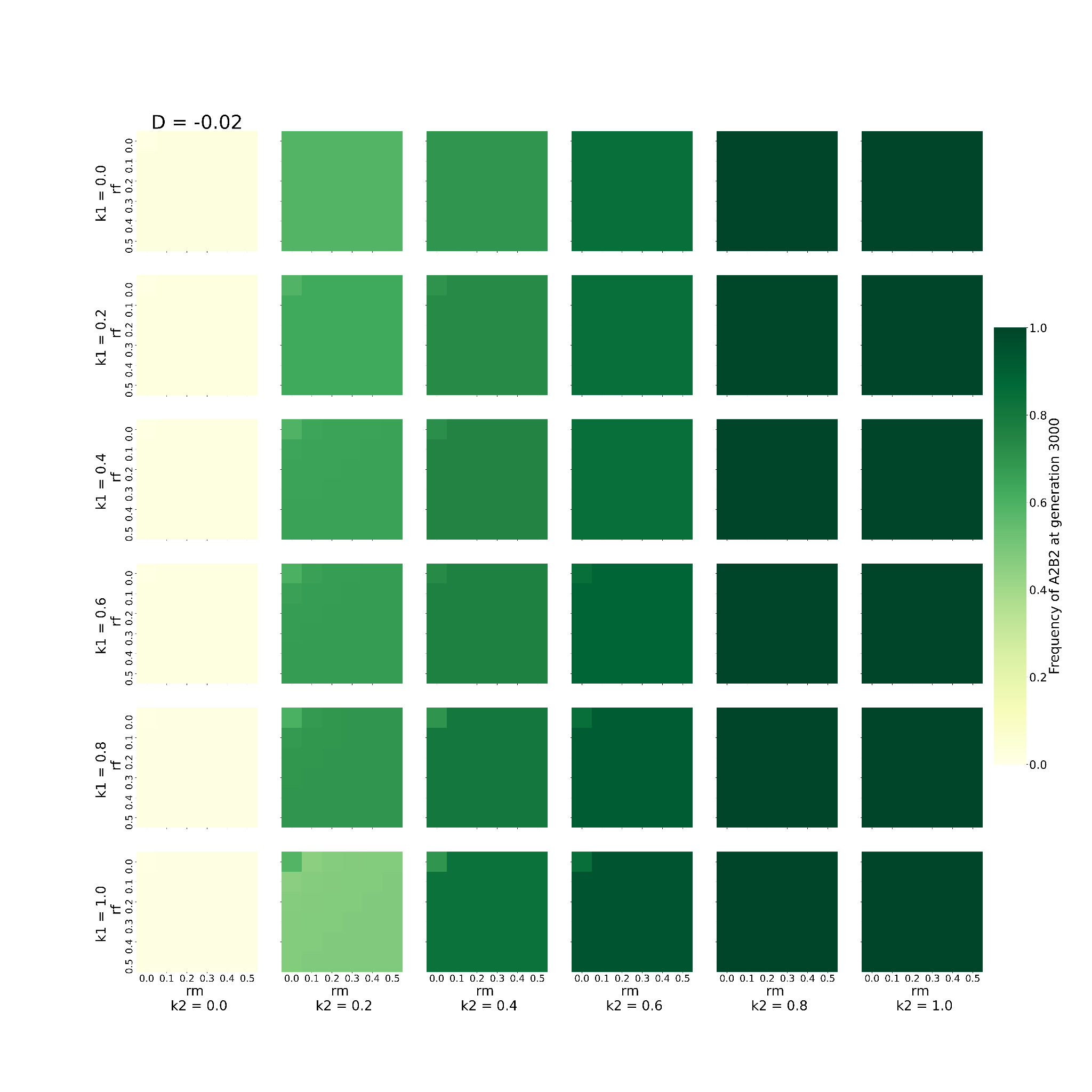

This graph shows the behaviour of the system when varying 2 of the 6 parameters, namely, rm, and rf, keeping D fixed to -0.02 (D= -0.02), given that the loci are present on the autosome. The remaining 3 parameters are kept constant at 0.2. Every block shows one combination of k1 and k2. k1 increases downwards, while k2 increases rightwards. The individual X axis for every k1-k2 denotes values of rm increasing *towards right* and the Y axis represents values of rf *increasing downwards*. Every box (rm-rf combination) is coloured to denote the frequency of A2B2 haplotype at 3000th generation. The greener the colour, the higher the frequency.

**Figure - S5.2**

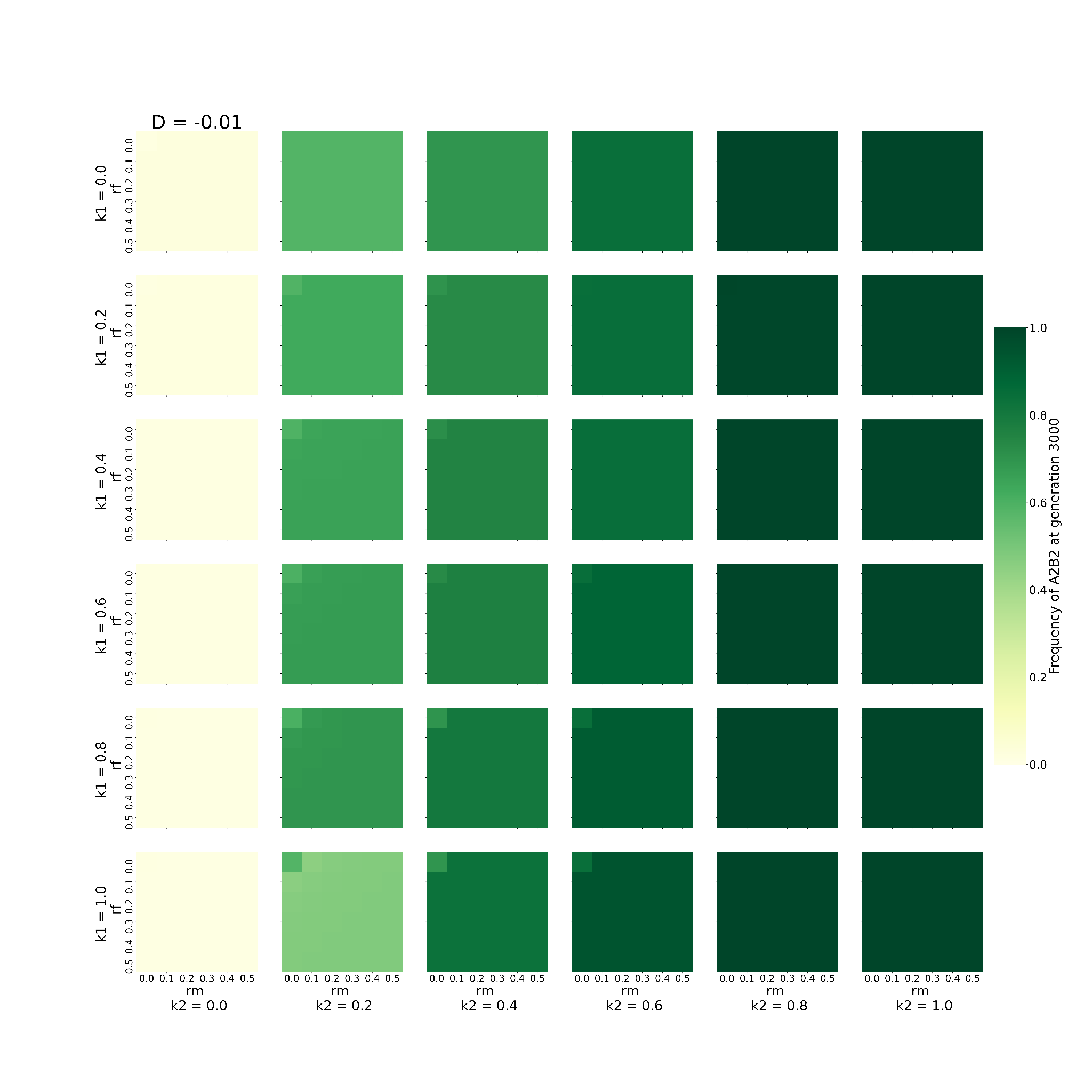

This graph shows the behaviour of the system when varying 2 of the 6 parameters, namely, rm, and rf, keeping D fixed to -0.01 (D= -0.01), given that the loci are present on the autosome. The remaining 3 parameters are kept constant at 0.2. Every block shows one combination of k1 and k2. k1 increases downwards, while k2 increases rightwards. The individual X axis for every k1-k2 denotes values of rm increasing *towards right* and the Y axis represents values of rf *increasing downwards*. Every box (rm-rf combination) is coloured to denote the frequency of A2B2 haplotype at 3000th generation. The greener the colour, the higher the frequency.

**Figure - S5.3**

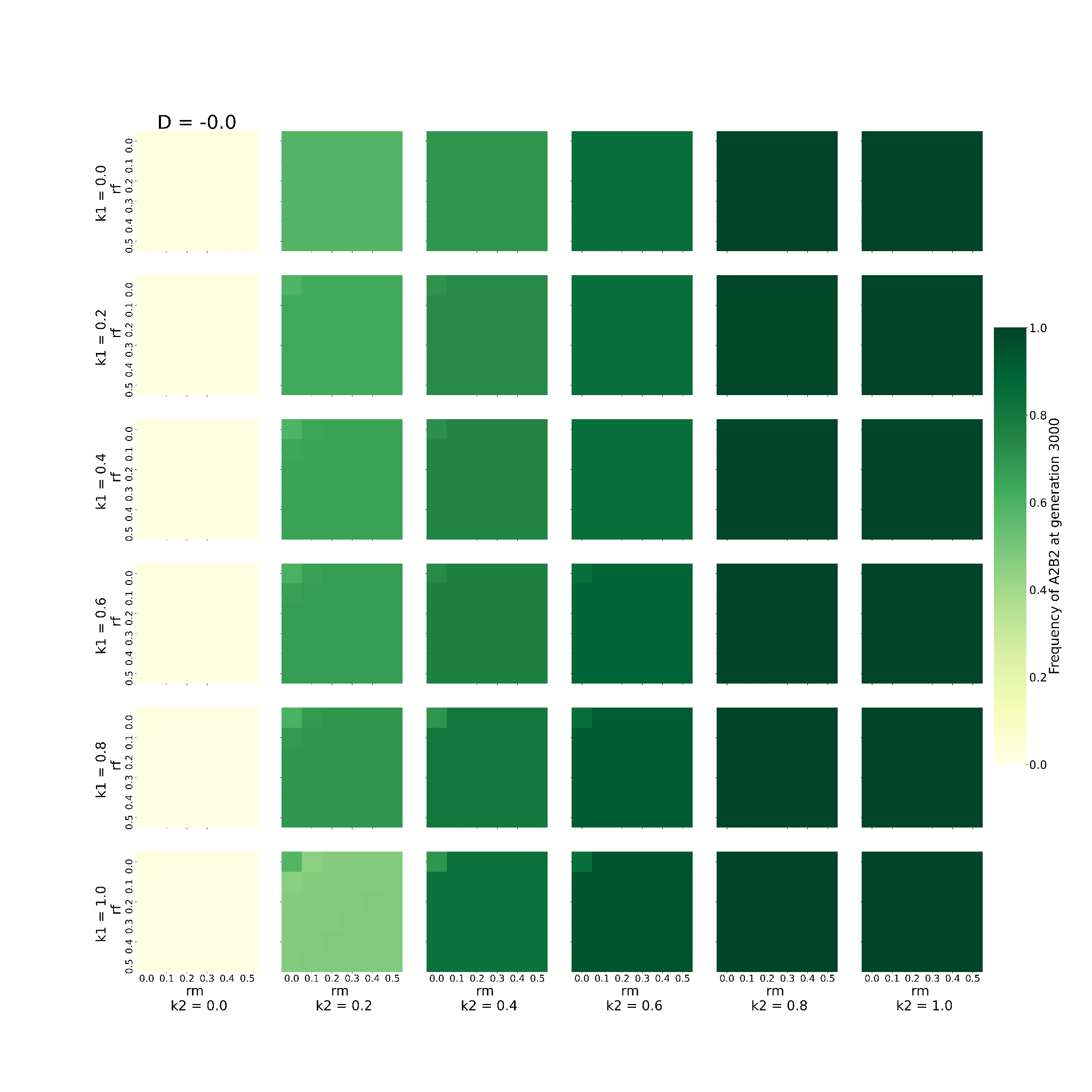

This graph shows the behaviour of the system when varying 2 of the 6 parameters, namely, rm, and rf, keeping D fixed to 0 (D= 0), given that the loci are present on the autosome. The remaining 3 parameters are kept constant at 0.2. Every block shows one combination of k1 and k2. k1 increases downwards, while k2 increases rightwards. The individual X axis for every k1-k2 denotes values of rm increasing *towards right* and the Y axis represents values of rf *increasing downwards*. Every box (rm-rf combination) is coloured to denote the frequency of A2B2 haplotype at 3000th generation. The greener the colour, the higher the frequency.

**Figure - S5.4**

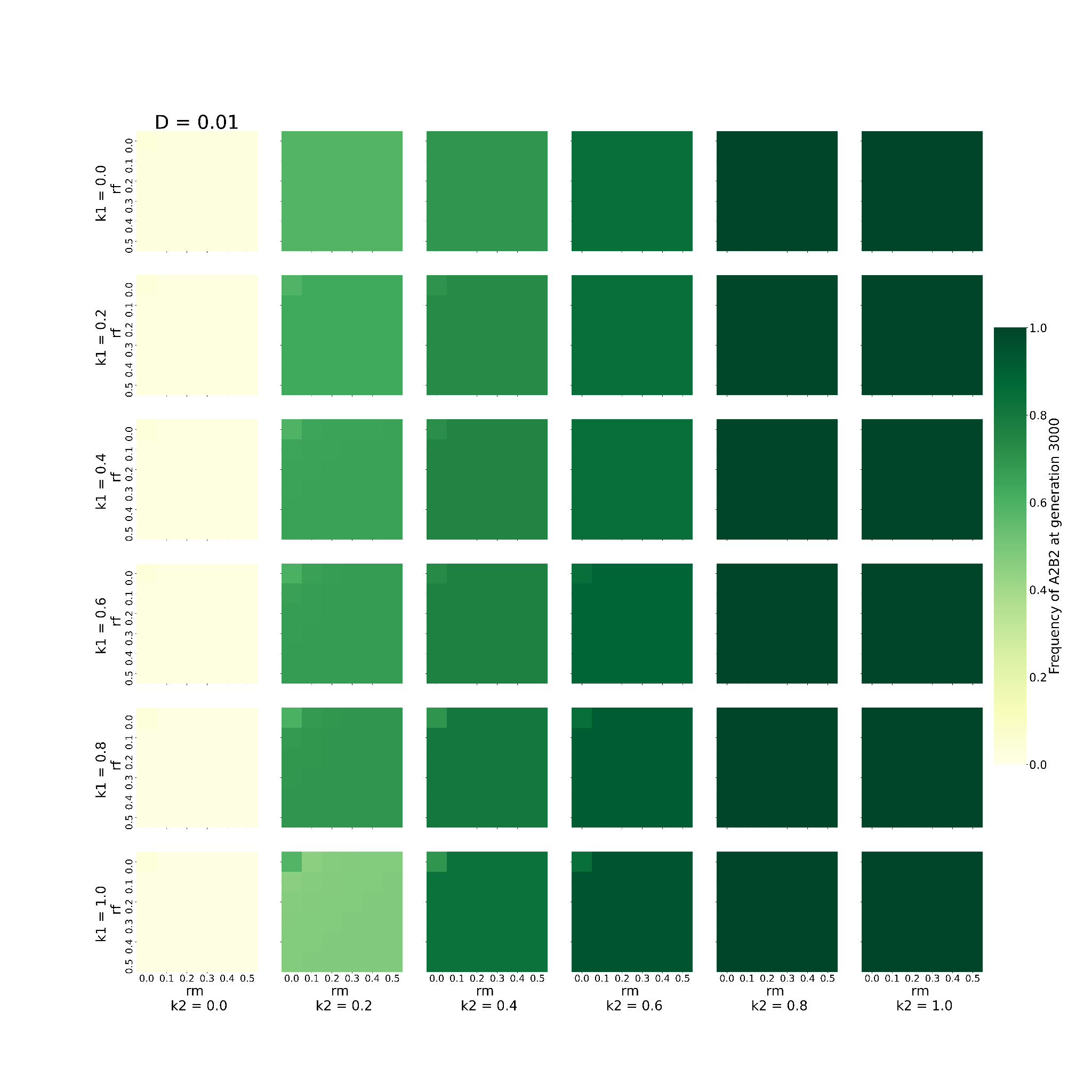

This graph shows the behaviour of the system when varying 2 of the 6 parameters, namely, rm, and rf, keeping D fixed to 0.01 (D= 0.01), given that the loci are present on the autosome. The remaining 3 parameters are kept constant at 0.2. Every block shows one combination of k1 and k2. k1 increases downwards, while k2 increases rightwards. The individual X axis for every k1-k2 denotes values of rm increasing *towards right* and the Y axis represents values of rf *increasing downwards*. Every box (rm-rf combination) is coloured to denote the frequency of A2B2 haplotype at 3000th generation. The greener the colour, the higher the frequency.

**Figure - S5.5**

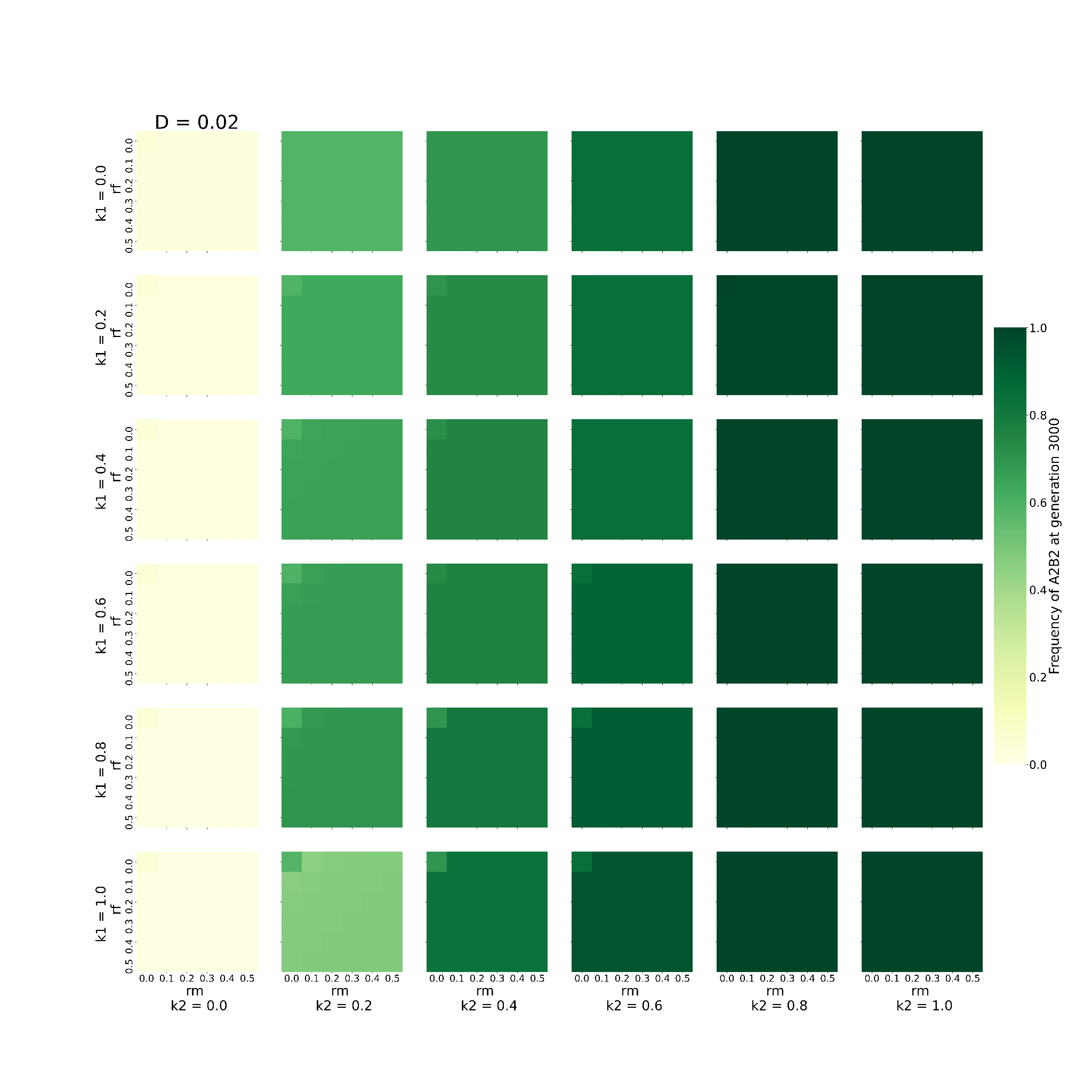

This graph shows the behaviour of the system when varying 2 of the 6 parameters, namely, rm, and rf, keeping D fixed to 0.02 (D= 0.02), given that the loci are present on the autosome. The remaining 3 parameters are kept constant at 0.2. Every block shows one combination of k1 and k2. k1 increases downwards, while k2 increases rightwards. The individual X axis for every k1-k2 denotes values of rm increasing *towards right* and the Y axis represents values of rf *increasing downwards*. Every box (rm-rf combination) is coloured to denote the frequency of A2B2 haplotype at 3000th generation. The greener the colour, the higher the frequency.

##### **The effect of varying Initial Linkage Disequilibrium and recombination rates on the time taken by A2B2 to get fixed when the loci are present on autosomes**

**Figure - S6.1**

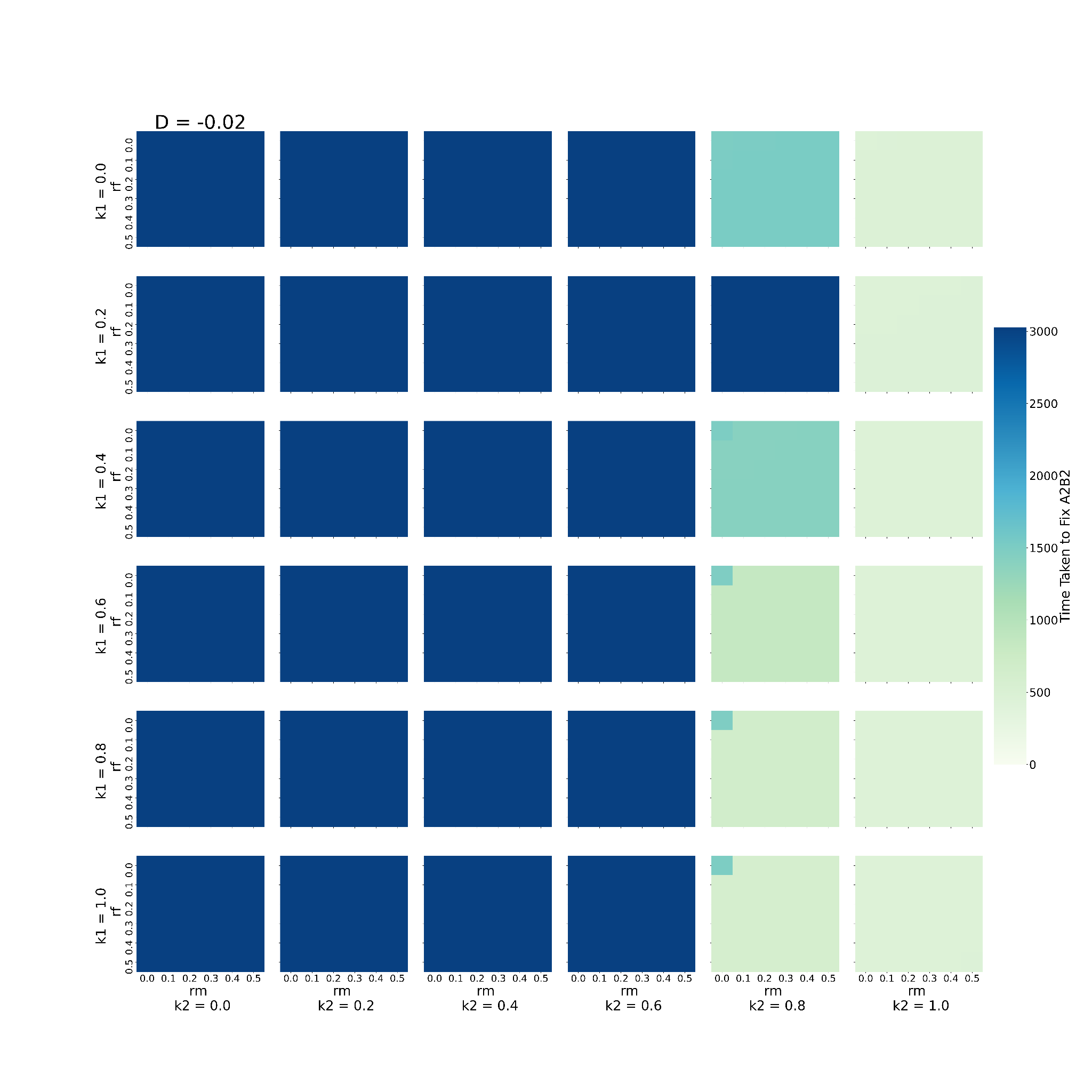

This graph shows the behaviour of the system when varying 2 of the 6 parameters, namely, rm, and rf, keeping D fixed to -0.02 (D= -0.02), given that the loci are present on the autosome. The remaining 3 parameters are kept constant at 0.2. Every block shows one combination of k1 and k2. k1 increases downwards, while k2 increases rightwards. The individual X axis for every k1-k2 denotes values of rm increasing *towards right* and the Y axis represents values of rf *increasing downwards*. Every box (rm-rf combination) is coloured to denote the time taken to fix A2B2 in the population. Bluer colours indicate the higher time taken to fix A2B2.

**Figure - S6.2**

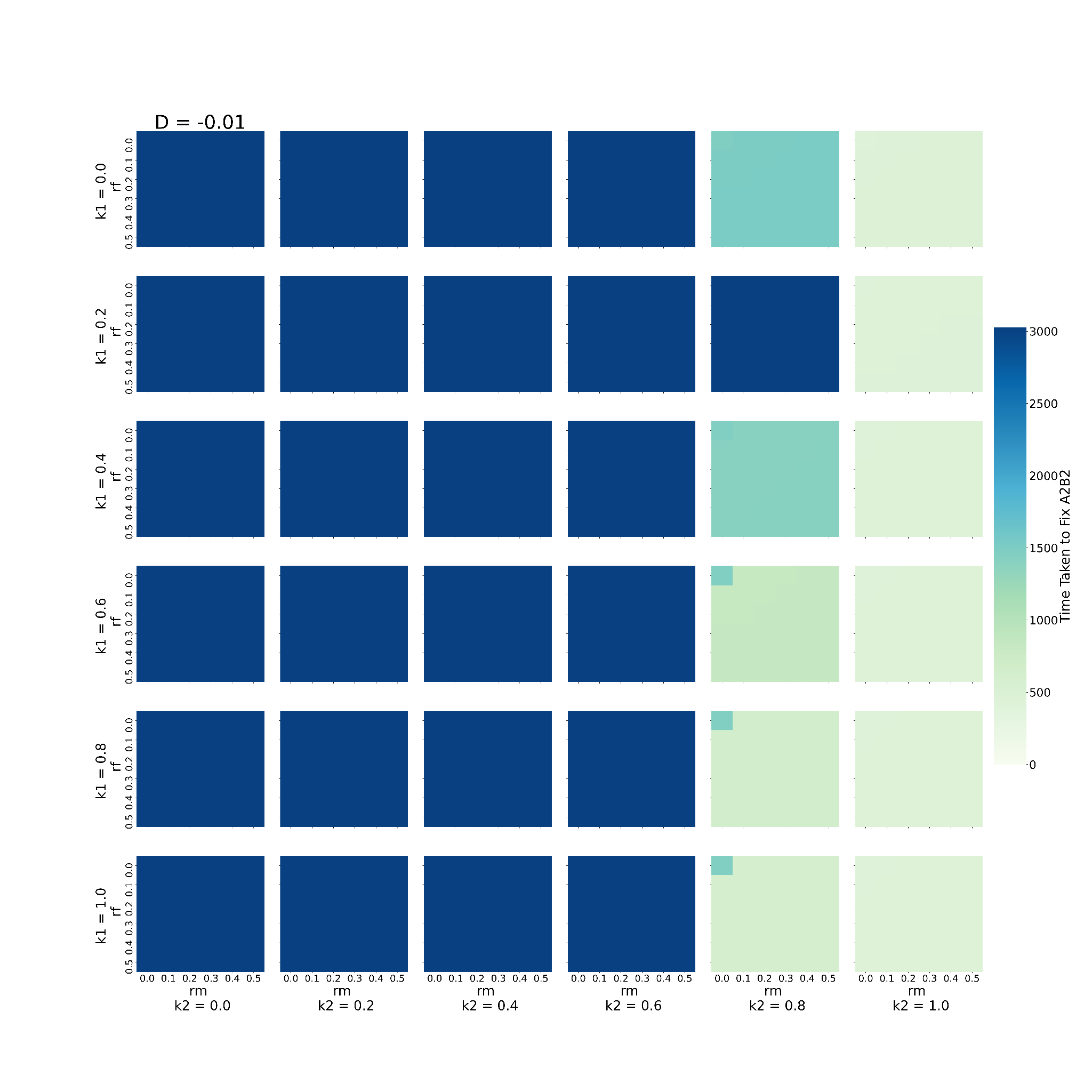

This graph shows the behaviour of the system when varying 2 of the 6 parameters, namely, rm, and rf, keeping D fixed to -0.01 (D= -0.01), given that the loci are present on the autosome. The remaining 3 parameters are kept constant at 0.2. Every block shows one combination of k1 and k2. k1 increases downwards, while k2 increases rightwards. The individual X axis for every k1-k2 denotes values of rm increasing *towards right* and the Y axis represents values of rf *increasing downwards*. Every box (rm-rf combination) is coloured to denote the time taken to fix A2B2 in the population. Bluer colours indicate the higher time taken to fix A2B2.

**Figure - S6.3**

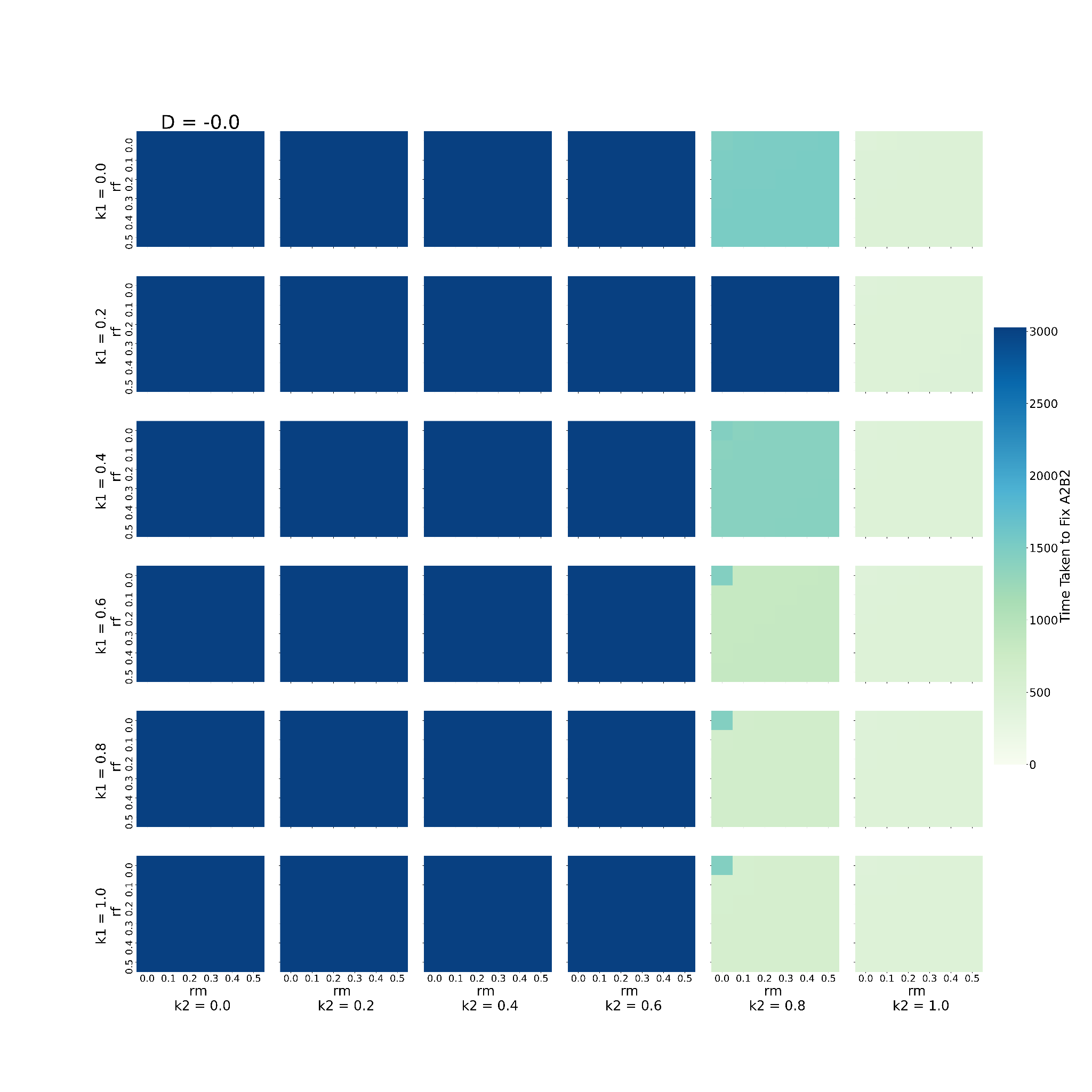

This graph shows the behaviour of the system when varying 2 of the 6 parameters, namely, rm, and rf, keeping D fixed to 0 (D= 0), given that the loci are present on the autosome. The remaining 3 parameters are kept constant at 0.2. Every block shows one combination of k1 and k2. k1 increases downwards, while k2 increases rightwards. The individual X axis for every k1-k2 denotes values of rm increasing *towards right* and the Y axis represents values of rf *increasing downwards*. Every box (rm-rf combination) is coloured to denote the time taken to fix A2B2 in the population. Bluer colours indicate the higher time taken to fix A2B2.

**Figure - S6.4**

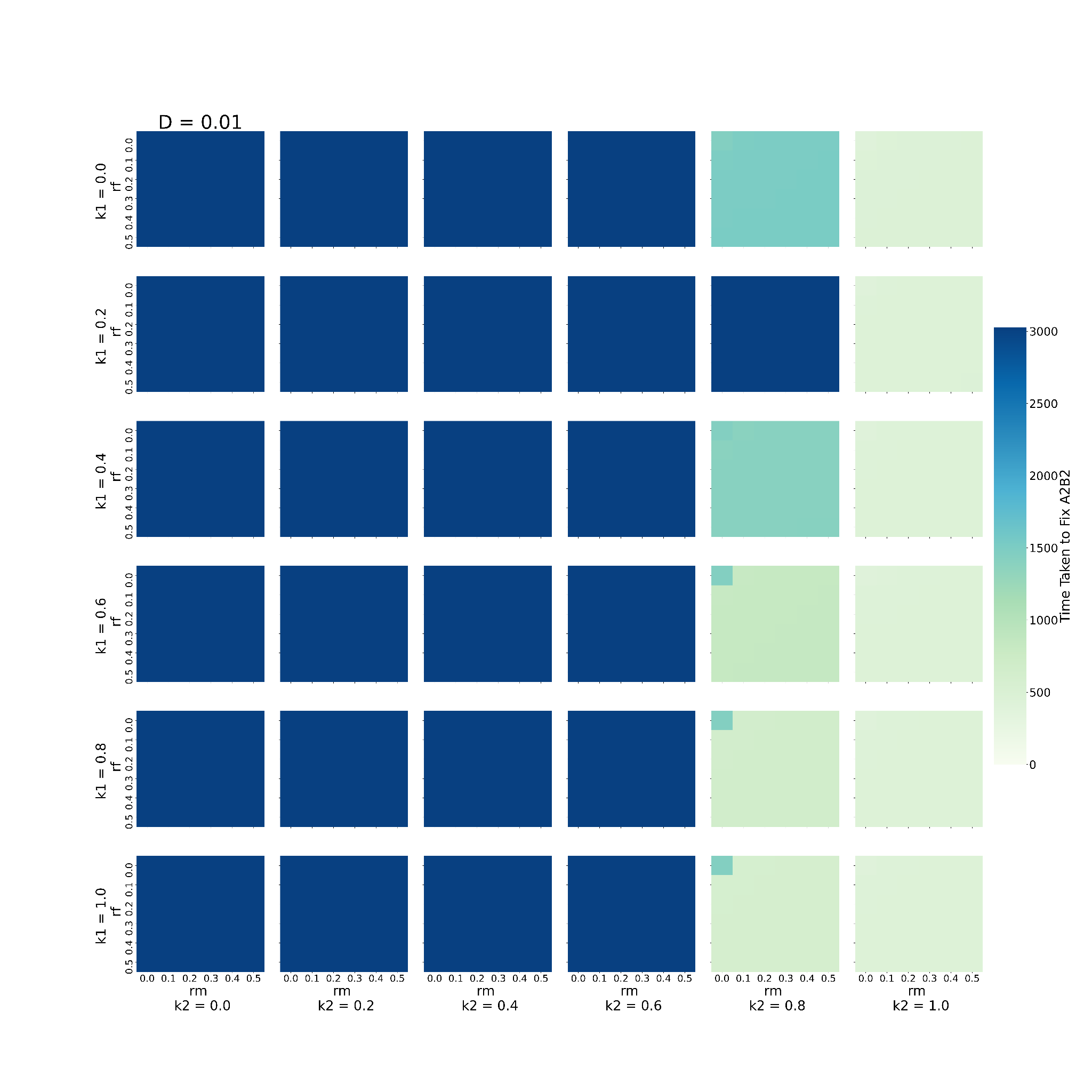

This graph shows the behaviour of the system when varying 2 of the 6 parameters, namely, rm, and rf, keeping D fixed to 0.01 (D= 0.01), given that the loci are present on the autosome. The remaining 3 parameters are kept constant at 0.2. Every block shows one combination of k1 and k2. k1 increases downwards, while k2 increases rightwards. The individual X axis for every k1-k2 denotes values of rm increasing *towards right* and the Y axis represents values of rf *increasing downwards*. Every box (rm-rf combination) is coloured to denote the time taken to fix A2B2 in the population. Bluer colours indicate the higher time taken to fix A2B2.

**Figure - S6.5**

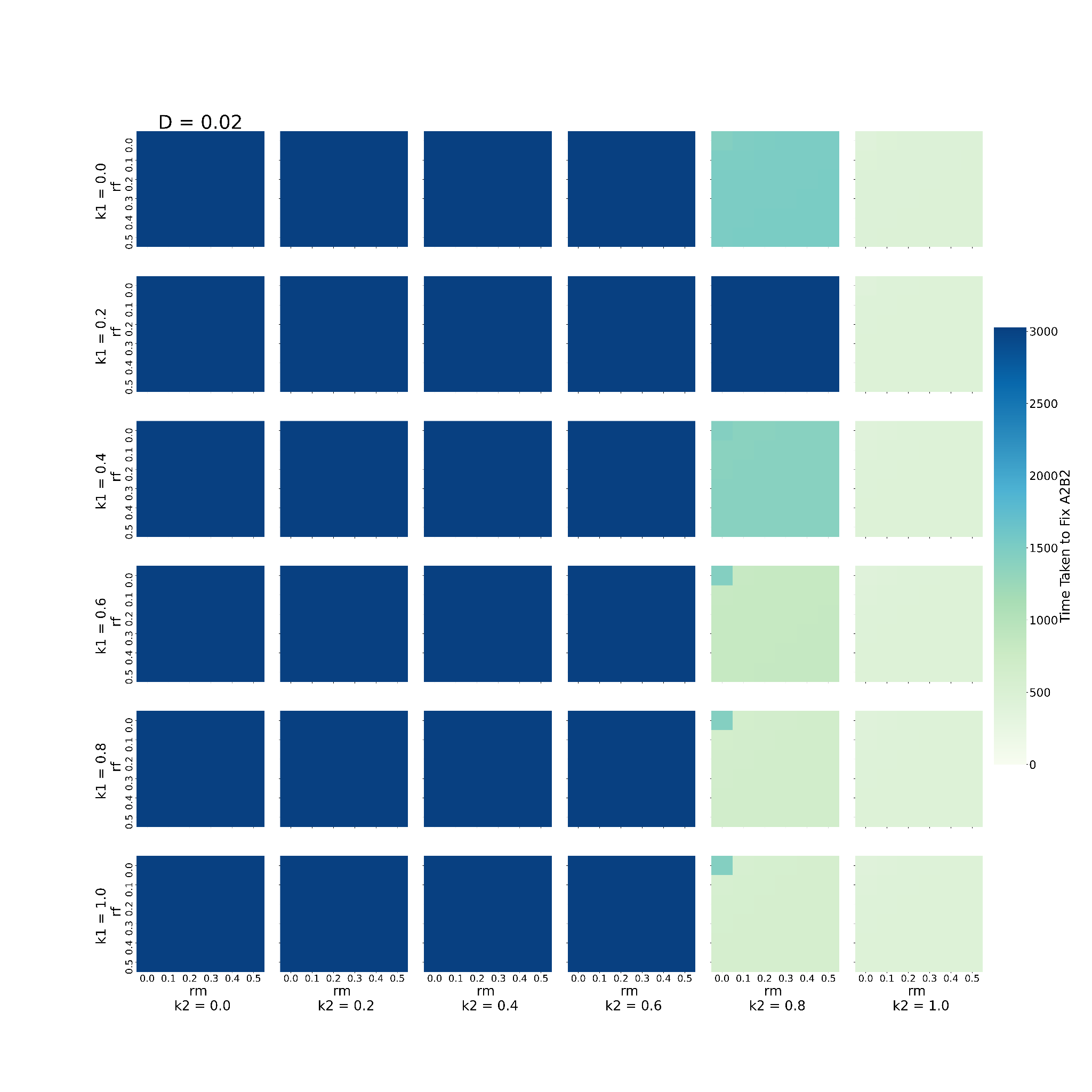

This graph shows the behaviour of the system when varying 2 of the 6 parameters, namely, rm, and rf, keeping D fixed to 0.02 (D= 0.02), given that the loci are present on the autosome. The remaining 3 parameters are kept constant at 0.2. Every block shows one combination of k1 and k2. k1 increases downwards, while k2 increases rightwards. The individual X axis for every k1-k2 denotes values of rm increasing *towards right* and the Y axis represents values of rf *increasing downwards*. Every box (rm-rf combination) is coloured to denote the time taken to fix A2B2 in the population. Bluer colours indicate the higher time taken to fix A2B2.

##### **The effect of varying recombination rates in females and initial linkage disequilibrium coefficient on the frequency of A2B2 haplotype when the loci are X-linked**

**Figure - S7**

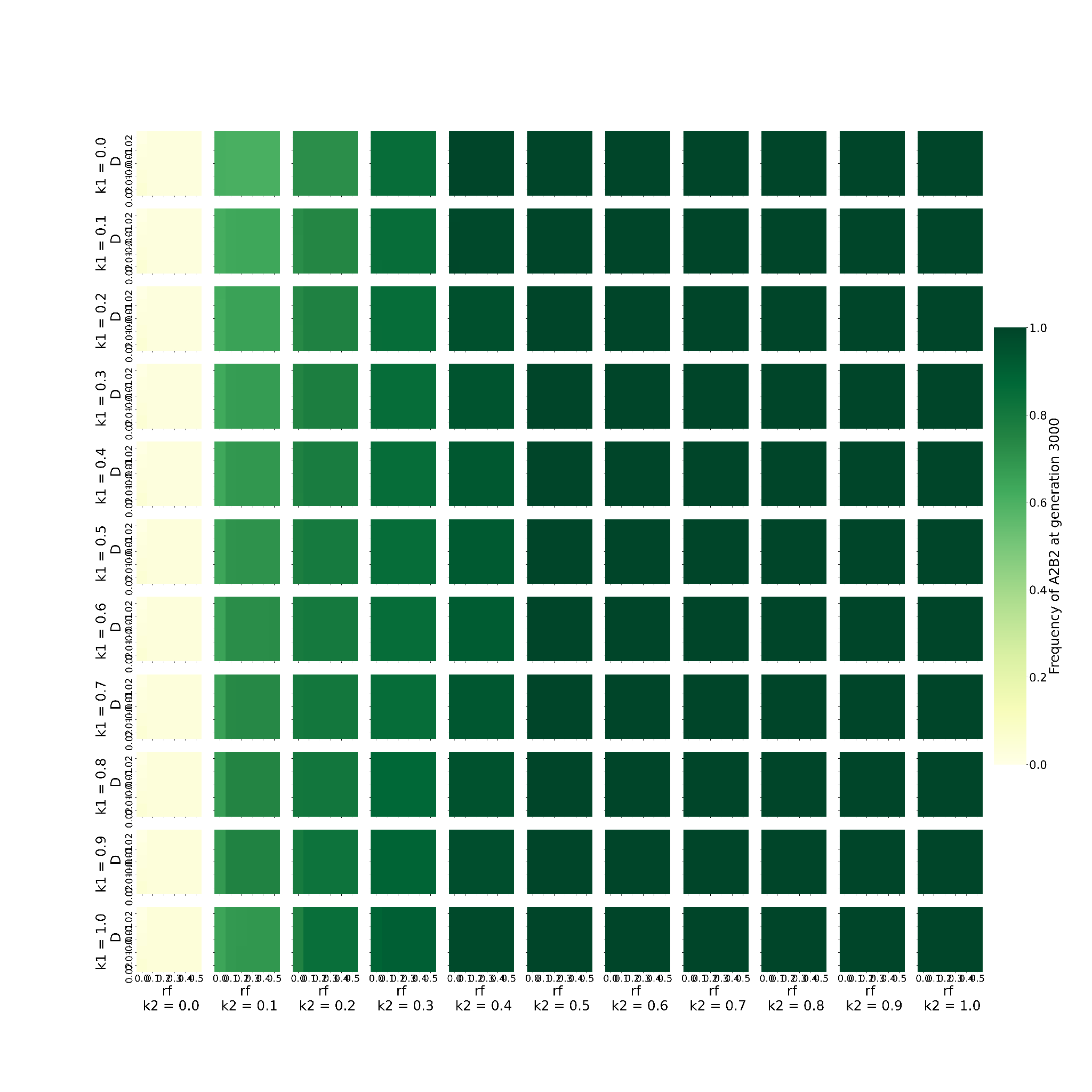

This graph shows the behaviour of the system when varying 2 of the 5 parameters, namely, rf, and D when the loci are X-linked. The remaining 3 parameters are kept constant at 0.2. Every block shows one combination of k1 and k2. k1 increases downwards, while k2 increases rightwards. The individual X axis for every k1-k2 denotes values of rf increasing *towards right* and the Y axis represents values of D *increasing downwards*. Every box (rf-D combination) is coloured to denote the frequency of A2B2 haplotype at 3000th generation. The greener the colour, the higher the frequency.

##### **The effect of varying recombination rates in females and initial linkage disequilibrium coefficient on the time taken by A2B2 haplotype to get fixed when the loci are X-linked**

**Figure - S8**

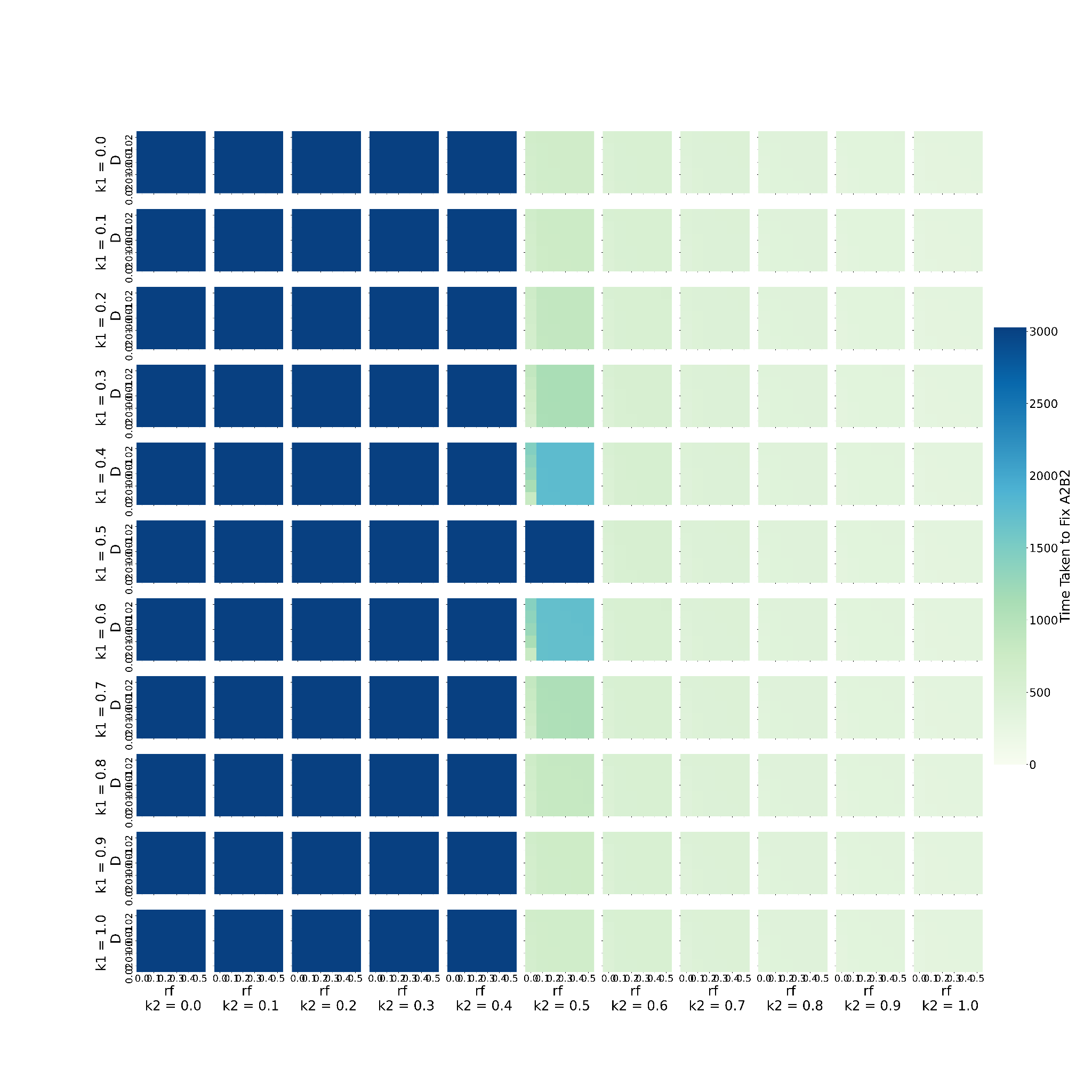

This graph shows the behaviour of the system when varying 2 of the 5 parameters, namely, rf, and D when the loci are X-linked. The remaining 3 parameters are kept constant at 0.2. Every block shows one combination of k1 and k2. k1 increases downwards, while k2 increases rightwards. The individual X axis for every k1-k2 denotes values of rf increasing *towards right* and the Y axis represents values of D *increasing downwards*. Every box (rf-D combination) is coloured to denote the time taken to fix A2B2 in the population. Bluer colours indicate the higher time taken to fix A2B2.

##### **Supplementary video 1 *‘A-h-Frequency.mp4’*** **:** This video shows the behaviour of the system when varying 3 of the 6 parameters, namely, h (dominance coefficient), sm (male selection coefficient) , and sf (female selection coefficient), given that the loci are present on the autosome. For every image in the video, h is fixed at a certain value (specified on top left). Every block shows one combination of k1 and k2. k1 increases downwards, while k2 increases rightwards. The individual X axis for every k1-k2 denotes values of sm *increasing towards right* and the Y axis represents values of sf *increasing downwards*. Every box (sm-sf combination) is coloured to denote the frequency of A2B2 haplotype at 3000th generation. The greener the colour, the higher the frequency. The video comprises graphs given in supplementary material (S1.1, S1.2,S1.3,S1.4,S1.5) and is composed of 5 loops of increasing h from 0.1 to 0.49

**Supplementary video 2 *‘X-h-Frequency.mp4’*:** This video shows the behaviour of the system when varying 3 of the 6 parameters, namely, h (dominance coefficient), sm (male selection coefficient), and sf (female selection coefficient), given that the loci are present on the X chromosome. For every image in the video, h is fixed at a certain value (specified on top left). Every block shows one combination of k1 and k2. k1 increases downwards, while k2 increases rightwards. The individual X axis for every k1-k2 denotes values of sm *increasing towards right* and the Y axis represents values of sf *increasing downwards*. Every box (sm-sf combination) is coloured to denote the frequency of A2B2 haplotype at 3000th generation. The greener the colour, the higher the frequency. The video comprises graphs given in supplementary material (S3.1, S3.2,S3.3,S3.4,S3.5) and is composed of 5 loops of increasing h from 0.1 to 0.49.

**Supplementary video 3 *‘A-h-Time.mp4’*:** This video shows the behaviour of the system when varying 3 of the 6 parameters, namely, h (dominance coefficient), sm (male selection coefficient), and sf (female selection coefficient), given that the loci are present on the autosome. For every image in the video, h is fixed at a certain value (specified on top left). Every block shows one combination of k1 and k2. k1 increases downwards, while k2 increases rightwards. The individual X axis for every k1-k2 denotes values of sm *increasing towards right* and the Y axis represents values of sf *increasing downwards*. Every box (sm-sf combination) is coloured to denote the time taken to fix A2B2 in the population. Bluer colours indicate the higher time taken to fix A2B2. The bluest colour indicates that the haplotype did not get fixed in 3000 generations. The video comprises graphs given in supplementary material (S2.1, S2.2,S2.3,S2.4,S2.5) and is composed of 5 loops of increasing h from 0.1 to 0.49.

**Supplementary video 4 *‘X-h-Time.mp4’*:** The video shows the behaviour of the system when varying 3 of the 6 parameters, namely, h (dominance coefficient), sm (male selection coefficient), and sf (female selection coefficient), given that the loci are present on the X chromosome. For every image in the video, h is fixed at a certain value (specified on top left). Every block shows one combination of k1 and k2. k1 increases downwards, while k2 increases rightwards. The individual X axis for every k1-k2 denotes values of sm *increasing towards right* and the Y axis represents values of sf *increasing downwards*. Every box (sm-sf combination) is coloured to denote the time taken to fix A2B2 in the population. Bluer colours indicate the higher time taken to fix A2B2. The bluest colour indicates that the haplotype did not get fixed in 3000 generations. The video comprises graphs given in supplementary material (S4.1, S4.2,S4.3,S4.4,S4.5) and is composed of 5 loops of increasing h from 0.1 to 0.49.

**Supplementary video 5 *‘A-D-Frequency.mp4’*:** The video shows the behaviour of the system when varying 3 of the 6 parameters, namely, D (initial linkage disequilibrium), rm (male recombination rate) and rf (female recombination rate) given that the loci are present on the autosome. For every image in the video, D is kept fixed to a certain value (specified on top left). Every block shows one combination of k1 and k2. k1 increases downwards, while k2 increases rightwards. The individual X axis for every k1-k2 denotes values of rm *increasing towards right* and the Y axis represents values of rf *increasing downwards*. Every box (rm-rf combination) is coloured to denote the frequency of A2B2 haplotype at 3000th generation. The greener the colour, the higher the frequency. The video comprises graphs given in supplementary material (S5.1, S5.2,S5.3,S5.4,S5.5) and is composed of 5 loops of increasing D from -0.02 to 0.02.

**Supplementary video 6 *‘A-D-Time.mp4’*:** The video shows the behaviour of the system when varying 3 of the 6 parameters, namely, D (initial linkage disequilibrium), rm (male recombination rate) and rf (female recombination rate) given that the loci are present on the autosome. For every image in the video, D is kept fixed to a certain value (specified on top left). Every block shows one combination of k1 and k2. k1 increases downwards, while k2 increases rightwards. The individual X axis for every k1-k2 denotes values of rm *increasing towards right* and the Y axis represents values of rf *increasing downwards*. Every box (rm-rf combination) is coloured to denote the time taken to fix A2B2 in the population. Bluer colours indicate the higher time taken to fix A2B2. The bluest colour indicates that the haplotype did not get fixed in 3000 generations. The video comprises graphs given in supplementary material (S4.1, S6.2,S6.3,S6.4,S6.5) andis composed of 5 loops of increasing D from -0.02 to 0.02.

### Codes used for simulating the system:

**Code 1 ‘PSAcommented.py’: This code was used to look at the entire 8 dimensional parameter space for loci on the autosomes, and to calculate the fraction of parameter space where IaSC gets resolved and average time taken to resolve it for every k1-k2. The data generated from this code was used in constructing Figure 1 and Figure 2.**

#importing libraries

import numpy as np

import matplotlib.pyplot as plt

import seaborn as sns

from random import uniform

import openpyxl as op

from numpy import mean, std

Checklist=np.arange(1-0.020,1+0.020,0.001) # a list to check if all haplotype frequencies add up to approximately 1

checklist=[round(i,3) for i in Checklist]

T=3000

SF=np.arange(0.2,1.1,0.1)

Sf=[0.001]+[round(x,1) for x in SF] # range of values for female selection coefficient

HD=np.arange(0.1,0.5,0.1)

H=[round(x,1) for x in HD]+[0.49] # range of values for dominance coefficient

RF = np.arange(0.1,0.5,0.1)

Rf= [round(x,1) for x in RF] # range of values for female recombination rates

Rm=Rf # range of values for male recombination rates

Krange=np.arange(0,1.01,0.1)

krange=[round(k,2) for k in Krange] # range of values for k1 and k2 (effect of B2 on A1 and A2)

for k1 in krange: # iterating over k1

for k2 in krange: # iterating over k2

#number of fixations achieved for each allele/haplotype

A2f_fix=0

B2f_fix=0

A2B2f_fix=0

A2m_fix=0

B2m_fix=0

A2B2m_fix=0

\# time taken to achieve fixation

A2f_t=[]

B2f_t=[]

A2B2f_t=[]

A2m_t=[]

B2m_t=[]

A2B2m_t=[]

total=0 # keeps count of the total number of cases explored for each k1-k2

for h in H: # iterating over values of dominance coefficients

for sf in Sf: # iterating over values of female selection coefficient

SM=np.arange(max(0,min(1,sf*h/(1 - h*(1-sf)))),max(0,min(1,sf*(1-h)/h*(1-sf)))+0.00001,0.1) # determining range of male selection coefficients based on Connallon and Clark (2010)

Sm=[round(x,2) for x in SM]

for sm in Sm: # iterating over values of male selection coefficients

#fitness scheme

f11= 1

f13= 1-k1*sf

f31= 1-sf

f33= 1-(1-k2)*sf

f12= 1-k1*h*sf

f32= 1 - (1+h - h*(1-k2)-k2)*sf

f21= 1- h*sf

f23= 1 - abs((1-k1-k2)*h*sf)

f22r= 1- abs(h*sf*(1-k1))

f22c= 1-(1-k2)*sf*h

m11= 1-sm

m21= 1 - h*sm

m33= 1

m12=1 - sm

m13=1 - sm

m22r=1 - h*sm

m22c=1 - h*sm

m23=1 - h*sm

m31=1

m32=1

A2f=max(0.01,min(0.99,(sm*(1-h)-sf*h)/((sm+sf)*(1-2*h)))) # setting initial frequency of A2 allele in females

A1f=1-A2f # setting initial frequency of A1 allele in females

A2m=max(0.01,min(0.99,(sm*(1-h)-sf*h)/((sm+sf)*(1-2*h)))) # setting initial frequency of A2 allele in males

A1m=1-A2m # setting initial frequency of A1 allele in females

B1=0.95 # setting initial frequency of B1 and B2 alleles in population

B2=0.05

\# calculating initial frequencies of each haplotype

x1=A1f*B1

x2=A1f*B2

x3=A2f*B1

x4=A2f*B2

y1=A1m*B1

y2=A1m*B2

y3=A2m*B1

y4=A2m*B2

possible_D=[0.25,1-x1,x1-0,1-x2,x2-0,1-x3,x3-0,1-x4,x4-0,1-y1,y1-0,1-y2,y2-0,1-y3,y3-0,1-y4,y4-0] # determining the maximum possible value of linkage disequilibrium (D) based on initial frequencies

Drange=np.arange(-min(possible_D),min(possible_D),0.01) # setting the range of values for D

for D in Drange: # iterating over values of linkage disequilibrium

for rf in Rf: # iterating over values of female recombination rates

for rm in Rm: # iterating over the values of male recombination rates

\# generating the time series

\# calculating initial frequencies of each haplotype

x1=A1f*B1

x2=A1f*B2

x3=A2f*B1

x4=A2f*B2

y1=A1m*B1

y2=A1m*B2

y3=A2m*B1

y4=A2m*B2

x1+=D

x2+=-D

x3+=-D

x4+=D

y1+=D

y2+=-D

y3+=-D

y4+=D

\# time series

X1=[x1]

X2=[x2]

X3=[x3]

X4=[x4]

Y1=[y1]

Y2=[y2]

Y3=[y3]

Y4=[y4]

\# keeping track of whether alleles or haplotypes are fixed

A2fB=False

B2fB=False

A2B2fB=False

A2mB=False

B2mB=False

A2B2mB=False

total+=1 # incrementing the total count of cases covered by 1

for t in range(T):

\# calculating haplotype frequencies in (t+1)^th generation

wm = x1*y1*m11 + (x1*y2+x2*y1)*m12 + x2*y2*m13 + (x1*y3+x3*y1)*m21 + (x2*y3+x3*y2)*m22r + (x1*y4+x4*y1)*m22c + (x2*y4+x4*y2)*m23 + x3*y3*m31 + (x3*y4+x4*y3)*m32 + x4*y4*m33

wf = x1*y1*f11 + (x1*y2+x2*y1)*f12 + x2*y2*f13 + (x1*y3+x3*y1)*f21 + (x2*y3+x3*y2)*f22r + (x1*y4+x4*y1)*f22c + (x2*y4+x4*y2)*f23 + x3*y3*f31 + (x3*y4+x4*y3)*f32 + x4*y4*f33

Y1+=[(2*x1*y1*m11 + (x1*y2+x2*y1)*m12 + (x1*y3+x3*y1)*m21 + (1-rm)*m22c*(x1*y4+x4*y1) + rm*m22r*(x2*y3+x3*y2))/(2*wm)]

Y2+=[(2*x2*y2*m13 + (x1*y2+x2*y1)*m12 + (x2*y4+x4*y2)*m23 + (1-rm)*m22r*(x3*y2+x2*y3) + rm*m22c*(x4*y1+x1*y4))/(2*wm)]

Y3+=[(2*x3*y3*m31 + (x1*y3+x3*y1)*m21 + (x3*y4+x4*y3)*m32 + (1-rm)*m22r*(x2*y3+x3*y2) + rm*m22c*(x4*y1+x1*y4))/(2*wm)]

Y4+=[(2*x4*y4*m33 + (x2*y4+x4*y2)*m23 + (x4*y3+x3*y4)*m32 + (1-rm)*m22c*(x4*y1+x1*y4) + rm*m22r*(x2*y3+x3*y2))/(2*wm)]

X1+=[(2*x1*y1*f11 + (x1*y2+x2*y1)*f12 + (x1*y3+x3*y1)*f21 + (1-rf)*f22c*(x1*y4+x4*y1) + rf*f22r*(x2*y3+x3*y2))/(2*wf)]

X2+=[(2*x2*y2*f13 + (x1*y2+x2*y1)*f12 + (x2*y4+x4*y2)*f23 + (1-rf)*f22r*(x3*y2+x2*y3) + rf*f22c*(x4*y1+x1*y4))/(2*wf)]

X3+=[(2*x3*y3*f31 + (x1*y3+x3*y1)*f21 + (x3*y4+x4*y3)*f32 + (1-rf)*f22r*(x2*y3+x3*y2) + rf*f22c*(x4*y1+x1*y4))/(2*wf)]

X4+=[(2*x4*y4*f33 + (x2*y4+x4*y2)*f23 + (x4*y3+x3*y4)*f32 + (1-rf)*f22c*(x4*y1+x1*y4) + rf*f22r*(x2*y3+x3*y2))/(2*wf)]

x1=X1[-1]

x2=X2[-1]

x3=X3[-1]

x4=X4[-1]

y1=Y1[-1]

y2=Y2[-1]

y3=Y3[-1]

y4=Y4[-1]

\# safety checks for negative frequencies

if x1<0 or x2<0 or x3<0 or x4<0:

print('WRONG X',x1,x2,x3,x4)

print('k1,k2,h,sm,sf,wf, D, rf, minD-',k1,k2,h,sm,sf,wf,D,rf,min(possible_D))

break

if y1<0 or y2<0 or y3<0 or y4<0:

print('WRONG Y',y1,y2,y3,y4)

print('k1,k2,h,sm,sf,wm, D, rf, minD-',k1,k2,h,sm,sf,wm,D,rf,min(possible_D))

break

if round((x3+x4),3)==1 and A2fB==False: # checking for A2 fixation in females

A2f_fix+=1

A2f_t+=[t]

A2fB=True

if round((x2+x4),3)==1 and B2fB==False: #checking for B2 fixation in females

B2f_fix+=1

B2f_t+=[t]

B2fB=True

if round(x4,3)==1 and A2B2fB==False: # checking for A2B2 fixation in females

A2B2f_fix+=1

A2B2f_t+=[t]

A2B2fB=True

if round((y3+y4),3)==1 and A2mB==False: # checking for A2 fixation in males

A2m_fix+=1

A2m_t+=[t]

A2mB=True

if round((y2+y4),3)==1 and B2mB==False: # checking for B2 fixation in males

B2m_fix+=1

B2m_t+=[t]

B2mB=True

if round(y4,3)==1 and A2B2mB==False: # checking for A2B2 fixation in males

A2B2m_fix+=1

A2B2m_t+=[t]

A2B2mB=True

#check for all haplotype frequencies adding upto 1

if round(x1+x2+x3+x4,3) not in checklist:

print(f'x not summing upto 1-{x1+x2+x3+x4}',x1,x2,x3,x4,t)

break

if round(y1+y2+y3+y4,3) not in checklist:

print(f'y not summing upto 1-{y1+y2+y3+y4}',y1,y2,y3,y4,t)

break

data=[round(A2f_fix/total,3),round(B2f_fix/total,3),round(A2B2f_fix/total,3),round(A2m_fix/total,3),round(B2m_fix/total,3),round(A2B2m_fix/total,3)] #rounding off values of fraction of conflict resolution to 3 decimal places

Worksheet=op.load_workbook('ParameterSpace fixationA.xlsx') #opening excel sheet to store the data collected

\# initializing sheets to store data for individual alleles/haplotypes

A2F=Worksheet['A2 female']

B2F=Worksheet['B2 female']

A2B2F=Worksheet['A2B2 female']

A2M=Worksheet['A2 male']

B2M=Worksheet['B2 male']

A2B2M=Worksheet['A2B2 male']

sheets=[A2F,B2F,A2B2F,A2M,B2M,A2B2M]

\# setting initial row and column values

ROW=0

COLUMN=0

#print('k1,k2',k1,k2)

for i in range(2,13,1): #iterating over rows in the sheet

if float(A2F.cell(row=i,column=1).value)==k1: # finding the row corresponding to k1

for j in range(2,13,1): #iterating over columns in the sheet

if float(A2F.cell(row=1,column=j).value)==k2: #finding the column corresponding to k2

ROW=i

COLUMN=j

#print(ROW,COLUMN)

for i in range(6):

sheets[i].cell(ROW,COLUMN).value=data[i] # adding data to the correct cell in the appropriate sheet

Worksheet.save('ParameterSpace fixationA.xlsx') # saving file

\# setting variables to None for reuse

Worksheet=None

A2F=None

B2F=None

A2B2F=None

A2M=None

B2M=None

A2B2M=None

listS=[A2f_t,B2f_t,A2B2f_t,A2m_t,B2m_t,A2B2m_t] # list of lists of time taken to fix the allele/haplotype

for x in listS:

if x==[]: # setting value to T+25 if no fixation was achieved

x+=[T+25]

data=[round(mean(x),3) for x in listS] # calculating mean time taken to fix the allele/haplotype

Worksheet=op.load_workbook('ParameterSpace timeA.xlsx') # loading time Workbook

\# initializing sheets

A2F=Worksheet['A2 female']

B2F=Worksheet['B2 female']

A2B2F=Worksheet['A2B2 female']

A2M=Worksheet['A2 male']

B2M=Worksheet['B2 male']

A2B2M=Worksheet['A2B2 male']

sheets=[A2F,B2F,A2B2F,A2M,B2M,A2B2M]

#print('k1,k2',k1,k2)

for i in range(2,13,1): # finding correct row and column to enter the data into

if float(A2F.cell(row=i,column=1).value)==k1:

for j in range(2,13,1):

if float(A2F.cell(row=1,column=j).value)==k2:

ROW=i

COLUMN=j

for i in range(6):

sheets[i].cell(ROW,COLUMN).value=data[i] # entering data

print (data[i])

Worksheet.save('ParameterSpace timeA.xlsx') # saving Workbook

\# setting variables to None for reuse

Worksheet=None

A2F=None

B2F=None

A2B2F=None

A2M=None

B2M=None

A2B2M=None

**Code 2 ‘PSXcommented.py’: This code was used to look at the entire 8 dimensional parameter space for X-linked loci, and to calculate the fraction of parameter space where IaSC gets resolved and average time taken to resolve it for every k1-k2. The data generated from this code was used in constructing Figure 1 and Figure 2.**

#import libraries

import numpy as np

import matplotlib.pyplot as plt

import seaborn as sns

from random import uniform

import openpyxl as op

from numpy import mean, std

Checklist=np.arange(1-0.020,1+0.020,0.001)# a list to check if all haplotype frequencies add up to approximately 1

checklist=[round(i,3) for i in Checklist]

T=3000

SF=np.arange(0.2,1.1,0.1)

Sf=[0.001]+[round(x,1) for x in SF] # range of values for female selection coefficient

HD=np.arange(0.1,0.5,0.1)

H=[round(x,1) for x in HD]+[0.49] # range of values for dominance coefficient

RF = np.arange(0.1,0.5,0.1)

Rf= [round(x,1) for x in RF] # range of values for female recombination rates

Krange=np.arange(0,1.01,0.1)

krange=[round(k,1) for k in Krange] # range of values for k1 and k2 (effect of B2 on A1 and A2)

for k1 in krange: # iterating over k1

for k2 in krange: # iterating over k2

#number of fixations achieved for each allele/haplotype

A2f_fix=0

B2f_fix=0

A2B2f_fix=0

A2m_fix=0

B2m_fix=0

A2B2m_fix=0

\# time taken to achieve fixation

A2f_t=[]

B2f_t=[]

A2B2f_t=[]

A2m_t=[]

B2m_t=[]

A2B2m_t=[]

total=0 # keeps count of the total number of cases explored for each k1-k2

for h in H: # iterating over values of dominance coefficient

for sf in Sf: # iterating over values of female selection coefficient

SM=np.arange(max(0,min(1,2*sf*h/(1+sf*h))),max(0,min(1,2*sf*(1-h)/(1-sf*h)))+0.00001,0.1) # determining range of male selection coefficients based on Connallon and Clark (2010)

Sm=[round(x,1) for x in SM]

for sm in Sm: # iterating over values of male selection coefficient

#fitness scheme

f11= 1

f13= 1-k1*sf

f31= 1-sf

f33= 1-(1-k2)*sf

f12= 1-k1*h*sf

f32= 1 - (1+h - h*(1-k2)-k2)*sf

f21= 1- h*sf

f23= 1 - abs((1-k1-k2)*h*sf)

f22r= 1- abs(h*sf*(1-k1))

f22c= 1-(1-k2)*sf*h

m1=1-sm

m2=1-sm

m3=1

m4=1

\# setting initial frequencies of alleles in different sexes

A1f=max(0.01,min(0.99,(2*sf*(1-h)-sm)/(2*sf*(1-2*h))))

A2f=1- A1f

A2m=max(0.01,min(0.99,(sm-2*sf*h)/(2*sf*(1-2*h))))

A1m=1-A2m

B1=0.95

B2=0.05

\# calculating initial frequencies for each haplotype

x1=A1f*B1

x2=A1f*B2

x3=A2f*B1

x4=A2f*B2

y1=A1m*B1

y2=A1m*B2

y3=A2m*B1

y4=A2m*B2

possible_D=[0.25,1-x1,x1-0,1-x2,x2-0,1-x3,x3-0,1-x4,x4-0,1-y1,y1-0,1-y2,y2-0,1-y3,y3-0,1-y4,y4-0] # determining the maximum possible value of linkage disequilibrium (D) based on initial frequencies

Drange=np.arange(-min(possible_D),min(possible_D),0.01) # setting the range of values for D

for D in Drange: # iterating over values of linkage disequilibrium

for rf in Rf: # iterating over values of female recombination rates

\# generating the time series

\# calculating initial frequencies of each haplotype

x1=A1f*B1

x2=A1f*B2

x3=A2f*B1

x4=A2f*B2

y1=A1m*B1

y2=A1m*B2

y3=A2m*B1

y4=A2m*B2

x1+=D

x2+=-D

x3+=-D

x4+=D

y1+=D

y2+=-D

y3+=-D

y4+=D

\# lists to store the time series

X1=[x1]

X2=[x2]

X3=[x3]

X4=[x4]

Y1=[y1]

Y2=[y2]

Y3=[y3]

Y4=[y4]

\# keeping track of whether alleles or haplotypes are fixed

A2B2_fixed=False

A2fB=False

B2fB=False

A2B2fB=False

A2mB=False

B2mB=False

A2B2mB=False

total+=1 # incrementing the total count of cases covered by 1

for t in range(T):

\# calculating haplotype frequencies in (t+1)^th generation

wm=x1*m1+x2*m2+x3*m3+x4*m4

wf=x1*y1*f11 + (x1*y2+x2*y1)*f12 + x2*y2*f13 + (x1*y3+x3*y1)*f21 + (x2*y3+x3*y2)*f22r + (x1*y4+x4*y1)*f22c + (x2*y4+x4*y2)*f23 + x3*y3*f31 + (x3*y4+x4*y3)*f32 + x4*y4*f33

Y1+=[x1*m1/wm]

Y2+=[x2*m2/wm]

Y3+=[x3*m3/wm]

Y4+=[x4*m4/wm]

X1+=[(2*x1*y1*f11 + (x1*y2+x2*y1)*f12 + (x1*y3+x3*y1)*f21 + (1-rf)*f22c*(x1*y4+x4*y1) + rf*f22r*(x2*y3+x3*y2))/(2*wf)]

X2+=[(2*x2*y2*f13 + (x1*y2+x2*y1)*f12 + (x2*y4+x4*y2)*f23 + (1-rf)*f22r*(x3*y2+x2*y3) + rf*f22c*(x4*y1+x1*y4))/(2*wf)]

X3+=[(2*x3*y3*f31 + (x1*y3+x3*y1)*f21 + (x3*y4+x4*y3)*f32 + (1-rf)*f22r*(x2*y3+x3*y2) + rf*f22c*(x4*y1+x1*y4))/(2*wf)]

X4+=[(2*x4*y4*f33 + (x2*y4+x4*y2)*f23 + (x4*y3+x3*y4)*f32 + (1-rf)*f22c*(x4*y1+x1*y4) + rf*f22r*(x2*y3+x3*y2))/(2*wf)]

x1=X1[-1]

x2=X2[-1]

x3=X3[-1]

x4=X4[-1]

y1=Y1[-1]

y2=Y2[-1]

y3=Y3[-1]

y4=Y4[-1]

\# safety checks for negative frequencies

if x1<0 or x2<0 or x3<0 or x4<0:

print('WRONG X',x1,x2,x3,x4)

print('k1,k2,h,sm,sf,wf, D, rf, minD-',k1,k2,h,sm,sf,wf,D,rf,min(possible_D))

break

if y1<0 or y2<0 or y3<0 or y4<0:

print('WRONG Y',y1,y2,y3,y4)

print('k1,k2,h,sm,sf,wm, D, rf, minD-',k1,k2,h,sm,sf,wm,D,rf,min(possible_D))

break

\# checking for fixation of different alleles or haplotypes

if round((x3+x4),3)==1 and A2fB==False:

A2f_fix+=1

A2f_t+=[t]

A2fB=True

if round((x2+x4),3)==1 and B2fB==False:

B2f_fix+=1

B2f_t+=[t]

B2fB=True

if round(x4,3)==1 and A2B2fB==False:

A2B2f_fix+=1

A2B2f_t+=[t]

A2B2fB=True

if round((y3+y4),3)==1 and A2mB==False:

A2m_fix+=1

A2m_t+=[t]

A2mB=True

if round((y2+y4),3)==1 and B2mB==False:

B2m_fix+=1

B2m_t+=[t]

B2mB=True

if round(y4,3)==1 and A2B2mB==False:

A2B2m_fix+=1

A2B2m_t+=[t]

A2B2mB=True

\# check for all haplotype frequencies adding upto 1

if round(x1+x2+x3+x4,3) not in checklist:

print(f'x not summing upto 1-{x1+x2+x3+x4}',x1,x2,x3,x4,t)

break

if round(y1+y2+y3+y4,3) not in checklist:

print(f'y not summing upto 1-{y1+y2+y3+y4}',y1,y2,y3,y4,t)

break

data=[round(A2f_fix/total,3),round(B2f_fix/total,3),round(A2B2f_fix/total,3),round(A2m_fix/total,3),round(B2m_fix/total,3),round(A2B2m_fix/total,3)] # rounding off values of fraction of conflict resolution to 3 decimal places

Worksheet=op.load_workbook('ParameterSpace fixationX.xlsx') # opening excel sheet to store the data collected

\# initializing sheets to store data for individual alleles/haplotypes

A2F=Worksheet['A2 female']

B2F=Worksheet['B2 female']

A2B2F=Worksheet['A2B2 female']

A2M=Worksheet['A2 male']

B2M=Worksheet['B2 male']

A2B2M=Worksheet['A2B2 male']

sheets=[A2F,B2F,A2B2F,A2M,B2M,A2B2M]

\# setting initial row and column values

ROW=0

COLUMN=0

#print('k1,k2',k1,k2)

for i in range(2,13,1): # iterating over rows in the sheet

if float(A2F.cell(row=i,column=1).value)==k1: # finding the row corresponding to k1

for j in range(2,13,1): # iterating over columns in the sheet

if float(A2F.cell(row=1,column=j).value)==k2: # finding the column corresponding to k2

ROW=i

COLUMN=j

#print(ROW,COLUMN)

for i in range(6):

sheets[i].cell(ROW,COLUMN).value=data[i] # adding data to the correct cell in the appropriate sheet

Worksheet.save('ParameterSpace fixationX.xlsx') # saving file

\# setting variables to None for reuse

Worksheet=None

A2F=None

B2F=None

A2B2F=None

A2M=None

B2M=None

A2B2M=None

listS=[A2f_t,B2f_t,A2B2f_t,A2m_t,B2m_t,A2B2m_t] # list of lists of time taken to fix the allele/haplotype

for x in listS:

if x==[]: # setting value to T+25 if no fixation was achieved

x+=[T+25]

data=[round(mean(x),3) for x in listS] # calculating mean time taken to fix the allele/haplotype

Worksheet=op.load_workbook('ParameterSpace timeX.xlsx') # loading time Workbook

\# initializing sheets

A2F=Worksheet['A2 female']

B2F=Worksheet['B2 female']

A2B2F=Worksheet['A2B2 female']

A2M=Worksheet['A2 male']

B2M=Worksheet['B2 male']

A2B2M=Worksheet['A2B2 male']

sheets=[A2F,B2F,A2B2F,A2M,B2M,A2B2M]

#print('k1,k2',k1,k2)

for i in range(2,13,1): # finding correct row and column to enter the data into

if float(A2F.cell(row=i,column=1).value)==k1:

for j in range(2,13,1):

if float(A2F.cell(row=1,column=j).value)==k2:

ROW=i

COLUMN=j

#print(ROW,COLUMN)

for i in range(6):

sheets[i].cell(ROW,COLUMN).value=data[i] # entering data

print (data[i])

Worksheet.save('ParameterSpace timeX.xlsx') # saving Workbook

\# setting variables to None for reuse

Worksheet=None

A2F=None

B2F=None

A2B2F=None

A2M=None

B2M=None

A2B2M=None

**Code 3 ‘comparison.py’: This code was used to visualize the data generated from Code 1 and Code 2, and construct Figures 1 and 2.**

#importing libraries

import openpyxl as op

import seaborn as sns

import matplotlib.pyplot as plt

import numpy as np

#loading worksheets with the data

Afile=op.load_workbook('ParameterSpace fixationA.xlsx')

Atile=op.load_workbook('ParameterSpace timeA.xlsx')

Xfile=op.load_workbook('ParameterSpace fixationX.xlsx')

Xtile=op.load_workbook('ParameterSpace timeX.xlsx')

files=[Afile,Atile,Xfile,Xtile]

alldata=[]#will be in sequence of Afreq,Atime, Xfreq, Xtile

for i in range(len(files)):

Worksheet=files[i]

#creating objects for individual sheets within the file

A2F=Worksheet['A2 female']

B2F=Worksheet['B2 female']

A2B2F=Worksheet['A2B2 female']

A2M=Worksheet['A2 male']

B2M=Worksheet['B2 male']

A2B2M=Worksheet['A2B2 male']

sheets=[[A2F,A2M],[B2F,B2M],[A2B2F,A2B2M]] #clubbing the sheets

data=[[] for sheet in sheets]

for j,sheet in enumerate(sheets): # iterating over sheets

for r in range(2,sheet[0].max_row+1): # iterating over rows

row=[]

for c in range(2,sheet[0].max_column+1): # iterating over columns & adding values to a list

if i in [1,3]: # setting avg fixation time to T+25 for the (k1,k2) where no fixation occurs

val=(sheet[0].cell(r,c).value + sheet[1].cell(r,c).value)/2 # taking average of the parameter value over males and females

if val==0:

val=3025

row+=[val]

else:

row+=[(sheet[0].cell(r,c).value + sheet[1].cell(r,c).value)/2] # taking average of the parameter value over males and females

data[j]+=[row]

alldata+=[data] # adding list to main matrix

\# resetting sheet objects

A2F=None

B2F=None

A2B2F=None

A2M=None

B2M=None

A2B2M=None

#plotting the data extracted from the files

figfreq,axfreq=plt.subplots(3,2,sharex=True,sharey=True,figsize=(17,20),gridspec_kw={'hspace':0.25,'wspace':0.125}) #setting figure parameters

freqdata= [alldata[i] for i in [0,2]] # making matrix of fraction of fixation data

cbar_ax = figfreq.add_axes([.91, .3, .03, .4]) # setting the colorbar axis

for j,data in enumerate(freqdata): # iterating over autosomal and x fixation data

for i,map in enumerate(data): # iterating over A2, B2, and A2B2

\# setting row and column values

aOrx=0 # Autosomal or X-linked

if j>0:aOrx=1

row=i

column=aOrx

#plotting the heatmap within subplots

g=sns.heatmap(map,vmin=0,vmax=1,yticklabels=[0,0.1,0.2,0.3,0.4,0.5,0.6,0.7,0.8,0.9,1],xticklabels=[0,0.1,0.2,0.3,0.4,0.5,0.6,0.7,0.8,0.9,1],cmap='cividis',cbar=True,cbar_ax=cbar_ax, cbar_kws={'shrink': 1.8,"use_gridspec":False},ax=axfreq[row,column])

#formatting the heatmap

axfreq[row,column].set_yticklabels([0,0.1,0.2,0.3,0.4,0.5,0.6,0.7,0.8,0.9,1],fontsize=15)

axfreq[row,column].set_xticklabels([0,0.1,0.2,0.3,0.4,0.5,0.6,0.7,0.8,0.9,1],fontsize=15)

\# setting subplot titles and labelling rows (on the left) and columns (bottom)

if row==0:

Title=''

if aOrx==0:Title+='Autosomal Loci'

else:Title+='X-linked Loci'

axfreq[row,column].set_title(Title,fontsize=30)

if column==0:

ylabel=''

if row==0:ylabel='A2'

if row==1:ylabel='B2'

if row==2:ylabel='A2B2'

axfreq[row,column].set_ylabel(ylabel+'\n k1',fontsize=25)

if row==2:

axfreq[row,column].set_xlabel('k2',fontsize=25)

\# customizing colorbar

cbar_ax.set_yticklabels([round(x,2) for x in np.arange(0,1.1,0.2)],fontsize=20)

cbar_ax.set_ylabel('Fraction of Parameter Space where A2B2 gets fixed',fontsize=25)

figfreq.savefig('X-A comparing ratio of ParameterSpace.tiff',dpi=300) # save figure

figtime,axtime=plt.subplots(3,2,sharex=True,sharey=True,figsize=(17,20),gridspec_kw={'hspace':0.25,'wspace':0.125}) # setting figure parameters

timedata= [alldata[i] for i in [1,3]] # constructing matrix of the average time of fixation

cbar_ax=None # colorbar axis reset

cbar_ax = figtime.add_axes([.91, .3, .03, .4]) # setting new colorbar axis

for j,data in enumerate(timedata): # iterating over autosomes and x data

for i,map in enumerate(data): # iterating over A2, B2, and A2B2

\# setting row and column

aOrx=0 # Autosomal or X-linked

if j>0:aOrx=1

row=i

column=aOrx

#plotting the heatmap

g=sns.heatmap(map,vmin=0,vmax=3025,yticklabels=[0,0.1,0.2,0.3,0.4,0.5,0.6,0.7,0.8,0.9,1],xticklabels=[0,0.1,0.2,0.3,0.4,0.5,0.6,0.7,0.8,0.9,1],cmap='cividis',cbar_ax=cbar_ax,cbar=True,cbar_kws={'shrink': 1.8,"use_gridspec":False},ax=axtime[row,column])

\# formatting the heatmap

axtime[row,column].set_yticklabels([0,0.1,0.2,0.3,0.4,0.5,0.6,0.7,0.8,0.9,1],fontsize=15)

axtime[row,column].set_xticklabels([0,0.1,0.2,0.3,0.4,0.5,0.6,0.7,0.8,0.9,1],fontsize=15)

\# setting subplot titles and labelling rows (on the left) and columns (bottom)

if row==0:

Title=''

if aOrx==0:Title+='Autosomal Loci'

else:Title+='X-linked Loci'

axtime[row,column].set_title(Title,fontsize=30)

if column==0:

ylabel=''

if row==0:ylabel='A2'

if row==1:ylabel='B2'

if row==2:ylabel='A2B2'

axtime[row,column].set_ylabel(ylabel+'\n k1',fontsize=25)

if row==2:

axtime[row,column].set_xlabel('k2',fontsize=25)

\# formatting the colorbar

cbar_ax.set_yticklabels(range(0,3026,500),fontsize=15)

cbar_ax.set_ylabel('Average Time Taken to achieve Fixation',fontsize=25)

figtime.savefig('X-A comparing time.tiff',dpi=300) #saving the figure

**Code 4 ‘Autosome_hsmsf.py’: This code was used to know how dominance coefficient (h) and selection coefficients (sm and sf) affected the system with different kinds of modifiers, when the loci were present on autosomes. This generated the figures present in video 1 ‘A-h-Frequency.mp4’ and in video 3 ‘A-h-Time.mp4’.**

\# importing libraries

import numpy as np

import matplotlib.pyplot as plt

import seaborn as sns

from random import uniform

h=0.1 # set value of dominance coefficient

SF=np.arange(0.2,1.01,0.2)

Sf=[0.001]+[round(x,2) for x in SF] # range of selection coefficient for females

Checklist=np.arange(1-0.020,1+0.020,0.001)

checklist=[round(i,3) for i in Checklist] # a checklist to cross check whether the frequencies are summing upto approximately 1 at every time step

T=3000 # number of generations

rf=0.2 # female recombination rate

rm =0.2 # male recombination rate

Krange=np.arange(0,1.01,0.2)

krange=[round(k,2) for k in Krange] # range of values for k1 and k2

timedata=[[[[T+25 for sm in Sf] for sf in Sf] for k2 in krange] for k1 in krange] # array storing fixation time data

freqdata=[[[[T+25 for sm in Sf] for sf in Sf] for k2 in krange] for k1 in krange] # array storing frequency of A2B2 at 3000th generation

for sfind,sf in enumerate(Sf): # iterating over values of sf

for smind,sm in enumerate(Sf): # iterating over values of sm

\# setting initial frequencies of various alleles

A2f=max(0.01,min(0.99,(sm*(1-h)-sf*h)/((sm+sf)*(1-2*h))))

A1f=1-A2f

A2m=max(0.01,min(0.99,(sm*(1-h)-sf*h)/((sm+sf)*(1-2*h))))

A1m=1-A2m

B1=0.95

B2=0.05

D=0 # setting value of linkage disequilibrium to 0

for k1ind,k1 in enumerate(krange): # iterating over values of k1

for k2ind,k2 in enumerate(krange): # iterating over values of k2

\# calculating initial frequencies of every haplotype

x1=A1f*B1+D

x2=A1f*B2-D

x3=A2f*B1-D

x4=A2f*B2+D

y1=A1m*B1+D

y2=A1m*B2-D

y3=A2m*B1-D

y4=A2m*B2+D

#Storing frequencies

X1=[x1]

X2=[x2]

X3=[x3]

X4=[x4]

Y1=[y1]

Y2=[y2]

Y3=[y3]

Y4=[y4]

#fitness scheme

f11= 1

f13= 1-k1*sf

f31= 1-sf

f33= 1-(1-k2)*sf

f12= 1-k1*h*sf

f32= 1 - (1+h - h*(1-k2)-k2)*sf

f21= 1- h*sf

f23= 1 - abs((1-k1-k2)*h*sf)

f22r= 1- abs(h*sf*(1-k1))

f22c= 1-(1-k2)*sf*h

m11= 1-sm

m21= 1 - h*sm

m33= 1

m12=1 - sm

m13=1 - sm

m22r=1 - h*sm

m22c=1 - h*sm

m23=1 - h*sm

m31=1

m32=1

A2B2_fixed=False # a variable to keep check of whether the haplotype A2B2 is fixed

\# generating time series

for t in range(T):

\# computing haplotype frequencies at (t+1)th generation

wm = x1*y1*m11 + (x1*y2+x2*y1)*m12 + x2*y2*m13 + (x1*y3+x3*y1)*m21 + (x2*y3+x3*y2)*m22r + (x1*y4+x4*y1)*m22c + (x2*y4+x4*y2)*m23 + x3*y3*m31 + (x3*y4+x4*y3)*m32 + x4*y4*m33

wf=x1*y1*f11 + (x1*y2+x2*y1)*f12 + x2*y2*f13 + (x1*y3+x3*y1)*f21 + (x2*y3+x3*y2)*f22r + (x1*y4+x4*y1)*f22c + (x2*y4+x4*y2)*f23 + x3*y3*f31 + (x3*y4+x4*y3)*f32 + x4*y4*f33

Y1+=[(2*x1*y1*m11 + (x1*y2+x2*y1)*m12 + (x1*y3+x3*y1)*m21 + (1-rm)*m22c*(x1*y4+x4*y1) + rm*m22r*(x2*y3+x3*y2))/(2*wm)]

Y2+=[(2*x2*y2*m13 + (x1*y2+x2*y1)*m12 + (x2*y4+x4*y2)*m23 + (1-rm)*m22r*(x3*y2+x2*y3) + rm*m22c*(x4*y1+x1*y4))/(2*wm)]

Y3+=[(2*x3*y3*m31 + (x1*y3+x3*y1)*m21 + (x3*y4+x4*y3)*m32 + (1-rm)*m22r*(x2*y3+x3*y2) + rm*m22c*(x4*y1+x1*y4))/(2*wm)]

Y4+=[(2*x4*y4*m33 + (x2*y4+x4*y2)*m23 + (x4*y3+x3*y4)*m32 + (1-rm)*m22c*(x4*y1+x1*y4) + rm*m22r*(x2*y3+x3*y2))/(2*wm)]

X1+=[(2*x1*y1*f11 + (x1*y2+x2*y1)*f12 + (x1*y3+x3*y1)*f21 + (1-rf)*f22c*(x1*y4+x4*y1) + rf*f22r*(x2*y3+x3*y2))/(2*wf)]

X2+=[(2*x2*y2*f13 + (x1*y2+x2*y1)*f12 + (x2*y4+x4*y2)*f23 + (1-rf)*f22r*(x3*y2+x2*y3) + rf*f22c*(x4*y1+x1*y4))/(2*wf)]

X3+=[(2*x3*y3*f31 + (x1*y3+x3*y1)*f21 + (x3*y4+x4*y3)*f32 + (1-rf)*f22r*(x2*y3+x3*y2) + rf*f22c*(x4*y1+x1*y4))/(2*wf)]

X4+=[(2*x4*y4*f33 + (x2*y4+x4*y2)*f23 + (x4*y3+x3*y4)*f32 + (1-rf)*f22c*(x4*y1+x1*y4) + rf*f22r*(x2*y3+x3*y2))/(2*wf)]

x1=X1[-1]

x2=X2[-1]

x3=X3[-1]

x4=X4[-1]

y1=Y1[-1]

y2=Y2[-1]

y3=Y3[-1]

y4=Y4[-1]

#checking for haplotype frequencies summing upto 1

if round(x1+x2+x3+x4,3) not in checklist:

print(f'x not summing upto 1-{x1+x2+x3+x4}',x1,x2,x3,x4,t)

break

if round(y1+y2+y3+y4,3) not in checklist:

print(f'y not summing upto 1-{y1+y2+y3+y4}',y1,y2,y3,y4,t)

break

\# checking for fixation of A2B2

if round((x4+y4)/2,3)==1 and A2B2_fixed==False:

timedata[k1ind][k2ind][sfind][smind]=t # adding the time of fixation to array

A2B2_fixed=True

\# adding the frequency of 3000th generation to array

if t==T-1:

freqdata[k1ind][k2ind][sfind][smind]=round((x4+y4)/2,3)

figfreq,axfreq=plt.subplots(len(krange),len(krange),sharey=True,sharex=True,figsize=(40,40)) #setting figure parameters

cbar_ax = figfreq.add_axes([.91, .3, .03, .4]) # setting the colorbar axis

for i,k1 in enumerate(freqdata): # iterating over frequency data

for j,map in enumerate(k1): # iterating over frequency data

g=sns.heatmap(map,vmin=0,vmax=1,cmap='YlGn',cbar=True,cbar_ax=cbar_ax, cbar_kws={'shrink': 1.8,"use_gridspec":False},ax=axfreq[i,j]) # plotting the heatmap

\# formatting the heatmap

axfreq[i,j].set_yticklabels(Sf,fontsize=25)

axfreq[i,j].set_xticklabels(Sf,fontsize=25)

\# setting subplot titles and labelling rows (on the left) and columns (bottom)

if i==len(timedata)-1:

axfreq[i,j].set_xlabel(f'sm \n k2 = {krange[j]}', fontsize=35)

if j==0:

axfreq[i,j].set_ylabel(f'k1 = {krange[i]} \n sf', fontsize=35)

axfreq[0,0].set_title(f'h = {h}',fontsize=50)

\# customizing colorbar

cbar_ax.set_yticklabels([round(x,2) for x in np.arange(0,1.1,0.2)],fontsize=30)

cbar_ax.set_ylabel(f'Frequency of A2B2 at generation {T}',fontsize=35)

figfreq.savefig(f'Freq at 3000 h={h}.tiff',dpi = 300) # save figure

cbar_ax=None # colorbar axis reset

figtime,axtime=plt.subplots(len(krange),len(krange),sharey=True,sharex=True,figsize=(40,40),gridspec_kw={'hspace':0.25,'wspace':0.125}) # setting figure parameters)

cbar_ax = figtime.add_axes([.91, .3, .03, .4]) # setting new colorbar axis

for i,k1 in enumerate(timedata): # iterating over time data

for j,map in enumerate(k1): # iterating over time data

g=sns.heatmap(map,vmin=0,vmax=T+25,yticklabels=Sf,xticklabels=Sf,cmap='GnBu',cbar=True,cbar_ax=cbar_ax, cbar_kws={'shrink': 1.8,"use_gridspec":False},ax=axtime[i,j]) # plotting the heatmap

\# formatting the heatmap

axtime[i,j].set_yticklabels(Sf,fontsize=25)

axtime[i,j].set_xticklabels(Sf,fontsize=25)

\# setting subplot titles and labelling rows (on the left) and columns (bottom)

if i==len(timedata)-1:

axtime[i,j].set_xlabel(f'sm \n k2 = {krange[j]}',fontsize=35)

if j==0:

axtime[i,j].set_ylabel(f'k1 = {krange[i]} \n sf', fontsize=35)

axtime[0,0].set_title(f'h = {h}',fontsize=50)

\# formatting the colorbar

cbar_ax.set_yticklabels(range(0,3026,500),fontsize=30)

cbar_ax.set_ylabel('Time Taken to Fix A2B2',fontsize=35)

figtime.savefig(f'Time h={h}.tiff', dpi=300)

**Code 5 ‘X_hsmsf.py’: This code was used to know how dominance coefficient (h) and selection coefficients (sm and sf) affected the system with different kinds of modifiers, when the loci were present on X chromosomes. This generated the figures present in video 2 ‘X-h-Frequency.mp4’ and in video 4 ‘X-h-Time.mp4’.**

\# importing libraries

import numpy as np

import matplotlib.pyplot as plt

import seaborn as sns

from random import uniform

h=0.1 # setting value of dominance coefficient for current run

SF=np.arange(0.2,1.01,0.2)

Sf=[0.001]+[round(x,2) for x in SF] # range of selection coefficient for females

Checklist=np.arange(1-0.020,1+0.020,0.001)

checklist=[round(i,3) for i in Checklist] # a checklist to cross check whether the frequencies are summing upto approximately 1 at every time step

T=3000 # number of generations

rf=0.2 # female recombination rate

Krange=np.arange(0,1.01,0.2)

krange=[round(k,2) for k in Krange] # range of values for k1 and k2

timedata=[[[[T+25 for sm in Sf] for sf in Sf] for k2 in krange] for k1 in krange] # array storing fixation time data

freqdata=[[[[T+25 for sm in Sf] for sf in Sf] for k2 in krange] for k1 in krange] # array storing frequency of A2B2 at 3000th generation

for sfind,sf in enumerate(Sf): # iterating over values of selection coefficients in females and males

for smind,sm in enumerate(Sf):

\# setting initial frequencies of various alleles

A1f=max(0.01,min(0.99,(2*sf*(1-h)-sm)/(2*sf*(1-2*h))))

A2f=1- A1f

A2m=max(0.01,min(0.99,(sm-2*sf*h)/(2*sf*(1-2*h))))

A1m=1-A2m

B1=0.95

B2=0.05

for i,k1 in enumerate(krange): # iterating over values of k1 and k2

for j,k2 in enumerate(krange):

\# setting initial frequencies for different haplotypes assuming initial linkage disequilibrium to be 0

x1=A1f*B1

x2=A1f*B2

x3=A2f*B1

x4=A2f*B2

y1=A1m*B1

y2=A1m*B2

y3=A2m*B1

y4=A2m*B2

#Storing frequencies

X1=[x1]

X2=[x2]

X3=[x3]

X4=[x4]

Y1=[y1]

Y2=[y2]

Y3=[y3]

Y4=[y4]

#fitness scheme

f11= 1

f13= 1-k1*sf

f31= 1-sf

f33= 1-(1-k2)*sf

f12= 1-k1*h*sf

f32= 1 - (1+h - h*(1-k2)-k2)*sf

f21= 1- h*sf

f23= 1 - abs((1-k1-k2)*h*sf)

f22r= 1- abs(h*sf*(1-k1))

f22c= 1-(1-k2)*sf*h

m1=1-sm

m2=1-sm

m3=1

m4=1

#time series

A2B2_fixed=False # a variable to keep check of whether the haplotype A2B2 is fixed

for t in range(T):

\# computing haplotype frequencies at (t+1)th generation

wm=x1*m1+x2*m2+x3*m3+x4*m4

wf=x1*y1*f11 + (x1*y2+x2*y1)*f12 + x2*y2*f13 + (x1*y3+x3*y1)*f21 + (x2*y3+x3*y2)*f22r + (x1*y4+x4*y1)*f22c + (x2*y4+x4*y2)*f23 + x3*y3*f31 + (x3*y4+x4*y3)*f32 + x4*y4*f33

Y1+=[x1*m1/wm]

Y2+=[x2*m2/wm]

Y3+=[x3*m3/wm]

Y4+=[x4*m4/wm]

X1+=[(2*x1*y1*f11 + (x1*y2+x2*y1)*f12 + (x1*y3+x3*y1)*f21 + (1-rf)*f22c*(x1*y4+x4*y1) + rf*f22r*(x2*y3+x3*y2))/(2*wf)]

X2+=[(2*x2*y2*f13 + (x1*y2+x2*y1)*f12 + (x2*y4+x4*y2)*f23 + (1-rf)*f22r*(x3*y2+x2*y3) + rf*f22c*(x4*y1+x1*y4))/(2*wf)]

X3+=[(2*x3*y3*f31 + (x1*y3+x3*y1)*f21 + (x3*y4+x4*y3)*f32 + (1-rf)*f22r*(x2*y3+x3*y2) + rf*f22c*(x4*y1+x1*y4))/(2*wf)]

X4+=[(2*x4*y4*f33 + (x2*y4+x4*y2)*f23 + (x4*y3+x3*y4)*f32 + (1-rf)*f22c*(x4*y1+x1*y4) + rf*f22r*(x2*y3+x3*y2))/(2*wf)]

x1=X1[-1]

x2=X2[-1]

x3=X3[-1]

x4=X4[-1]

y1=Y1[-1]

y2=Y2[-1]

y3=Y3[-1]

y4=Y4[-1]

\# checking for fixation of A2B2

if round((x4+y4)/2,3)==1 and A2B2_fixed==False:

timedata[i][j][sfind][smind]=t # adding the time of fixation to array

A2B2_fixed=True

\# adding the frequency of 3000th generation to array

if t==T-1:

freqdata[i][j][sfind][smind]=round((x4+y4)/2,3)

#checking for haplotype frequencies summing upto 1

if round(x1+x2+x3+x4,3) not in checklist:

print(f'x not summing upto 1-{x1+x2+x3+x4}',x1,x2,x3,x4,t)

break

if round(y1+y2+y3+y4,3) not in checklist:

print(f'y not summing upto 1-{y1+y2+y3+y4}',y1,y2,y3,y4,t)

break

figfreq,axfreq=plt.subplots(len(krange),len(krange),sharey=True,sharex=True,figsize=(40,40)) #setting figure parameters

cbar_ax = figfreq.add_axes([.91, .3, .03, .4]) # setting the colorbar axis

for i,k1 in enumerate(freqdata): # iterating over frequency data

for j,map in enumerate(k1): # iterating over frequency data

g=sns.heatmap(map,vmin=0,vmax=1,cmap='YlGn',cbar=True,cbar_ax=cbar_ax, cbar_kws={'shrink': 1.8,"use_gridspec":False},ax=axfreq[i,j]) # plotting the heatmap

\# formatting the heatmap

axfreq[i,j].set_yticklabels(Sf,fontsize=25)

axfreq[i,j].set_xticklabels(Sf,fontsize=25)

\# setting subplot titles and labelling rows (on the left) and columns (bottom)

if i==len(timedata)-1:

axfreq[i,j].set_xlabel(f'sm \n k2 = {krange[j]}', fontsize=35)

if j==0:

axfreq[i,j].set_ylabel(f'k1 = {krange[i]} \n sf', fontsize=35)

axfreq[0,0].set_title(f'h = {h}',fontsize=50)

\# customizing colorbar

cbar_ax.set_yticklabels([round(x,2) for x in np.arange(0,1.1,0.2)],fontsize=30)

cbar_ax.set_ylabel(f'Frequency of A2B2 at generation {T}',fontsize=35)

figfreq.savefig(f'X linked Freq at 3000 h={h}.tiff',dpi = 300) # save figure

cbar_ax=None # colorbar axis reset

figtime,axtime=plt.subplots(len(krange),len(krange),sharey=True,sharex=True,figsize=(40,40),gridspec_kw={'hspace':0.25,'wspace':0.125}) # setting figure parameters)

cbar_ax = figtime.add_axes([.91, .3, .03, .4]) # setting new colorbar axis

for i,k1 in enumerate(timedata): # iterating over time data

for j,map in enumerate(k1): # iterating over time data

g=sns.heatmap(map,vmin=0,vmax=T+25,yticklabels=Sf,xticklabels=Sf,cmap='GnBu',cbar=True,cbar_ax=cbar_ax, cbar_kws={'shrink': 1.8,"use_gridspec":False},ax=axtime[i,j]) # plotting the heatmap

\# formatting the heatmap

axtime[i,j].set_yticklabels(Sf,fontsize=25)

axtime[i,j].set_xticklabels(Sf,fontsize=25)

\# setting subplot titles and labelling rows (on the left) and columns (bottom)

if i==len(timedata)-1:

axtime[i,j].set_xlabel(f'sm \n k2 = {krange[j]}',fontsize=35)

if j==0:

axtime[i,j].set_ylabel(f'k1 = {krange[i]} \n sf', fontsize=35)

axtime[0,0].set_title(f'h = {h}',fontsize=50)

\# formatting the colorbar

cbar_ax.set_yticklabels(range(0,3026,500),fontsize=30)

cbar_ax.set_ylabel('Time Taken to Fix A2B2',fontsize=35)

figtime.savefig(f'X linked Time h={h}.tiff', dpi=300)

**Code 5 ‘Autosome_Drmrf.py’: This code was used to know how initial linkage disequilibrium (D) and recombination rates (rm and rf) affected the system with different kinds of modifiers, when the loci were present on autosomes. This generated the figures present in video 5 ‘A-D-Frequency.mp4’ and in video 6 ‘A-D-Time.mp4’.**

\# importing libraries

import numpy as np

import matplotlib.pyplot as plt

import seaborn as sns

from random import uniform

RF=np.arange(0,0.6,0.1)

Rf=[round(x,2) for x in RF] # range of recombination rates

h=0.2 # setting value of dominance coefficient

sf=0.2 # setting value of female selection coefficient

sm=0.2 # setting value of male selection coefficient

Checklist=np.arange(1-0.020,1+0.020,0.001)

checklist=[round(i,3) for i in Checklist] # a checklist to cross check whether the frequencies are summing upto approximately 1 at every time step

T=3000 # number of generations

Krange=np.arange(0,1.01,0.2)

krange=[round(k,2) for k in Krange] # range of values for k1 and k2

\# determining initial frequencies of various alleles

A2f=max(0.01,min(0.99,(sm*(1-h)-sf*h)/((sm+sf)*(1-2*h))))

A1f=1-A2f

A2m=max(0.01,min(0.99,(sm*(1-h)-sf*h)/((sm+sf)*(1-2*h))))

A1m=1-A2m

B1=0.95

B2=0.05

\# determining initial frequencies of various haplotypes

x1=A1f*B1

x2=A1f*B2

x3=A2f*B1

x4=A2f*B2

y1=A1m*B1

y2=A1m*B2

y3=A2m*B1

y4=A2m*B2

\# determining possible values of D

possible_D=[1-x1,x1-0,1-x2,x2-0,1-x3,x3-0,1-x4,x4-0,1-y1,y1-0,1-y2,y2-0,1-y3,y3-0,1-y4,y4-0]

DRange=np.arange(-min(possible_D),min(possible_D),0.01)

Drange=[round(x,2) for x in DRange]

print(Drange)

D=Drange[4] # setting value of linkage disequilibrium for the current run

timedata=[[[[T+25 for rm in Rf] for rf in Rf] for k2 in krange] for k1 in krange] # array storing fixation time data

freqdata=[[[[T+25 for rm in Rf] for rf in Rf] for k2 in krange] for k1 in krange] # array storing frequency of A2B2 at 3000th generation

for rmind,rm in enumerate(Rf): # iterating over values of rm and rf

for rfind,rf in enumerate(Rf):

for k1ind,k1 in enumerate(krange): # iterating over values of k1 and k2

for k2ind,k2 in enumerate(krange):

#fitness scheme

f11= 1

f13= 1-k1*sf

f31= 1-sf

f33= 1-(1-k2)*sf

f12= 1-k1*h*sf

f32= 1 - (1+h - h*(1-k2)-k2)*sf

f21= 1- h*sf

f23= 1 - abs((1-k1-k2)*h*sf)

f22r= 1- abs(h*sf*(1-k1))

f22c= 1-(1-k2)*sf*h

m11= 1-sm

m21= 1 - h*sm

m33= 1

m12=1 - sm

m13=1 - sm

m22r=1 - h*sm

m22c=1 - h*sm

m23=1 - h*sm

m31=1

m32=1

#time series

x1=A1f*B1

x2=A1f*B2

x3=A2f*B1

x4=A2f*B2

y1=A1m*B1

y2=A1m*B2

y3=A2m*B1

y4=A2m*B2

x1+=D

x2+=-D

x3+=-D

x4+=D

y1+=D

y2+=-D

y3+=-D

y4+=D

#Storing frequencies

X1=[x1]

X2=[x2]

X3=[x3]

X4=[x4]

Y1=[y1]

Y2=[y2]

Y3=[y3]

Y4=[y4]

A2B2_fixed=False # a variable to keep check of whether the haplotype A2B2 is fixed

\# generating time series

for t in range(T):

\# computing haplotype frequencies at (t+1)th generation

wm = x1*y1*m11 + (x1*y2+x2*y1)*m12 + x2*y2*m13 + (x1*y3+x3*y1)*m21 + (x2*y3+x3*y2)*m22r + (x1*y4+x4*y1)*m22c + (x2*y4+x4*y2)*m23 + x3*y3*m31 + (x3*y4+x4*y3)*m32 + x4*y4*m33

wf = x1*y1*f11 + (x1*y2+x2*y1)*f12 + x2*y2*f13 + (x1*y3+x3*y1)*f21 + (x2*y3+x3*y2)*f22r + (x1*y4+x4*y1)*f22c + (x2*y4+x4*y2)*f23 + x3*y3*f31 + (x3*y4+x4*y3)*f32 + x4*y4*f33

Y1+=[(2*x1*y1*m11 + (x1*y2+x2*y1)*m12 + (x1*y3+x3*y1)*m21 + (1-rm)*m22c*(x1*y4+x4*y1) + rm*m22r*(x2*y3+x3*y2))/(2*wm)]

Y2+=[(2*x2*y2*m13 + (x1*y2+x2*y1)*m12 + (x2*y4+x4*y2)*m23 + (1-rm)*m22r*(x3*y2+x2*y3) + rm*m22c*(x4*y1+x1*y4))/(2*wm)]

Y3+=[(2*x3*y3*m31 + (x1*y3+x3*y1)*m21 + (x3*y4+x4*y3)*m32 + (1-rm)*m22r*(x2*y3+x3*y2) + rm*m22c*(x4*y1+x1*y4))/(2*wm)]

Y4+=[(2*x4*y4*m33 + (x2*y4+x4*y2)*m23 + (x4*y3+x3*y4)*m32 + (1-rm)*m22c*(x4*y1+x1*y4) + rm*m22r*(x2*y3+x3*y2))/(2*wm)]

X1+=[(2*x1*y1*f11 + (x1*y2+x2*y1)*f12 + (x1*y3+x3*y1)*f21 + (1-rf)*f22c*(x1*y4+x4*y1) + rf*f22r*(x2*y3+x3*y2))/(2*wf)]

X2+=[(2*x2*y2*f13 + (x1*y2+x2*y1)*f12 + (x2*y4+x4*y2)*f23 + (1-rf)*f22r*(x3*y2+x2*y3) + rf*f22c*(x4*y1+x1*y4))/(2*wf)]

X3+=[(2*x3*y3*f31 + (x1*y3+x3*y1)*f21 + (x3*y4+x4*y3)*f32 + (1-rf)*f22r*(x2*y3+x3*y2) + rf*f22c*(x4*y1+x1*y4))/(2*wf)]

X4+=[(2*x4*y4*f33 + (x2*y4+x4*y2)*f23 + (x4*y3+x3*y4)*f32 + (1-rf)*f22c*(x4*y1+x1*y4) + rf*f22r*(x2*y3+x3*y2))/(2*wf)]

x1=X1[-1]

x2=X2[-1]

x3=X3[-1]

x4=X4[-1]

y1=Y1[-1]

y2=Y2[-1]

y3=Y3[-1]

y4=Y4[-1]

#checking for haplotype frequencies summing upto 1

if round(x1+x2+x3+x4,3) not in checklist:

print(f'x not summing upto 1-{x1+x2+x3+x4}',x1,x2,x3,x4,t)

break

if round(y1+y2+y3+y4,3) not in checklist:

print(f'y not summing upto 1-{y1+y2+y3+y4}',y1,y2,y3,y4,t)

break

\# checking whether A2B2 has fixed in the population

if round((x4+y4)/2,3)==1 and A2B2_fixed==False:

timedata[k1ind][k2ind][rfind][rmind]=t # adding the time of fixation to array

A2B2_fixed=True

\# adding the frequency of A2B2 at 3000th generation to array

if t==T-1:

freqdata[k1ind][k2ind][rfind][rmind]=round((x4+y4)/2,3)

figfreq,axfreq=plt.subplots(len(krange),len(krange),sharey=True,sharex=True,figsize=(40,40)) #setting figure parameters

cbar_ax = figfreq.add_axes([.91, .3, .03, .4]) # setting the colorbar axis

for i,k1 in enumerate(freqdata): # iterating over frequency data

for j,map in enumerate(k1): # iterating over frequency data

g=sns.heatmap(map,vmin=0,vmax=1,cmap='YlGn',cbar=True,cbar_ax=cbar_ax, cbar_kws={'shrink': 1.8,"use_gridspec":False},ax=axfreq[i,j]) # plotting the heatmap

\# formatting the heatmap

axfreq[i,j].set_yticklabels(Rf,fontsize=25)

axfreq[i,j].set_xticklabels(Rf,fontsize=25)

\# setting subplot titles and labelling rows (on the left) and columns (bottom)

if i==len(timedata)-1:

axfreq[i,j].set_xlabel(f'rm \n k2 = {krange[j]}', fontsize=35)

if j==0:

axfreq[i,j].set_ylabel(f'k1 = {krange[i]} \n rf', fontsize=35)

axfreq[0,0].set_title(f'D = {D}',fontsize=50)

\# customizing colorbar

cbar_ax.set_yticklabels([round(x,2) for x in np.arange(0,1.1,0.2)],fontsize=30)

cbar_ax.set_ylabel(f'Frequency of A2B2 at generation {T}',fontsize=35)

figfreq.savefig(f'Freq at 3000 D={D}.tiff',dpi = 300) # save figure

figtime,axtime=plt.subplots(len(krange),len(krange),sharey=True,sharex=True,figsize=(40,40),gridspec_kw={'hspace':0.25,'wspace':0.125}) # setting figure parameters)

cbar_ax = figtime.add_axes([.91, .3, .03, .4]) # setting new colorbar axis

for i,k1 in enumerate(timedata): # iterating over time data

for j,map in enumerate(k1): # iterating over time data

g=sns.heatmap(map,vmin=0,vmax=T+25,yticklabels=Rf,xticklabels=Rf,cmap='GnBu',cbar=True,cbar_ax=cbar_ax, cbar_kws={'shrink': 1.8,"use_gridspec":False},ax=axtime[i,j]) # plotting the heatmap

\# formatting the heatmap

axtime[i,j].set_yticklabels(Rf,fontsize=25)

axtime[i,j].set_xticklabels(Rf,fontsize=25)

\# setting subplot titles and labelling rows (on the left) and columns (bottom)

if i==len(timedata)-1:

axtime[i,j].set_xlabel(f'rm \n k2 = {krange[j]}',fontsize=35)

if j==0:

axtime[i,j].set_ylabel(f'k1 = {krange[i]} \n rf', fontsize=35)

axtime[0,0].set_title(f'D = {D}',fontsize=50)

\# formatting the colorbar

cbar_ax.set_yticklabels(range(0,3026,500),fontsize=30)

cbar_ax.set_ylabel('Time Taken to Fix A2B2',fontsize=35)

figtime.savefig(f'Time D={D}.tiff', dpi=300)

**Code 7 ‘X_Drf.py’: This code was used to know how initial linkage disequilibrium (D) and female recombination rate (rf) affected the system with different kinds of modifiers, when the loci were present on X chromosomes. This generated the figures S7 and S8.**

\# import libraries

import numpy as np

import matplotlib.pyplot as plt

import seaborn as sns

from random import uniform

RF=np.arange(0,0.6,0.1)

Rf=[round(x,2) for x in RF] # range of female recombination rates

h=0.2 # setting value of dominance coefficient

sf=0.2 # setting value of female selection coefficient

sm=0.2 # setting value of male selection coefficient

Checklist=np.arange(1-0.020,1+0.020,0.001)

checklist=[round(i,3) for i in Checklist] # a checklist to cross check whether the frequencies are summing upto approximately 1 at every time step

T=3000 # number of generations

Krange=np.arange(0,1.01,0.1)

krange=[round(k,2) for k in Krange] # range of values for k1 and k2

\# determining initial frequencies of various alleles

A1f=max(0.01,min(0.99,(2*sf*(1-h)-sm)/(2*sf*(1-2*h))))

A2f=1- A1f

A2m=max(0.01,min(0.99,(sm-2*sf*h)/(2*sf*(1-2*h))))

A1m=1-A2m

B1=0.95

B2=0.05

\# determining initial frequencies of various haplotypes

x1=A1f*B1

x2=A1f*B2

x3=A2f*B1

x4=A2f*B2

y1=A1m*B1

y2=A1m*B2

y3=A2m*B1

y4=A2m*B2

\# determining possible values of D

possible_D=[1-x1,x1-0,1-x2,x2-0,1-x3,x3-0,1-x4,x4-0,1-y1,y1-0,1-y2,y2-0,1-y3,y3-0,1-y4,y4-0]

DRange=np.arange(-min(possible_D),min(possible_D),0.01)

Drange=[round(x,2) for x in DRange]

print(Drange)

data_time=[[[[T+25 for rf in Rf] for D in Drange] for k2 in krange] for k1 in krange] # array storing fixation time data

data_freq=[[[[] for D in Drange] for k2 in krange] for k1 in krange] # array storing frequency of A2B2 at 3000 generation

for k1ind, k1 in enumerate(krange): # iterating over values of k1 and k2

for k2ind, k2 in enumerate(krange):

for Dind, D in enumerate(Drange): # iterating over values of initial linkage disequilibrium

for rfind, rf in enumerate(Rf): # iterating over values of female recombination rates

\# setting initial frequencies of each haplotype along with initial linkage disequilibrium

x1=A1f*B1

x2=A1f*B2

x3=A2f*B1

x4=A2f*B2

y1=A1m*B1

y2=A1m*B2

y3=A2m*B1

y4=A2m*B2

x1+=D

x2+=-D

x3+=-D

x4+=D

y1+=D

y2+=-D

y3+=-D

y4+=D

#Storing frequencies

X1=[x1]

X2=[x2]

X3=[x3]

X4=[x4]

Y1=[y1]

Y2=[y2]

Y3=[y3]

Y4=[y4]

#fitness scheme

f11= 1

f13= 1-k1*sf

f31= 1-sf

f33= 1-(1-k2)*sf

f12= 1-k1*h*sf

f32= 1 - (1+h - h*(1-k2)-k2)*sf

f21= 1- h*sf

f23= 1 - abs((1-k1-k2)*h*sf)

f22r= 1- abs(h*sf*(1-k1))

f22c= 1-(1-k2)*sf*h

m1=1-sm

m2=1-sm

m3=1

m4=1

#time series

A2B2fixed = False # a variable to keep check of whether the haplotype A2B2 is fixed

for t in range(T):

\# computing haplotype frequencies at (t+1)th generation

wm=x1*m1+x2*m2+x3*m3+x4*m4

wf=x1*y1*f11 + (x1*y2+x2*y1)*f12 + x2*y2*f13 + (x1*y3+x3*y1)*f21 + (x2*y3+x3*y2)*f22r + (x1*y4+x4*y1)*f22c + (x2*y4+x4*y2)*f23 + x3*y3*f31 + (x3*y4+x4*y3)*f32 + x4*y4*f33

Y1+=[x1*m1/wm]

Y2+=[x2*m2/wm]

Y3+=[x3*m3/wm]

Y4+=[x4*m4/wm]

X1+=[(2*x1*y1*f11 + (x1*y2+x2*y1)*f12 + (x1*y3+x3*y1)*f21 + (1-rf)*f22c*(x1*y4+x4*y1) + rf*f22r*(x2*y3+x3*y2))/(2*wf)]

X2+=[(2*x2*y2*f13 + (x1*y2+x2*y1)*f12 + (x2*y4+x4*y2)*f23 + (1-rf)*f22r*(x3*y2+x2*y3) + rf*f22c*(x4*y1+x1*y4))/(2*wf)]

X3+=[(2*x3*y3*f31 + (x1*y3+x3*y1)*f21 + (x3*y4+x4*y3)*f32 + (1-rf)*f22r*(x2*y3+x3*y2) + rf*f22c*(x4*y1+x1*y4))/(2*wf)]

X4+=[(2*x4*y4*f33 + (x2*y4+x4*y2)*f23 + (x4*y3+x3*y4)*f32 + (1-rf)*f22c*(x4*y1+x1*y4) + rf*f22r*(x2*y3+x3*y2))/(2*wf)]

x1=X1[-1]

x2=X2[-1]

x3=X3[-1]

x4=X4[-1]

y1=Y1[-1]

y2=Y2[-1]

y3=Y3[-1]

y4=Y4[-1]

\# checking whether A2B2 has fixed in the population

if round(x4+y4,3)==2 and A2B2fixed==False:

data_time[k1ind][k2ind][Dind][rfind]=t # adding time of fixation to array

A2B2fixed=True

\# adding the frequency of A2B2 at 3000th generation to array

if t==T-1:

data_freq[k1ind][k2ind][Dind]+=[round((x4+y4)/2,3)]

\# checking for haplotype frequencies summing upto 1

if round(x1+x2+x3+x4,3) not in checklist:

print(f'x not summing upto 1-{x1+x2+x3+x4}',x1,x2,x3,x4,t)

if round(y1+y2+y3+y4,3) not in checklist:

print(f'y not summing upto 1-{y1+y2+y3+y4}',y1,y2,y3,y4,t)

figfreq,axfreq=plt.subplots(len(krange),len(krange),sharey=True,sharex=True,figsize=(40,40)) #setting figure parameters

cbar_ax = figfreq.add_axes([.91, .3, .03, .4]) # setting the colorbar axis

for i,k1 in enumerate(data_freq): # iterating over frequency data

for j,map in enumerate(k1): # iterating over frequency data

g=sns.heatmap(map,vmin=0,vmax=1,cmap='YlGn',cbar=True,cbar_ax=cbar_ax, cbar_kws={'shrink': 1.8,"use_gridspec":False},ax=axfreq[i,j]) # plotting the heatmap

\# formatting the heatmap

axfreq[i,j].set_yticklabels(Drange,fontsize=25)

axfreq[i,j].set_xticklabels(Rf,fontsize=25)

\# setting subplot titles and labelling rows (on the left) and columns (bottom)

if i==len(data_freq)-1:

axfreq[i,j].set_xlabel(f'rf \n k2 = {krange[j]}', fontsize=35)

if j==0:

axfreq[i,j].set_ylabel(f'k1 = {krange[i]} \n D', fontsize=35)

#axfreq[0,0].set_title(f'D = {D}',fontsize=50)

\# customizing colorbar

cbar_ax.set_yticklabels([round(x,2) for x in np.arange(0,1.1,0.2)],fontsize=30)

cbar_ax.set_ylabel(f'Frequency of A2B2 at generation {T}',fontsize=35)

figfreq.savefig(f'X-linked Drf Freq at 3000 generation.tiff',dpi = 300) # save figure

figtime,axtime=plt.subplots(len(krange),len(krange),sharey=True,sharex=True,figsize=(40,40),gridspec_kw={'hspace':0.25,'wspace':0.125}) # setting figure parameters)

cbar_ax = figtime.add_axes([.91, .3, .03, .4]) # setting new colorbar axis

for i,k1 in enumerate(data_time): # iterating over time data

for j,map in enumerate(k1): # iterating over time data

g=sns.heatmap(map,vmin=0,vmax=T+25,yticklabels=Drange,xticklabels=Rf,cmap='GnBu',cbar=True,cbar_ax=cbar_ax, cbar_kws={'shrink': 1.8,"use_gridspec":False},ax=axtime[i,j]) # plotting the heatmap

\# formatting the heatmap

axtime[i,j].set_yticklabels(Drange,fontsize=25)

axtime[i,j].set_xticklabels(Rf,fontsize=25)

\# setting subplot titles and labelling rows (on the left) and columns (bottom)

if i==len(data_time)-1:

axtime[i,j].set_xlabel(f'rf \n k2 = {krange[j]}',fontsize=35)

if j==0:

axtime[i,j].set_ylabel(f'k1 = {krange[i]} \n D', fontsize=35)

\# formatting the colorbar

cbar_ax.set_yticklabels(range(0,3026,500),fontsize=30)

cbar_ax.set_ylabel('Time Taken to Fix A2B2',fontsize=35)

figtime.savefig(f'X-linked Drf Time.tiff', dpi=300)
